# A network of Notch-dependent and -independent *her* genes controls neural stem and progenitor cells in the zebrafish thalamic proliferation zone

**DOI:** 10.1101/2022.09.15.508073

**Authors:** Christian E. Sigloch, Dominik Spitz, Wolfgang Driever

**Affiliations:** Developmental Biology, Institute Biology 1, Faculty of Biology, University of Freiburg, Hauptstrasse 1, 79104 Freiburg, Germany; Spemann Graduate School of Biology and Medicine SGBM, University of Freiburg, Albertstraße 19A, 79104 Freiburg, Germany; Signalling Research Centres CIBSS and BIOSS, University of Freiburg, Schänzlestraße 18, 79104 Freiburg, Germany; Renal Division, Department of Medicine, Faculty of Medicine and Medical Center, University of Freiburg, 79106 Freiburg, Germany

**Keywords:** neurogenesis, neural stem cells, HES/HER genes, Delta-Notch signaling, neural proliferation zone, thalamus, zebrafish

## Abstract

Neural proliferation zones mediate brain growth, and employ Delta/Notch signaling and HES/HER transcription factors to balance neural stem cell (NSC) maintenance and generation of progenitors and neurons. We investigated Notch-dependency and function of *her* genes in the thalamic proliferation zone of developing zebrafish larvae. Nine Notch-dependent genes, *her2, her4.1-5, her12, her15.1-2*, and two Notch-independent genes, *her6, her9*, are differentially expressed, and define distinct NSC and progenitor populations. *her6* prominently executes patterning information to maintain NSCs and the zona limitans intrathalamica Shh signaling activity. *her6, her9* double mutants reveal that Notch-independent *her* genes predominantly regulate NSC maintenance and transition into the progenitor pool. Surprisingly, combined deletion of all Notch-dependent *her* genes does not affect NSCs or progenitor formation. Combined genetic manipulation of up to eleven Notch-dependent and -independent *her* genes revealed that Notch-dependent *her* genes may regulate progenitor progression into neurogenesis, but not progenitor generation itself. The *her* gene network is partially redundant, with Notch-independent *her* genes better substituting for loss of Notch-dependent genes than vice versa. Together, *her* gene regulatory feedback loops and crossregulation contribute to the observed robustness of NSC maintenance.

## Introduction

The zebrafish brain continually grows and adds new neurons throughout development and into adult stages, and has high capacity to regenerate neurons after injury (Kishimoto et al., 2012; Zupanc, 2001). The source of these new neurons are neural stem cells (NSCs) located in at least 16 adult neural proliferation zones (NPZs) (Grandel et al., 2006; Zupanc et al., 2005). These anatomically defined NPZs are mostly located at the ventricular wall (Adolf et al., 2006), where Hairy and Enhancer of split (HES/Her) transcription factors are expressed and regulate neurogenesis (Chapouton et al., 2011). During development, the embryonic neuroepithelium establishes highly active late embryonic and early larval proliferation zones, which develop into neural stem cell niche-like adult proliferation zones. So far, the mechanisms that balance NSC maintenance and neurogenesis in these larval NPZs are not well understood.

HES/Her factors in NPZs control neurogenesis by inhibiting proneural gene mediated differentiation (Ishibashi et al., 1994; Kageyama et al., 2007; Sasai et al., 1992). In mice, absence of the basic Helix loop Helix (bHLH) transcription factors HES1 and HES5 leads to accelerated differentiation, accompanied by loss of radial glia and later-born cell types (Hatakeyama et al., 2004). Moreover, in zebrafish and mouse, *Hes/her* genes were shown to be important for the maintenance of local stem cell populations at boundaries that form signaling and growth centers (Baek et al., 2006; Hatakeyama et al., 2004), for example the midbrain-hindbrain boundary (Geling et al., 2004) and the zona limitans intrathalamica (ZLI) (Scholpp et al., 2009).

In *Drosophila*, the expression of *E(spl)* is Notch-dependent (Jennings et al., 1994), while the expression of *hairy*, encoding a closely related bHLH factor, is directed by local patterning signals (Riddihough and Ish-Horowicz, 1991). However, this strict distinction is not conserved in vertebrates (Chapouton et al., 2011; Davis and Turner, 2001). In mouse, *Hes1* and *Hes5* are downstream targets of Notch signaling, although *Hes1* expression and stability are the result of additional Notch-independent input (Kageyama et al., 2007; Ohtsuka et al., 1999). In contrast to HES1 and HES5, the transcription factor HES6 functions as a negative regulator of HES1, thereby promoting differentiation (Bae et al., 2000), and its expression is controlled by proneural genes (Koyano-Nakagawa et al., 2000).

In zebrafish, the *her* gene family expressed in NPZs is significantly expanded compared to mammals (Zhou et al., 2012). Nine *Hes5* zebrafish homologs are located in two gene clusters, *her4.1* to *her4.5* and *her12* on chromosome 23, and *her15.1, her15.2* and *her2* on chromosome 11. There are two *Hes1* homologs, *her6* and *her9*, and four *Hes6* homologs, *hes6, her8a, her8.2* and *her13. her4* and *her15, her6* and *her9*, as well as *her8a* expression have been characterized in adult neural stem cell (NSC) zones (Chapouton et al., 2011; Zhou et al., 2012). Zebrafish homologs of *Hes1*, *Hes5* and *Hes6* have also been investigated in the neural plate during early embryonic development (Bae et al., 2005; Takke et al., 1999; Webb et al., 2011). So far however, little is known about potential cross-regulatory interactions or redundancies among *her* genes in NSCs and during neurogenesis in larval neural proliferation zones.

Here, we analyzed the impact of Notch signaling and *her* family genes on expression of specific *her* genes in NSCs and neural progenitor cells (NPCs) to reveal the organization of *her* gene regulatory networks in neurogenesis. We focus on the highly active early larval thalamic proliferation zone (TPZ) at the ventricular wall of the diencephalon in the thalamus proper (dorsal thalamus), the ZLI, and the prethalamus (Mueller, 2012; Scholpp and Lumsden, 2010), together constituting the thalamic complex. Based on *her* gene expression, we define distinct NSC populations in the TPZ. We generated compound mutants for two Notch-independent (*her6* and *her9)* and nine Notch-dependent *(her2, her4.1-5, her12, her15.1-2*) *her* genes. Based on loss-of-function and overexpression experiments, we generated a model for cross-regulatory interactions of Notch-dependent and -independent *her* genes in the TPZ. Our data indicate that Notch-independent *her* genes are important for NSC maintenance and patterning in the TPZ. Notch-dependent *her* genes however appear not strictly required for NSC maintenance but might rather modulate progression of neurogenesis.

## Results

### *her* family gene expression in the thalamic proliferation zones

We surveyed published information (www.zfin.org; Chapouton et al., 2011; Shankaran et al., 2007; Sieger et al., 2004; Zhou et al., 2012) on zebrafish *her* family genes for expression in the diencephalon, and selected homologs of mammalian *Hes1 (her6, her9), Hes5 (her2, her4.1-4.5, her 12, her15.1-15.2)* and *Hes6 (her8a, her8.2)* for analysis of their expression patterns in the TPZ by whole mount in situ hybridization (WISH; Fig. 1A-L; Fig. S1A-H). The five *her4.1-4.5* genes together with *her12* form a cluster on chromosome 23 (“*her4;12 cluster”)*, while the two *her15.1-15.2* genes form a cluster together with *her2* on chromosome 11 (“*her2;15 cluster”).* Genes within each cluster have high sequence similarity. Our *her4* probe was designed to detect expression of all five *her4* genes *(her4.1-her4.5)* combined (subsequently referred to as *her4).* Similarly, our *her15* probe detects *her15.1* and *her15.2* (referred to as *her15).* We focused our analysis on two days (48 hours post fertilization, hpf) to four days (96 hpf) old embryos and early larvae, when the TPZ is a highly active proliferation zone with a diversity of neurogenesis mechanisms involving *neurog1, ascl1* and *olig2* (Scholpp et al., 2009; Virolainen et al., 2012), (see also Fig. 7C and Fig. 11A, B).

**Fig. 1.**
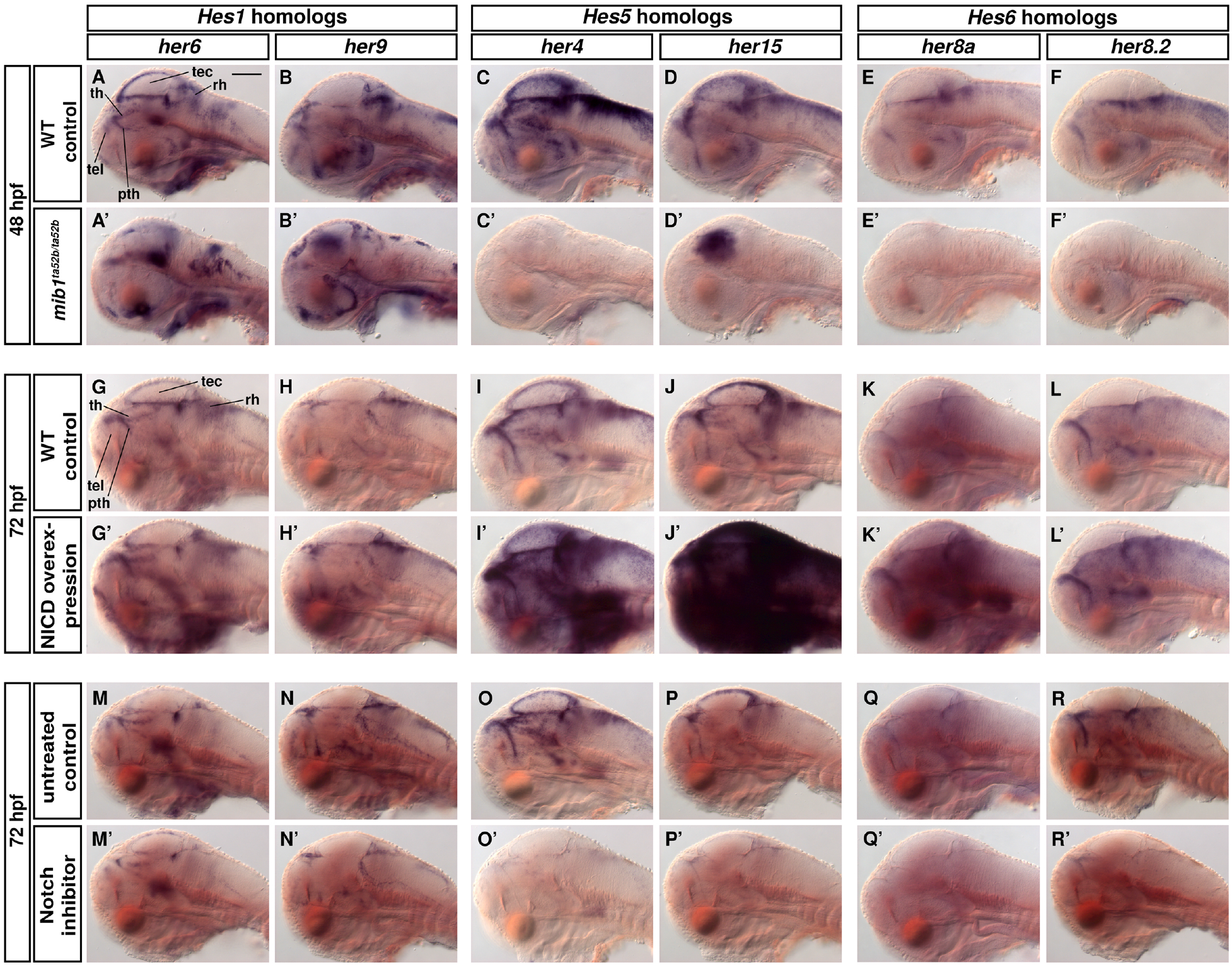
Notch signaling dependent and independent *her* gene expression. **(A-R’)** *her6, her9, her4 (her4.1-her4.5* cluster probe)*, her15 (her15.1* and *her15.2* cluster probe)*, her8a* and *her8.2* expression visualized by whole mount in situ hybridization (WISH). **(A-F)** 48 hpf WT sibling controls *(mib1^+/+^* or *mib1^+/ta52b^)* and **(A’-F’)** *mind bomb* mutant embryos *(mib1^ta52b^)* which lack Notch activity. **(G-L’)** Overactivated Notch signaling by heat-shock induced overexpression of the Notch intracellular domain (NICD) compared with heat shocked WT siblings at 72 hpf. **(M-R’)** Notch inhibition by LY-411575 in 72 hpf larvae in comparison to DMSO controls. Larvae were treated from 64 to 72 hpf with 10 μM LY-411575 or 2% DMSO (controls), and fixed at 72 hpf. Sagittal optical sections at the midline are shown (lateral views, single DIC image planes, anterior at left, dorsal at top). Three larvae were imaged per condition and one representative image was selected. Scale bar, 100 μm. Abbreviations: pth, prethalamus; rh, rhombencephalon; tec, tectum; tel, telencephalon; th, thalamus proper. An anatomical reference scheme is provided in Fig. 2E.

In the TPZ, *her6*, *her9*, *her4, her2*, and *her15* expression were detected with strong WISH signals. *her2* is clustered with *her15.1-15.2*, and has a similar expression domain. The expression of these *her* genes in the thalamic complex was mostly restricted to the wall of the diencephalic ventricle and the adjacent lateral ventricle (Fig. S1A’-H’).

In contrast, the *Hes6* homologs *her8a* and *her8.2* had weak WISH signals in the TPZ (Fig. S1C-D). *her8.2* and *her8a* are broadly expressed in neural proliferations zones at 48 hpf, but expression decreases on the fourth day of development, such that at 96 hpf we could detect only low levels of *her8.2* and very faint *her8a* expression in the TPZ (Fig. 1E,F,K,L and Fig. S1C-D). *her12* expression appears restricted to the hindbrain and later the midbrain-hindbrain boundary (Fig. S1E,E’; Fig. S2A,G), and thus, despite being in a cluster with *her4* genes, differs significantly in expression.

In summary, while expression of the *Hes6* homologs *her8a* and *her8.2* decreases during early TPZ formation, *Hes1 (her6, her9)* and *Hes5 (her2, her4.1-4.5, her15.1-15.2)* homologs likely provide the major *her* gene activity in the maturing TPZ.

### Notch dependency of neural *her* gene expression

Regulation of zebrafish *her* gene expression by Notch signaling has been characterized during embryonic stages (Pasini et al., 2004; Shankaran et al., 2007; Takke et al., 1999), but not in larval NPZs. To assess Notch dependency of expression of *Hes1, Hes5* and *Hes6* homologs, we used three approaches. We first analyzed *mind bomb1 (mib1^ta52b^)* mutants that lack Notch signaling due to impaired internalization of the ligand Delta (Itoh et al., 2003). Second, we globally overexpressed the Notch Intracellular Domain (NICD) by heat shock in 72 hpf larvae (Scheer and Campos-Ortega, 1999). Third, we treated larvae with the Notch inhibitor LY-411575 (Rothenaigner et al., 2011).

The *Hes1* homologs *her6* and *her9* are still expressed at largely normal levels in *mib1^ta52b^* mutant embryos (Fig. 1A-B’), and NICD overexpression only slightly increased *her9* and hardly at all *her6* expression (Fig. 1G-H’). Upon Notch inhibitor treatment, *her6* and *her9* expression continued, albeit at slightly reduced levels (Fig. 1M-N’). These experiments demonstrate *her6* and *her9* expression to be Notch-independent at analyzed larval stages.

In contrast, the expression of *Hes5 (her2, her4, her12* and *her15)* and *Hes6 (her8a* and *her8.2)* homologs is lost in *mib1^ta52b^* mutants (Fig. 1C’-F’, Fig. S2A-D’). The ectopic *her15* expression in the tectum of *mib1* mutants appeared very different from the expression in wild type (WT; Fig. 1D,D’), and may reflect dysregulation. Expression of *Hes5* homologs was strongly induced by NICD, while the *Hes6* homologs showed only a moderate increase in expression (Fig. 1I-L’). Upon Notch inhibitor treatment, expression of *Hes5* and *Hes6* homologs was nearly absent (Fig. 1O-R’; Fig. S2G-I’), except *her2*, which was expressed at reduced levels after LY-411575 treatment (Fig. S2J,J’). These findings suggest that, while *her2* and *her15* are located in the same cluster, *her15* is mostly Notch-dependent while *her2* may be only partly controlled by Notch.

To conclude, in the TPZ of 3-4 dpf larvae, expression of the *Hes1* homologs *her6* and *her9* are only weakly affected by perturbation of Notch signaling and are hereafter referred to as Notch-independent. In contrast, *her4* and *her15* expression are strictly dependent on Notch signaling. Given that *her8a* and *her8.2* have only low and declining levels of expression in the TPZ on the fourth day of development, we did not include these two genes in our further analysis.

### Differential expression of *her* genes in the thalamus proper, prethalamus and ZLI

We next investigated by fluorescent WISH whether Notch-dependent and Notch-independent *her* genes are differentially expressed in the TPZ (Fig. 2). *her4* and *her6* expression was mostly restricted to the periventricular TPZ wall, and largely co-expressed with the NSC marker *sox2* (Fig. S3A-B”). During early development, *her6* is expressed in the presumptive thalamic complex (Scholpp et al., 2009). During larval development, *her6* expression is absent from the ZLI, but maintained in the rostral thalamus and prethalamus (Fig. 2A,A’,F; Scholpp and Lumsden, 2010). At the ventricle along the midline, *her4* is strongly expressed where *her6* is absent, namely in the ZLI, caudal thalamus (cTh) and the pretectum (Fig. 2A). In the caudal thalamus, *her4* is also expressed laterally adjacent to the ventricular wall (Fig. 2A’). *her4* and *her15* were mostly co-expressed in the ZLI, caudal thalamus and pretectum (Fig. 2B,B’; for comparison, chromogenic *her4* WISH in Fig. 2G). *her9* was expressed only weakly in the ZLI and the cTh, and both *her6* and *her9* were expressed in rather non-overlapping territories (Fig. 2C,C’). We also directly compared the expression of both Notch-dependent *her4* and *her15* with Notch-independent *her6* and *her9* combined, using pairwise mixed probes, and found that their expression domains together covered the whole ventricular zone of the thalamic complex (Fig. 2D-D”’).

**Fig. 2.**
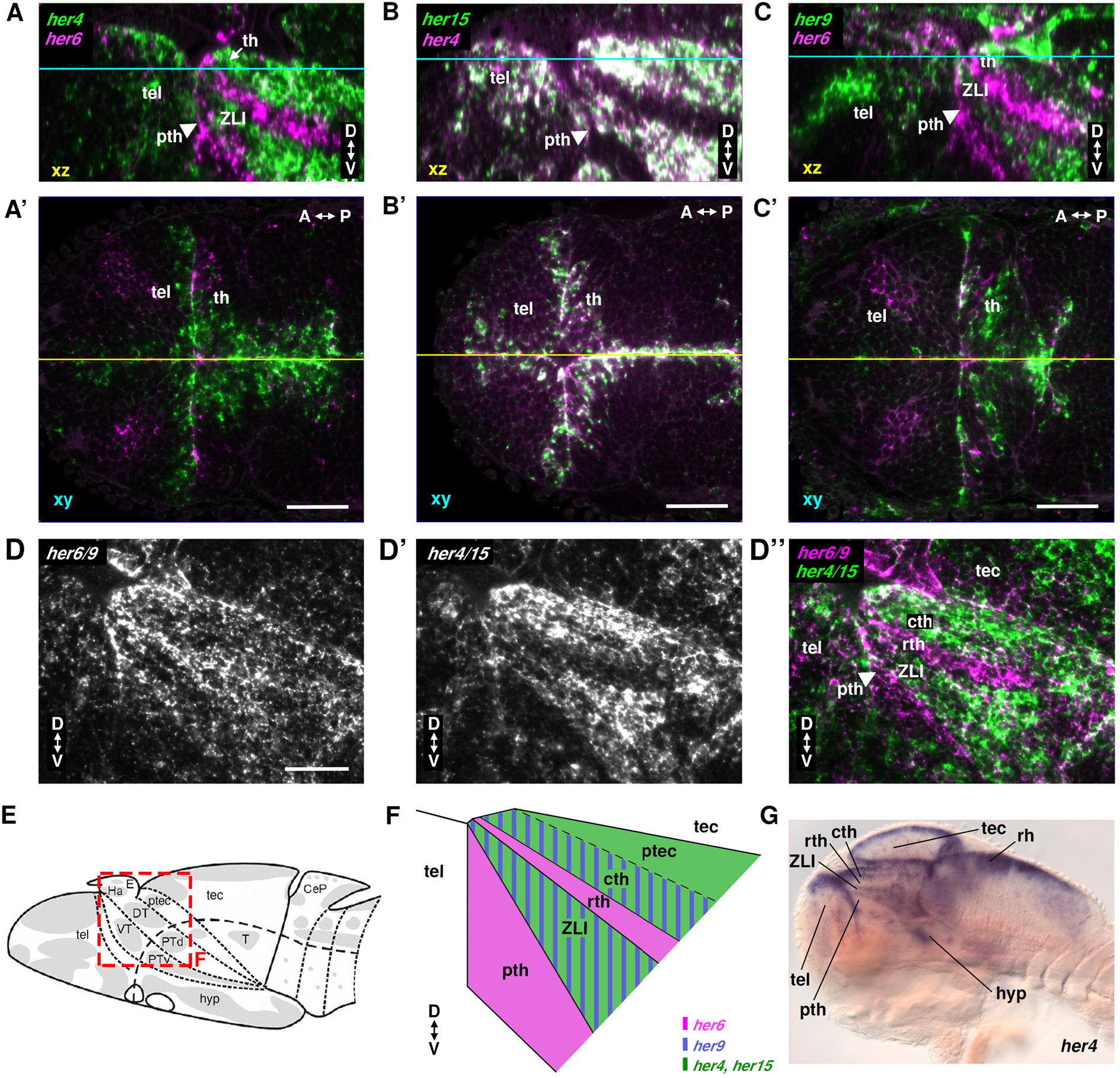
Co-expression analysis of *her* genes in the thalamus of 3 dpf larvae. **(A-D’’)** Confocal imaging of double-fluorescent WISHs at 3 dpf. **(A, A’)** Expression of *her4* (green) and *her6* (magenta). **(B, B’)** Expression of *her15* (green) and *her4* (magenta). **(C, C’)** Expression of *her9* (green) and *her6* (magenta). A, B and C are lateral views of midline sagittal orthogonal reconstructions of dorsal view (horizontal) stacks (levels indicated by yellow line in A’, B’ and C’ respectively). A’, B’ and C’ are dorsal views with the optical horizontal section level indicated in cyan in A, B and C. **(D-D’’)** Lateral views of combined *her6/her9* (D’’ magenta and D white, *her6* and *her9* probes mixed) and combined *her4/her15* (D’’ green and D’ white, *her4* and *her15* probes mixed) detection. D’’ compares *her6/her9* to *her4/her15* expression. **(E)** Scheme of a 3 dpf zebrafish larval brain modified from (Mueller and Wullimann, 2016). Proliferation zones are indicated in grey. The dashed line separates the alar from basal plate. **(F)** Schematic map shows distinct domains of *her* gene expression at the ventricular wall of the alar diencephalon. Striped areas indicate that genes are co-expressed or expressing cells intermingled. **(G)** *her4* WISH in a 3 dpf larva with anatomical annotation, sagittal optical section. **(A-G)** The *her4* probe detects *her4.1-her4.5* and the *her15* probe detects *her15.1* and *her15.2.* All scale bars, 50 μm. Abbreviations: A-P indicates anterior-posterior orientation, D-V dorsal-ventral orientation. CeP, cerebellar plate; DT, dorsal thalamus proper, subdivided into rostral thalamus (rth) and caudal thalamus (cth); E, epiphysis; Ha, habenula; hyp hypothalamus; ptec, pretectum; PTd, dorsal part of posterior tuberculum; pth, prethalamus; PTv, ventral part of posterior tuberculum; T, midbrain tegmentum; tel, telencephalon; tec, tectum opticum; VT, ventral thalamus or prethalamus (pth); ZLI, zona limitans intrathalamica.

Looking at combined expression domains, *her4, her15*, and, to a lesser extent, *her9* are co-expressed in the ZLI and the caudal thalamus, while *her6* is selectively expressed in prethalamus and rostral thalamus (Fig. 2F). Thus, interestingly, Notch-independent *her9* is expressed in the same anatomical area as the Notch-dependent *her4* and *her15*, while the second Notch-independent gene *her6* is expressed in largely exclusive territories devoid of other *her* gene expression.

When we analyzed *her4, her6*, and *her9* by triple fluorescent HCR at cellular resolution in the thalamus (Fig. S4), we realized that domains of high expression of each gene at the ventricle are mostly exclusive. In addition, in cells which may correlate to neural progenitors, we detected *her4* expression at lower level distant from the high *sox*2 expressing ventricular NSCs. To validate this hypothesis, we used triple fluorescent HCR to analyze *sox2, ascl1b*, and *neurog1* expression in the same anatomical region (Fig. S5; Movie M1). The data show high *sox2* expressing cells (Sox2^high^) at the ventricular walls largely devoid of *ascl1b* and *neurog1* expression, corresponding to NSCs. In addition, we find a *sox2* low expressing domain (Sox2^low^) in the region corresponding to low *her4* expression, which also expresses *ascl1b* or *neurog1*, and thus are neural progenitor cells (NPCs) in early neurogenesis.

Next, we analyzed *her4, her6* and *her9* in relation to proneural gene expression in the Sox2^low^ NPC region (Fig. S6). We observe that in this domain, *neurog1* expression largely overlaps with *her4* expression, but not with *her6* and *her9.* Similarly, expression of *ascl1a/b* is largely exclusive from the *her6* expressing regions (Fig. S6D). *ascl1a/b* expression overlaps widely with *dla* expression in the Sox2^low^ *her4* region (Fig. S6E), suggesting that Delta and Notch-dependent *her* genes control *neurog1* and *ascl1a/b* in the NPC region. To investigate in more detail the relation of Delta signaling and *her* genes in NSC and NPC regions, we analyzed combined expression of *dla* with *sox2, her4, her6* and *her9* (Fig. S7). Surprisingly, *dla* expression appears higher in the NPC Sox2^low^ region than in the Sox2^high^ ventricular zone (Fig. S7A). *her4* and *dla* expression largely overlap in the NPC region (Fig. S7B). In contrast, regions with highest levels of *her6* and *her9* expression are devoid of *dla* expression (Fig. S7C-D).

In conclusion, our data identify at least three fundamentally distinct *her* expressing cell populations in the TPZ (Fig. 11A,B): (1) Sox2^high^ NSCtype1 at the ventricular wall with predominant expression of Notch-independent *her6*, and absent *dla*; (2) Sox2^high^ NSCtype2 at the ventricular wall with high expression of Notch-dependent *her4* and/or *her15*, low *her9* expression, and in most regions *dla* expression. However, *her6* expression is absent from these regions; (3) Sox2^low^ NPCs away from the ventricular wall with *dla* expression, either *ascl1b* or *neurog1* expression, and low-level expression of Notch-dependent *her4.*

### Auto- and cross-regulation of Notch-independent *her* genes

Given the exclusive expression domains in the TPZ, we tested genetically whether *her6* and *her9* might cross-repress each other. We used CRISPR/Cas9 to delete large parts of the bHLH domains of *her6* and *her9*, and isolated mutant alleles with premature stop codons that leave the remaining aminoterminal peptides non-functional (Fig. 3A, B). We analyzed genetic interactions in *her6* and *her9* single and double mutants (Fig. 3C-N) using WISH probes that detect the shortened mutant transcripts. At 48 hpf, *her6* expression levels in several regions appear enhanced in *her6* mutants, and even stronger in *her6, her9* double mutants, while *her6* expression in *her9* mutants appears similar to WT in most brain regions (Fig. 3C-F’). Upregulation of *her6* in double mutants is even stronger at 96 hpf (Fig. 3G-H’). *her9* expression levels in *her6* and *her9* single and double mutants at 48 hpf appear not strongly affected, while *her9* expression at 96 hpf is slightly increased in double mutants (Fig. 3I-N’). To quantify potential cross-regulations independently, we also analyzed *her6* and *her9* mRNA levels by qPCR on genotyped 4 dpf embryonic heads. We found a significant upregulation of *her6* transcripts in *her6* mutants and of *her9* transcripts in *her9* mutants (Fig. 3O; p-values see Supplementary Table 5), indicating negative autoregulation. We next asked whether *her6* and *her9* cross-regulation is redundant, and analyzed *her6*, *her9* double mutants. Upregulation of *her6* and *her9* transcripts in *her6*, *her9* double mutant was significantly stronger than in single mutants (Fig. 3O). We conclude that *her6* and *her9* exert negative cross-regulation on each other, partially redundant with negative autoregulation. In fact, as revealed from lower *her6* expression in *her9* than in *her6* mutants, and from lower *her9* expression in *her6* than in *her9* mutants, negative autoregulation appears stronger than cross-regulation. Nevertheless, the strong upregulation of *her6* and *her9* in double mutants reveals a significant contribution of negative cross-regulation.

**Fig. 3.**
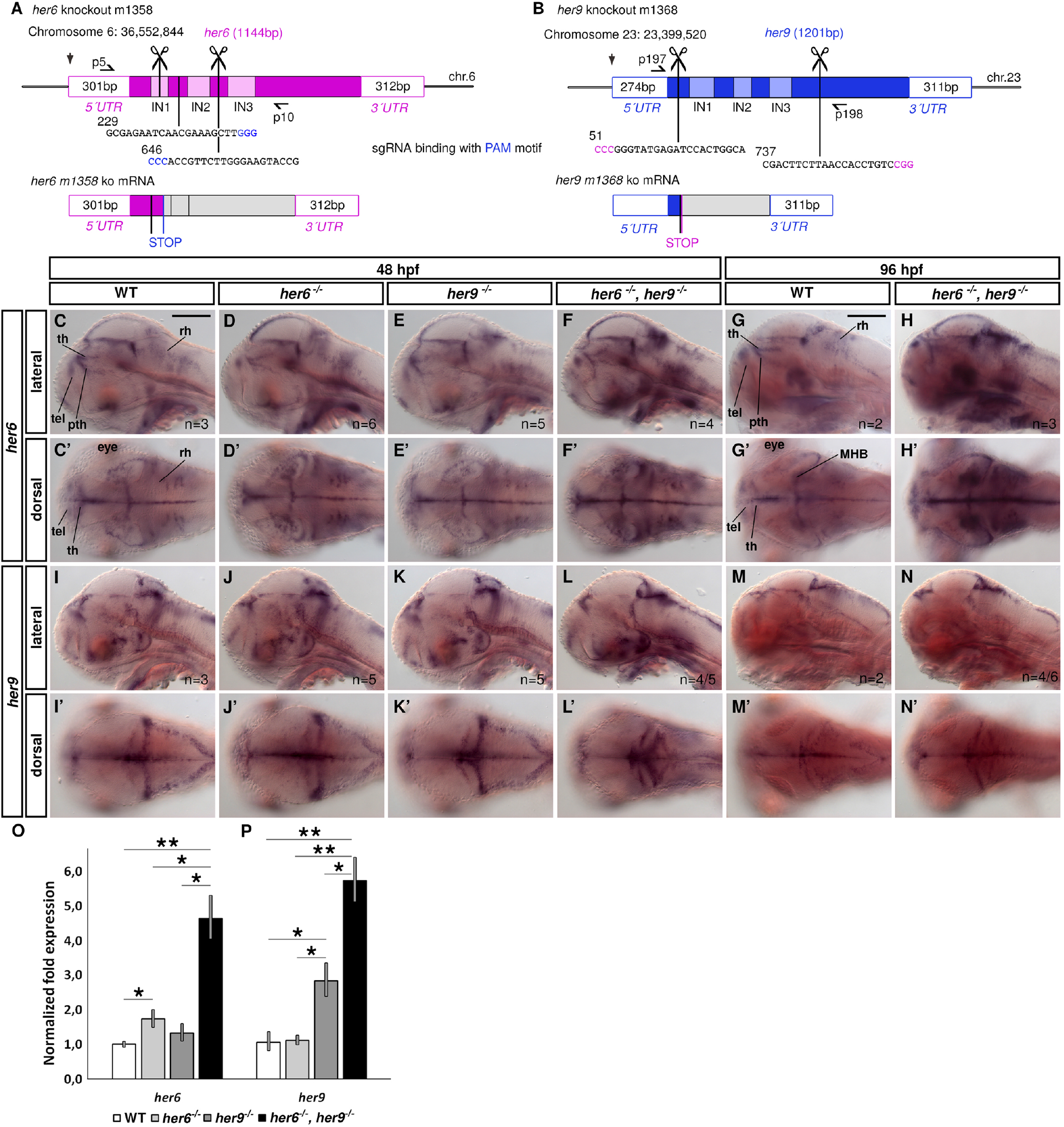
*her6* and *her9* single and double mutants reveal auto- and cross-regulation. **(A, B)** Schematic for CRISPR/Cas9 deletion of the bHLH domains within the *her6* and *her9* genes, respectively. Straight vertical lines show the sgRNA binding sites. Scissors indicate the endpoints of the small deletion after non-homologous end joining. Half-arrows indicate primers (with names) which were used in genotyping PCRs. **(C-N’)** WISH for *her6* **(C-H’)** or *her9* **(I-N’)** in WT (WT), *her6* and *her9* single or double mutants at indicated stages. (C-N) show lateral single sagittal optical plane images, (C’-N’) dorsal views. Numbers of embryos analyzed are indicated in lower right corner. If different phenotypes were observed, n/n indicates numbers of embryos with phenotype shown in panel / total analyzed. Scale bars, 200 μm. **(O, P)** qPCR analysis of *her6* or *her9* expression in *her6* and *her9* single and double mutants. Error bars, SEM. Bars with asterisks indicate significant differences (* p< 0.05; ** p< 0.01; for exact values see Supplementary Table 5). Abbreviations: MHB, midbrain-hindbrain boundary; pth, prethalamus; rh, rhombencephalon; tel, telencephalon; th, thalamus proper.

### Combined loss of *her6* and *her9* activity affects neural stem cells, progenitor cells and ZLI

We further analyzed *her6*, *her9* double mutants to determine the impact of loss of Notch-independent *her* gene function on NSCs, progenitors, and cell proliferation in the TPZ. We used an anti-Sox2 antibody to mark NSCs by whole mount immunofluorescence (Fig. 4A-D” and Movie M2). In WT larvae, Sox2 positive nuclei lined the thalamic ventricular walls, including the lateral ventricle away from the midline, and nuclei with lower Sox2 immunoreactivity (Sox2^low^) were also detected in subapical positions adjacent to the cell layer that forms the ventricular wall (Fig. 4A’ and cutout in A_1_). In the *her6, her9* double mutants, the subapical Sox2^low^ cell nuclei were strongly reduced or missing, and Sox2^high^ cell nuclei restricted to the ventricular wall (Fig. 4B’ and Bi; *her9* and *her6* single mutants each have largely normal distribution of Sox2 positive nuclei, data not shown). In addition, the lateral extension of Sox2^high^ nuclei along the lateral expansion of the ventricle was strongly reduced, indicating a loss of ventricular NSCs (Fig. 4A” and B”, white arrowheads and Movie M2).

**Fig. 4.**
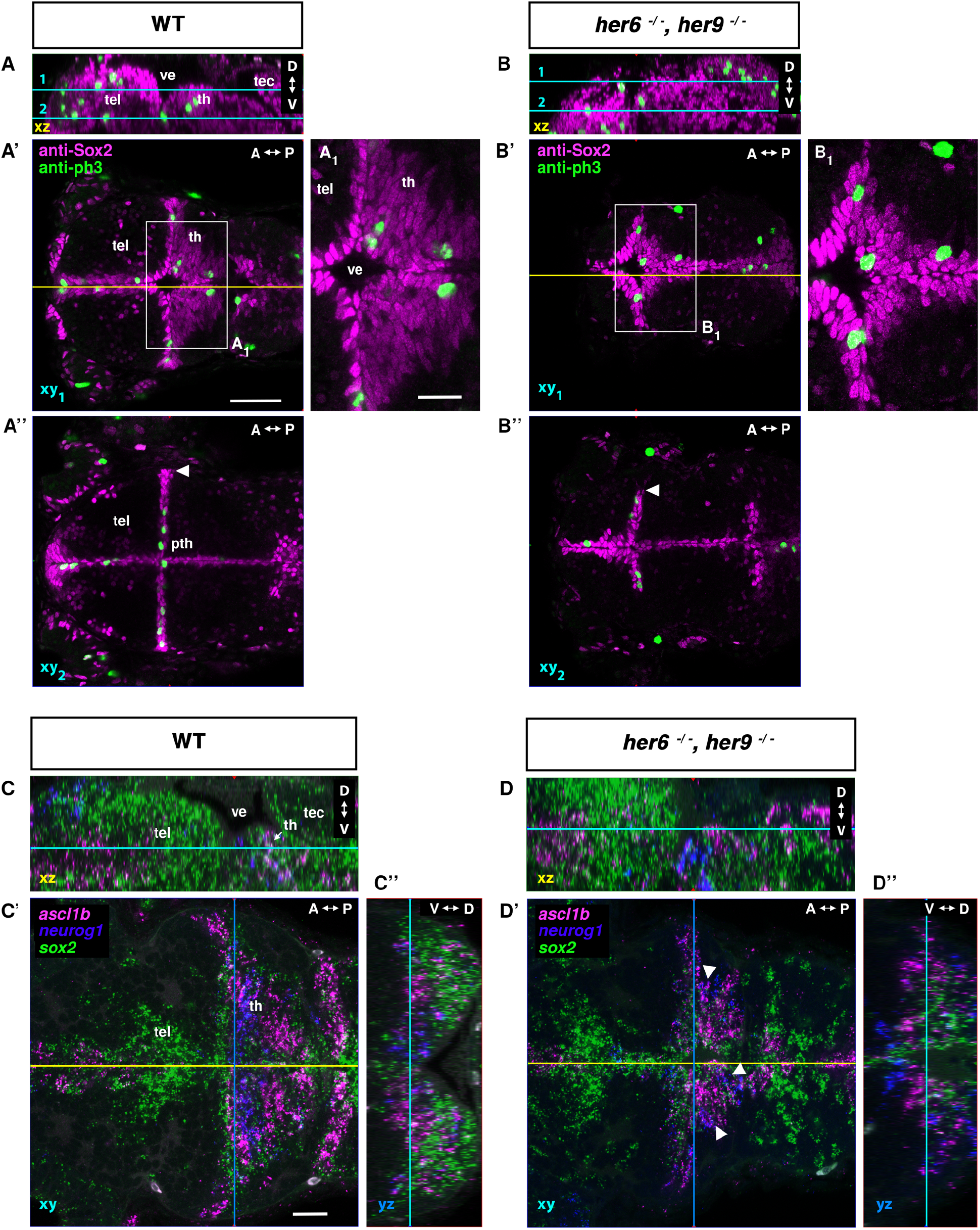
Neural stem and progenitor cell phenotypes in *her6, her9* double mutants. **(A-B’’)** Confocal image stacks of anti-Sox2 and anti-phospho-Histone H3 (pH3) immunofluorescence of 3 dpf WT control and *her6, her9* double mutants. **(A, B)** Lateral view midline sagittal XZ plane from orthogonal reconstructions, cyan lines indicating the dorsal view horizontal confocal planes 1 and 2 shown in A’, A” and B’, B”, respectively. The yellow lines in A’ and B’ indicate the midline sagittal planes in A and B. Plane 1 in A’ and B’ shows the dorsal part of the diencephalon with the thalamus proper and plane 2 in A’’ and B’’ is located more ventrally including the prethalamus. A_1_ and B_1_ are magnifications of the boxed areas in A’ and B’ respectively. Scale bar in A’ (for A-A”, B-B’’) is 50 μm; scale bar in A_1_ (also for B_1_) is 20 μm. The whole confocal stacks shown in A and B are also available as a combined Movie M2. **(C-D’’)** Whole mount fluorescent hybridization chain reaction (HCR) for detection of *sox2* (green), *ascl1b* (magenta) and *neurog1* (blue) expression. **(C, D)** Midline sagittal views of orthogonal reconstructions with cyan lines indicating the dorsal view horizontal image planes in C’ and D’. **(C’, D’)** Horizontal planes with blue lines indicate the level of frontal orthogonal reconstructions shown in C” and D”. C” and D” are frontal views at the lateral expansion of the third ventricle. Scale bar in C’, 20 μm. Numbers of embryos analyzed: A n=5; B n=4; C n=3; D n=3. Abbreviations: A-P indicates anterior-posterior, D-V dorsal-ventral orientation. pth, prethalamus; tec, tectum; tel, telencephalon; th, thalamus proper; ve, ventricle.

To determine whether the loss of Sox2 positive cells may correlate with reduced cell proliferation, we used an anti-phospho-Histone H3 (pH3) antibody to detect cells in mitosis. Compared to WT, *her6, her9* double mutants had a similar distribution of pH3 positive mitotic nuclei in the TPZs ventricular zone (Fig. 4A’,B’ and Movie M2; nuclei not counted). However, pH3 positive nuclei in *her6, her9* double mutants were largely absent in the lateral ventricle region, which is also devoid of Sox2 positive cells. *ccnd1* expression, which accumulates during G1 to S phase transition, was reduced in the TPZ especially along the lateral ventricle of *her6, her9* double mutants (Fig. S8). Thus, while mitotic activity of Sox2^high^ TPZ cells is not impaired, there are no mitotically active cells observed in the lateral ventricle wall devoid of Sox2^high^ cells.

Next, we investigated proneural gene expression in the Sox2^low^ NPCs in the TPZ. Fluorescent HCR-RNA FISH shows that *sox2* in Sox2^low^ regions is co-expressed with proneural genes (Fig. 4C-C”). The strongest co-expression is in the thalamus proper in Sox2^low^ expressing cells that are not in direct contact with the ventricle (Fig. S5A-F and Movie M1). Here, *sox2* expression was overlapping with *neurog1* or *ascl1b* in Sox2^low^ NPCs. More precisely, *ascl1b* is expressed in a narrow stripe in the rostral thalamus, while *neurog1* is expressed in the caudal thalamus (Fig. 4CC”; *ascl1a* is also expressed in prethalamus and pretectum, see also Fig. 11A-C). In *her6, her9* double mutants however, *neurog1* and *ascl1b* expression domains were intermingled (arrowheads) and the area of expression is smaller and narrower (Fig. 4D-D”). Taken together, we found *her6, her9* double mutants to have a smaller domain of *sox2^low^* expressing progenitors, and lack the proper spatial organization of *neurog1* and *ascl1b* NPCs.

Next, we tested whether loss of Sox2 positive NSCs may correlate with changes in the ZLI. In WT, *shha* in the thalamus is expressed in the basal plate and the ZLI (Scholpp and Lumsden, 2010). As development progresses, *shha* expression is maintained medially in close proximity to the ventricle and expands dorsally into the ZLI, where the expression extends along the walls of the lateral ventricle (Fig. 5A-D,I). While at 24 hpf the *shha* expression in the ZLI appears largely normal in *her6, her9* double mutants (Fig. 5A,A’,E,E’), at 72 hpf and 96 hpf we observe a progressive loss of the *shha* expression domain that normally demarcates the ZLI separating the prethalamus from the thalamus proper, both in its lateral and dorsal domains (Fig. 5). A partial loss of *shha* expression along the lateral ventricular walls was also observed in the ZLI of *her6* single mutants (Fig. S9), although the extent of loss of *shha* expression in this region varied among larvae (Fig. S9C’ and D’, white arrowheads). We interpret this variability as gradual loss of the ZLI cell population that forms this *shha* expression domain.

**Fig. 5.**
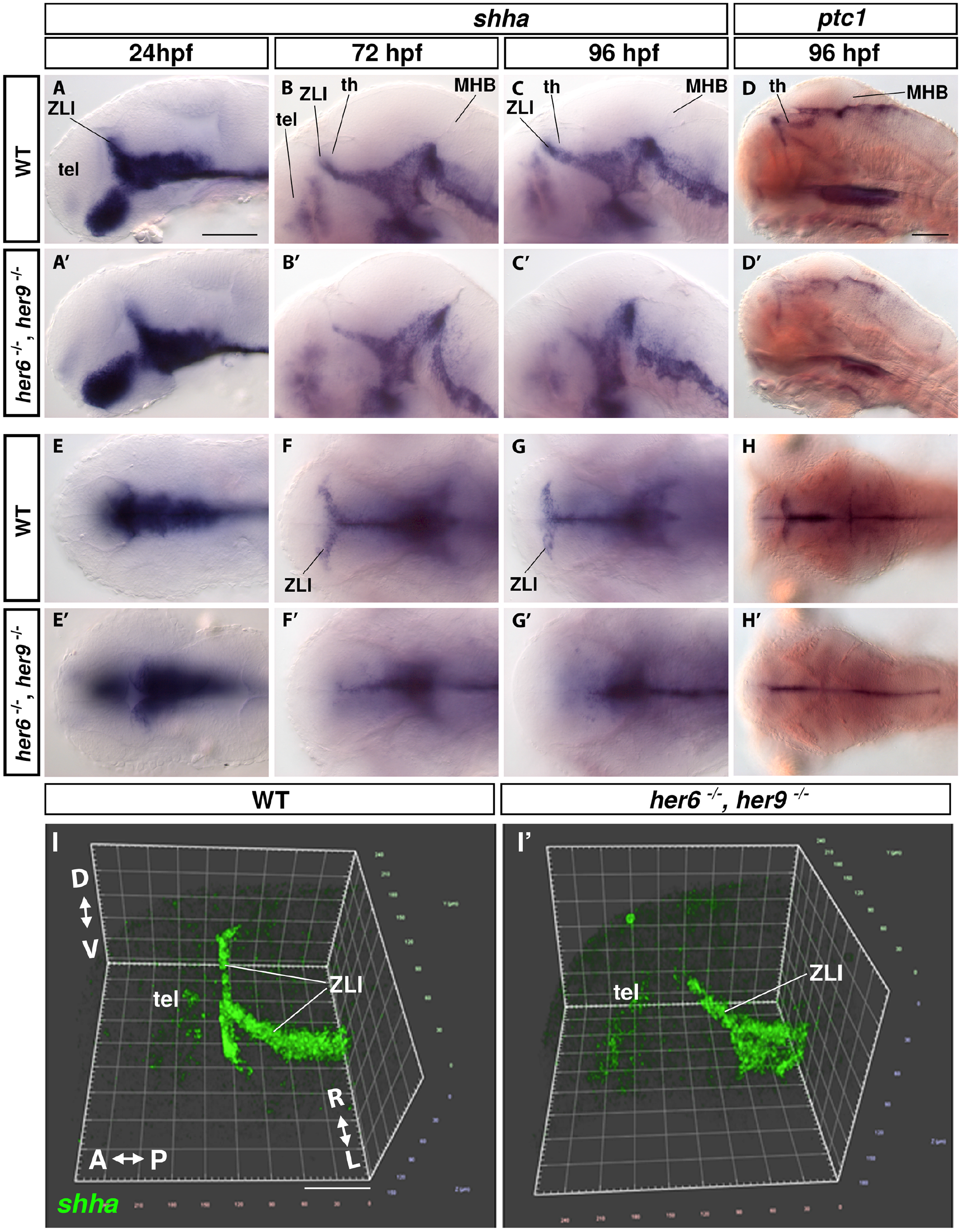
Loss of zona limitans intrathalamica *shha* expression in *her6, her9* double mutants. **(A-H’)** WISH showing *shha* (A-C’, E-G’) and *ptc1* (D, D’, H, H’) expression in WT and *her6, her9* double mutants. Genotypes are shown at left and the analyzed stages at top. **(A-D’)** Lateral views show single sagittal image planes at the midline. **(E-H’)** Dorsal views show single horizontal image planes at the level of the thalamus. Three larvae were imaged per condition and one representative image was selected. Expression of *shha* in *her6* and *her9* single mutants is shown in Fig. S9. Scale bar in A, 100 μm (for A-C’ and E-G’); scale bar in D, 100 μm (for D, D’, H, H’). **(I, I’)** 3D volume reconstruction of fluorescent WISH of *shha* in 3 dpf WT (I) and *her6, her9* double mutant (I’). Scale bar x-axis, 60 μm. A-P indicates anterior-posterior, D-V the dorsal-ventral, L-R the left-right orientation. Numbers of embryos analyzed: I n=3; I’ n=4. Abbreviations: MHB, midbrain-hindbrain boundary; tel, telencephalon; th, thalamus proper; ZLI, zona limitans intrathalamica.

### Loss of the two Notch-dependent *her* gene clusters does not affect NSCs and NPCs in the TPZ

The two *her4;12* and *her2;15* gene clusters active in the TPZ contain nine *Hes5* zebrafish homologs (Zhou et al., 2012), but no other genes (www.ensembl.org, accessed on 30. August 2021). To genetically inactivate these nine *her* genes, we generated large precise deficiencies in both chromosomal regions (Fig. 6A,B and Fig. S10). The 28 kb deficiency *Df(Chr23:her12,her4.1,her4.2,her4.3,her4.4,her4.5)m1364* or *m1365*, abbreviated here as *Df(her4;12)*, and the 30 kb deficiency *Df(Chr11:her2,her15.1,her15.2)m1490*, abbreviated here as *Df(her2;15)* were generated using the CRISPR/Cas9 system. The deletions were confirmed by sequencing (Fig. S10). WISH for *her4* and *her15* demonstrated that there were no transcripts left in theses mutants, and no translocations of the deleted sequences occurred (Fig. 6C-D’).

**Fig. 6.**
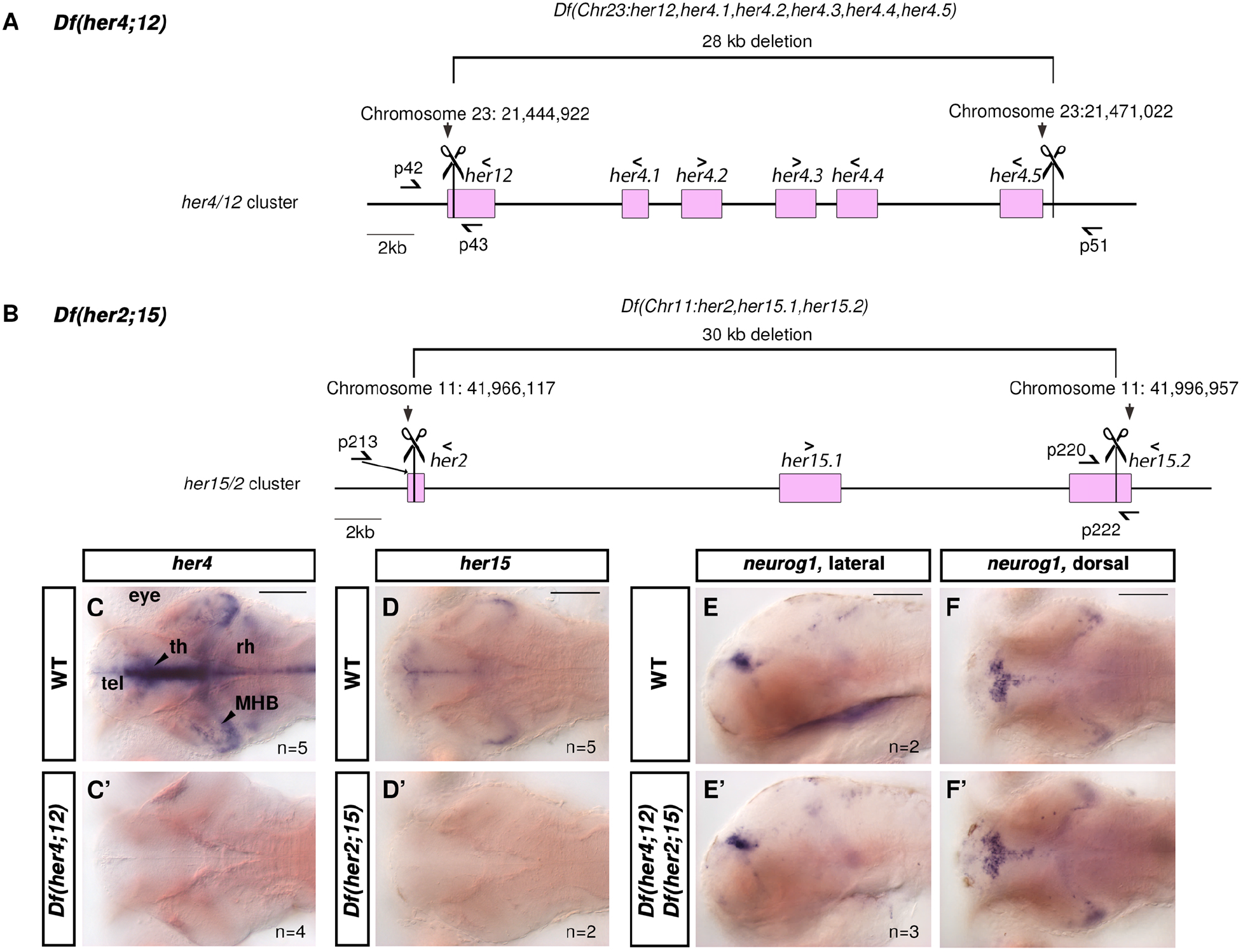
Genome editing strategy for *Df(her4;12)* and *Df(her2;15)* large chromosomal deletions: verification and *neurog1* expression. **(A, B)** Generation of large deficiencies using the CRISPR/Cas9 system. Scissors indicate the sgRNAs sequence positions, and square brackets indicate the deletion. 5’>3’ or < signs indicate the orientations of all genes (pink boxes) in the intervals. Half-arrows indicate binding sites of primers used in genotyping PCRs (tagged with primer names). **(C-F’)** WISH for *her4, her15* and *neurog1* mRNAs in 96 hpf larvae. No *her4* or *her15* signal is detectable in the respective mutants (C-D’, dorsal view horizontal optical planes). No change in *neurog1* expression is detectable in *Df(her4;12), Df (her2;15)* double deficiency mutants (E and E’, midsagittal; F and F’, horizontal optical sections). The *her4* probe detects *her4.1 to her4.5* transcripts, and the *her15* probe detects *her15.1* and *her15.2* transcripts. Numbers of embryos analyzed are indicated in lower right corners. Abbreviations: MHB, midbrain-hindbrain boundary; rh, rhombencephalon; tel, telencephalon; th, thalamus proper. Scale bars, 100 μm.

We could not observe any morphological changes in *Df(her4;12)* or *Df(her2;15)* mutant larvae until 30 dpf (data not shown) or in fixed 3 dpf larvae. Therefore, we generated larvae devoid of all nine Notch-dependent *her* genes in crosses of *Df(her4;12)*, *Df(her2;15)* double deficiency heterozygous parents. Surprisingly, double deficiency homozygous larvae also appeared morphologically largely normal. To assess whether thalamic neurogenesis may be affected, we analyzed *neurog1* expression and found it also unchanged (Fig. 6E-F’). To determine whether the NSC compartment was affected in these mutants, we performed immunofluorescence for Sox2 and pH3 (Fig. 7A-B” and Movie M3). The distribution of pH3 positive nuclei appeared unchanged in *Df(her4;12), Df(her2;15)* double mutants compared to WT embryos (Fig. 7A-B”) and the expression of *ccnd1* was unchanged (Fig. S8), suggesting normal proliferation in the TPZ. The Sox2^high^ nuclei layer covering the ventricular wall of the thalamus proper appeared intact (small arrow in Fig. 7A_1_,B_1_), and also the adjacent subapical Sox2^low^ nuclei appeared normal (arrowhead in Fig. 7A_1_,B_1_). The shape of the nuclei at the ventricular wall was spherical, and the subapical Sox2^low^ nuclei were elongated, both in WT and *Df(her4;12), Df(her2;15)* double mutants. Sox2^high^ nuclei lined the lateral ventricle along its full lateral extent (Fig. 7A” and B” arrowheads). Thus, NSCs appear to develop normally in the TPZ devoid of Notch-dependent *her* genes.

**Fig. 7.**
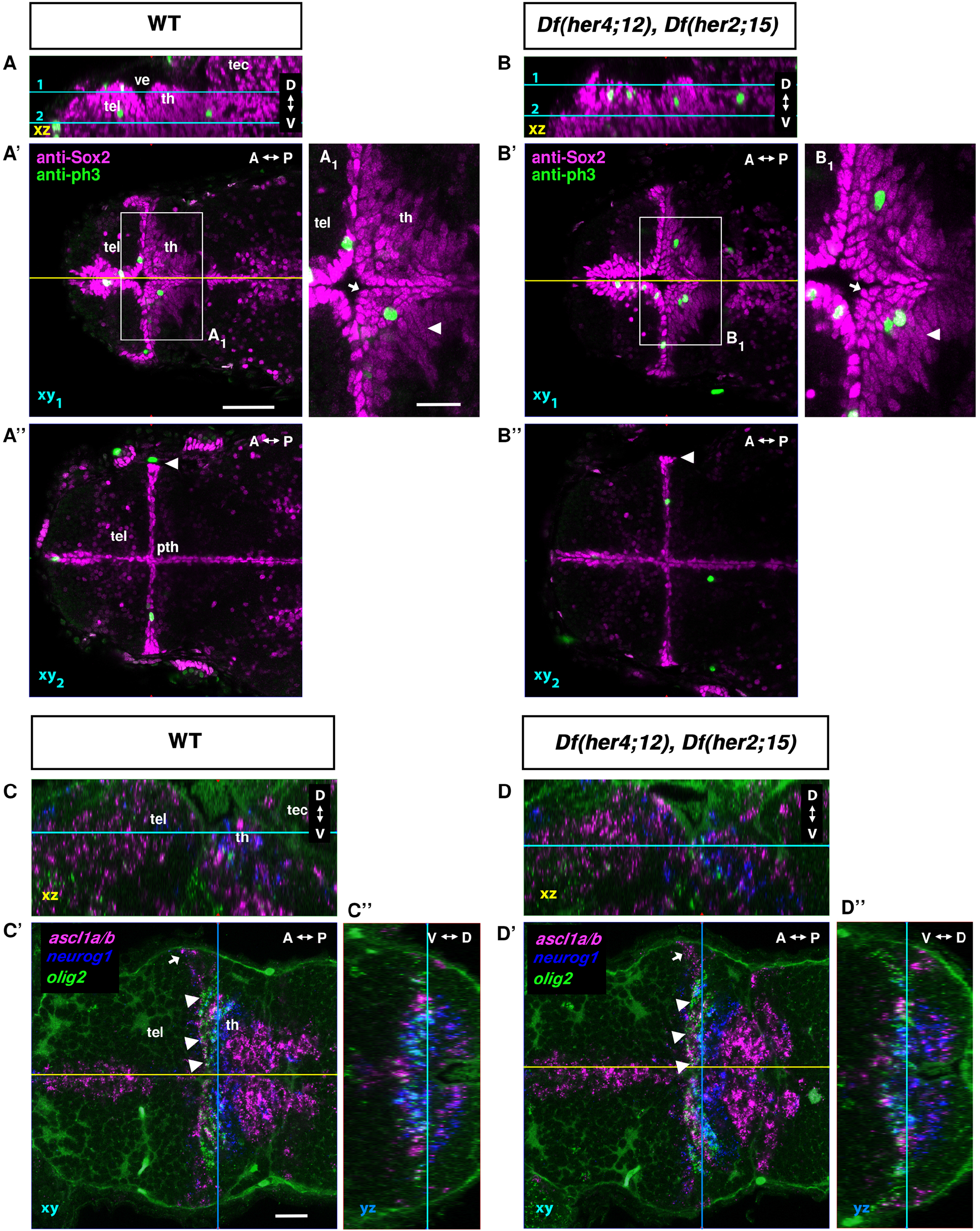
Sox2 and proneural gene expression in *Df(her4;12), Df(her2;15)* double deficiency mutants. **(A-B’’)** Confocal image stacks of anti-Sox2 and anti-pH3 immunostaining in 3 dpf WT **(A)** and *(her4;12); Df(her2;15)* double deficiency mutants **(B)**. **(A, B)** Sagittal midline XZ plane from orthogonal reconstructions, cyan lines indicating the dorsal view confocal planes 1 and 2 shown in A’, A’’ and B’, B’’. The yellow lines in A’ and B’ indicate the midline sagittal planes in A and B. Plane 1 in A’ and B’ shows the dorsal part of the diencephalon with the thalamus proper and plane 2 in A’’ and B’’ is located more ventrally showing the prethalamus. A_1_ and B_1_ are magnifications of the boxed area in A’ and B’, respectively. Scale bar in A’, 50 μm (for A-B’’); scale bar in A1, 20 μm (also for B1). The complete confocal stacks of larvae shown in A and B are combined in Supplementary Movie M3**. (C-D’’)** Whole mount fluorescent HCR mRNA detection for *olig2* (green), *neurog1* (blue) and *ascl1a/b* (magenta) in **(C-C’’)** WT and **(D-D’’)** *Df(her4;12); Df(her2;15)* double deficiency mutants. **(C, D)** Sagittal midline XZ plane from orthogonal reconstructions, with cyan lines indicating the confocal planes in C’ and D’. **(C’, D’)** Dorsal view horizontal confocal planes of the diencephalon. Blue lines indicate the frontal view orthogonal reconstruction shown in C’’ and D’’. Numbers of embryos analyzed: A n=4; B n=5; C n=3; D n=3. Scale bar in C’, 20 μm. Abbreviations: A-P indicates anterior-posterior, D-V dorsal-ventral orientation. pth, prethalamus; tec, tectum; tel, telencephalon; th, thalamus proper; ve, ventricle.

We also analyzed *her* gene expression in *her6, her9* and in *Df(her4;12), Df(her2;15)* double mutants (data not shown). *her6* and *her9* expression is largely unaffected in *Df(her4;12)*, *Df(her2;15)* double mutants in line with their Notch-independent regulation. Based on WISH, *her4* and *her15* expression appear slightly increased in the NPC area of *her6* and *her9* double mutants, but at the ventricular wall *her4* and *her15* expression do not invade into the *her6* expression domains in the rostral thalamus and prethalamus (data not shown), suggesting that mechanisms other than repression by *her6* limit *her4* and *her15* expression in these domains.

To determine whether also progenitor cells appear normal, we analyzed expression of *neurog1*, *ascl1a/b* and *olig2. olig2* is expressed in a stripe of cells parallel to the ventricular wall, adjacent to the expression of *neurog1* (Fig. 7C-C”, arrowheads). Both the *olig2* and *neurog1* expression domains appeared unchanged in *Df(her4;12), Df(her2;15)* double deficiency mutants (Fig. 7D-D”, arrowheads). Additionally, thalamic *ascl1a/b* expression domains appeared normal in double deficiency mutants (white arrows). Although more subtle changes in expression cannot be excluded, the periventricular stem and progenitor cell compartments of the TPZ appeared very similar to WT. Thus, surprisingly, in the absence of most Notch-dependent *her* genes, NSC and NPC populations appear unaffected. This is in stark contrast to the defects observed in the *her6*, *her9* double mutant.

### Her4 and Her6 overexpression reveal selective cross-regulation of *her* genes

We next analyzed potential cross-regulation between Notch-dependent and -independent *her* genes by heat shock driven overexpression of Her4 or Her6 using *Tg(hsp:her4-FLAG)* and *Tg(hsp:her6-FLAG)* transgenic lines. When *Tg(hsp:her4-FLAG)* larvae were analyzed by WISH 2.5 hours post heat shock, *her4* was confirmed to be overexpressed (Fig. 8A,A’), and we detected downregulation of *her15* expression (Fig. 8B,B’). Her4 overexpression however did not affect the expression levels of *her6* and *her9* (Fig. 8C-D’). In contrast, overexpression of Her6 strongly downregulated *her9* and both *her4* and *her15* (Fig. 8E-H’). This suggests that Notch-independent *her* genes negatively cross-regulate Notch-dependent *her* gene expression, while Notch-dependent *her* genes negatively cross-regulate other Notch-dependent *her* genes, but do not cross-regulate Notch-independent *her* gene expression.

**Fig. 8.**
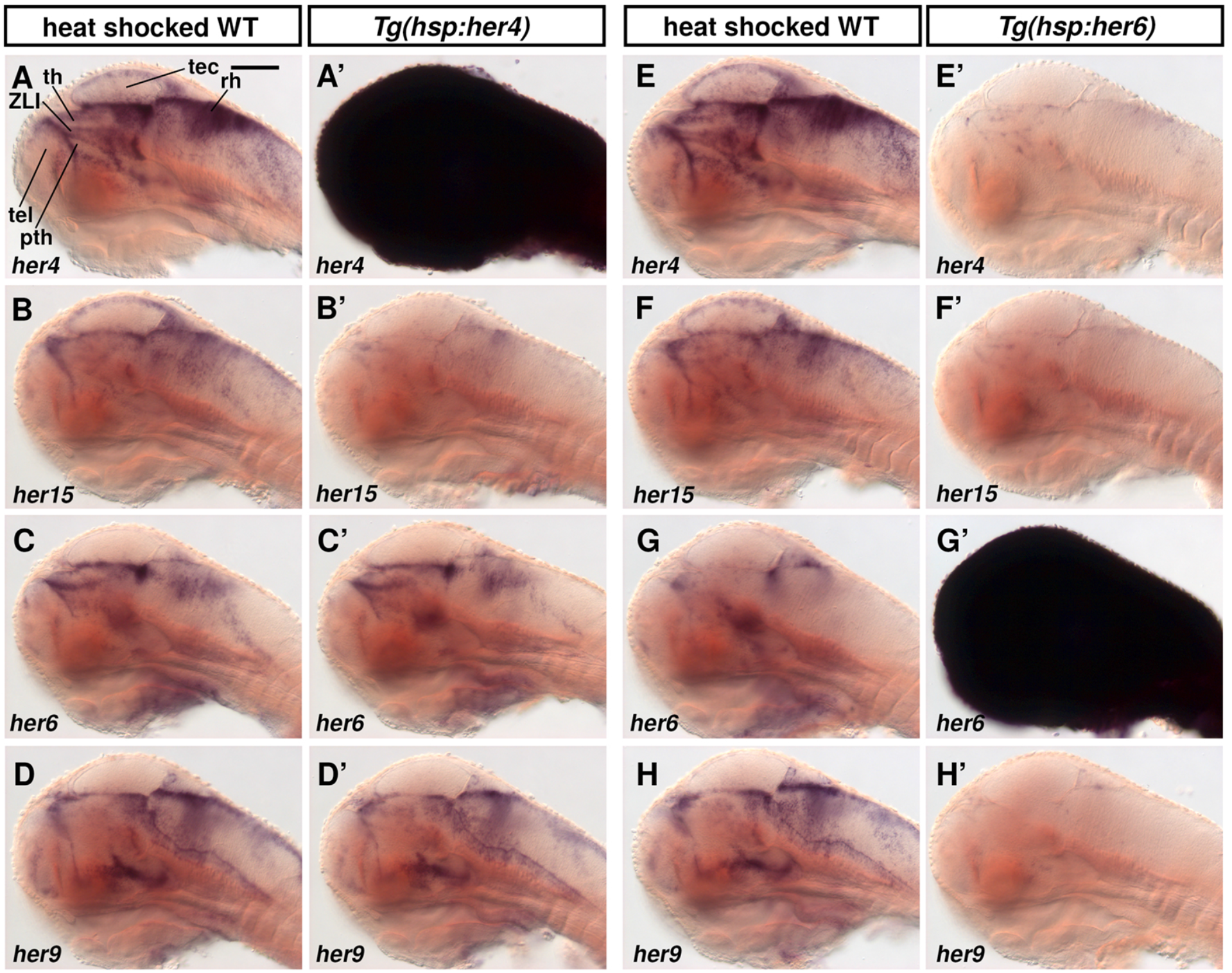
Overexpression of Her4 and Her6 reveals differential cross-regulation of *her* genes. **(A-H’)** WISH for *her* gene expression after heat shock induced overexpression of Her4 or Her6 in *Tg(hsp:her4-FLAG)* **(A’-D’)** and *Tg(hsp:her6-FLAG)* **(E’-H’)** transgenic embryos. **(A-H)** WT heat shocked sibling controls. A’ and G’ validate heat shock mediated high expression of *her4-FLAG* and *her6-FLAG* transgene transcripts. All larvae were heat shocked at 70 hpf for 30 min and were fixed at 72.5 hpf, except for G and G’ that were fixed at 71.5 hpf because *her6* transcript levels decrease faster after heat shock than *her4* transcript levels (Sigloch and Driever, unpublished). Three larvae were imaged per condition and one representative image is shown. The *her4* probe detects *her4.1-her4.5* and the *her15* probe detects *her15.1* and *her15.2.* Sagittal optical sections close to midline, anterior at left, dorsal up; scale bar, 100 μm. Abbreviations: pth, prethalamus; rh, rhombencephalon; tec, tectum; tel, telencephalon; th, thalamus proper; ZLI, zona limitans intrathalamica.

### Overexpression of Her6, but not Her4, strongly downregulates proneural gene expression

We next tested whether Her4 or Her6 might differentially regulate proneural gene expression by heat-shock driven overexpression at 70 hpf and fixation 2 hours after heat shock. Upon Her4 overexpression, surprisingly, *neurog1* expression appeared largely unchanged, while *ascl1b* expression was reduced in 9 out of 14 larvae (Fig. 9A-B’). This finding is surprising, because a strong effect of Notch-dependent *her* genes on proneural gene expression would be expected (Imayoshi et al., 2013; Takke et al., 1999). *neurod1* and *neurod6b* expression were unchanged two hours after *her4* heat shock (Fig. 9C-D’). *sox2* expression was slightly reduced in 9 out of 15 *her4* heat shocked larvae (Fig. 9E-E’). In contrast, overexpression of Her6 strongly downregulated expression of *neurog1* and *ascl1b*, and, to a lesser extent, also of the late progenitor markers *neurod1* and *neurod6b* (Fig. 9F-I’). In addition, expression of the stem cell marker *sox2* appeared slightly reduced in 11 of 15 Her6 heat shock larvae (Fig. 9J-J’), but more strongly affected than by heat-shock overexpressed Her4. We also tested whether overexpression of Her4 or Her6 might have a prolonged effect on proneural gene expression late after heat shock. We analyzed larvae 6 hours after beginning of the 30 min heat shock and found *neurog1* and *ascl1b* expression at normal levels (Fig. S11A-D’). Since *her6* consistently downregulated *neurog1* and *ascl1b* 2.5 hrs after heat shock, their expression appears to be back to normal only a few hours after overexpression of *her6* ends. This suggests that the NSC regulatory network is robust against perturbations in Her6 expression level, and rapidly re-establishes normal activity. Together, our findings reveal a strong regulatory impact of Her6 on progenitors and stem cells, and indicate that Notch-independent Her6 might be a much more potent inhibitor of neurogenesis compared to Notch-dependent Her4.

**Fig. 9.**
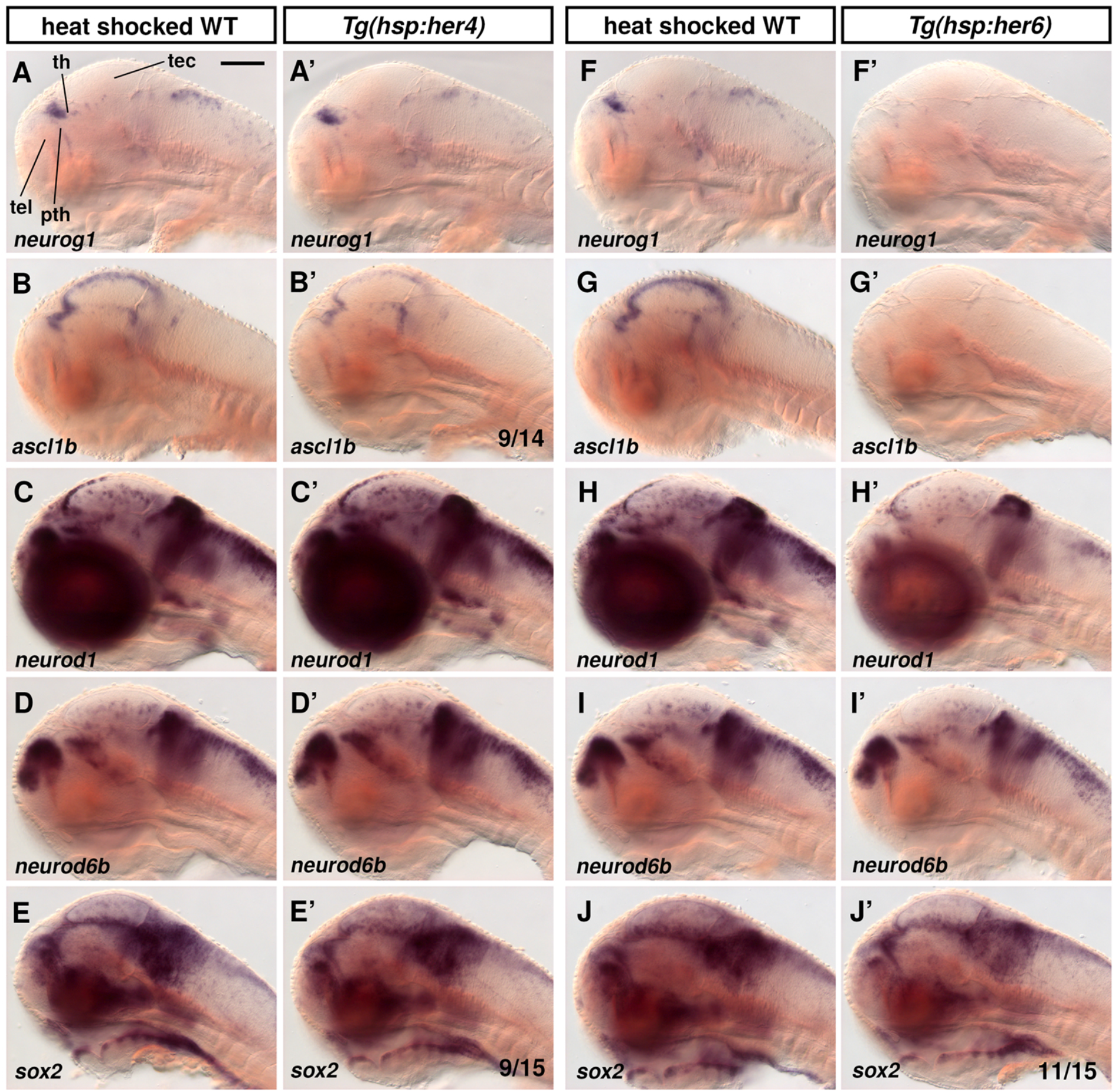
Overexpression of Her4 and Her6 differentially affect proneural gene expression. **(A-J’)** Expression of proneural genes and *sox2* after heat shock induced overexpression of Her4 or Her6. **(A’-E’)** *Tg(hsp:her4-FLAG) and* **(F’-J’)** *Tg(hsp:her6-FLAG)* were used to overexpress Her4 or Her6 by heat shock at 70 hpf for 30 min, and embryos were fixed at 72.5 hpf. (**A-J)** WT siblings were heat shocked as controls. For numbers of embryos analyzed see Supplementary Table 4. Analyzed conditions showed consistent WISH expression patterns in all embryos except B’, E’ and J’, for which the most representative patterns are shown and their numbers out of total analyzed provided in bottom right of these panels. Sagittal optical sections close to midline, anterior at left, dorsal up; scale bar, 100 μm. Abbreviations: pth, prethalamus; tec, tectum; tel, telencephalon; th, thalamus proper.

### Combined loss of activity of eleven Notch-dependent and -independent *her* genes

We next asked whether Notch-dependent and -independent *her* genes may functionally compensate for each other despite their different regulation. We genetically combined the two deficiencies eliminating all nine *Hes5* homologs together with mutant alleles of *her6* and *her9*, and named embryos homozygous mutant for all eleven *her* genes “*her* undecimal mutants” (*her*UDM; for *Df(her4;12), Df(her2;15), her6, her9).* The brains of *her*UDM embryos appeared severely malformed, and embryos did not survive beyond 5 dpf. When analyzed at 56 hpf (Fig. 10A-B’), *her*UDM embryos had the most severe morphological defects in neural tissues that undergo significant expansion by proliferation on the second day of development, such as the cerebellum, the ventral part of the retina, and the rhombic lip. Early forming neural tissues however appeared largely normal, which might be due to compensation by other *her* genes expressed during early stages of development, including *her3, her8a and her8.2* (Onichtchouk et al., 2010; Webb et al., 2011).

**Fig. 10.**
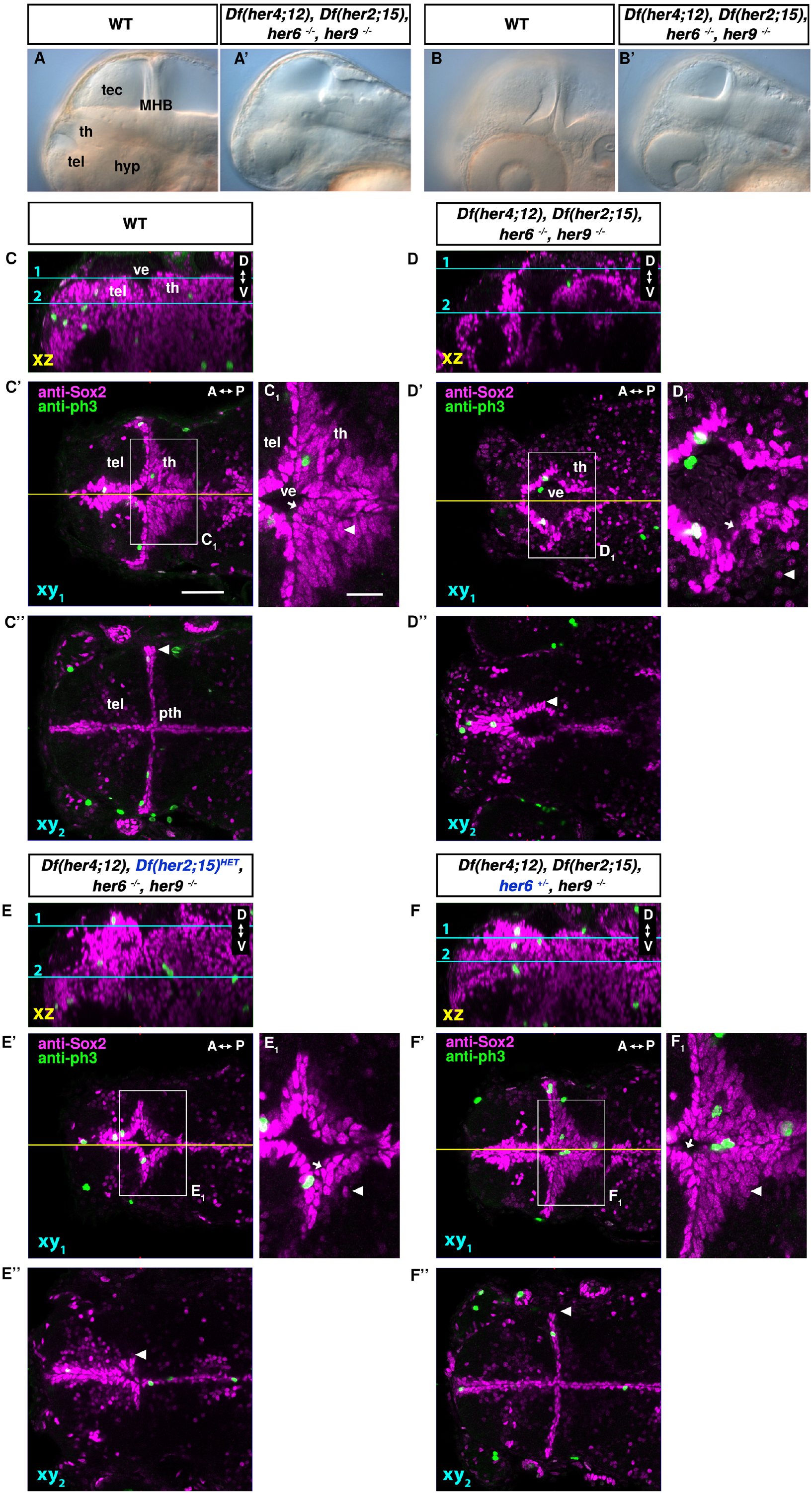
Analysis of the *her* activity depleted *her*UDM mutant phenotype. **(A-B’)** Live phenotype of WT and *Df(her4;12), Df(her2;15), her6, her9* combined mutant (*her*UDM) embryos, lateral views at 56 hpf; (A) midsagittal optical section, (B) parasagittal optical section at level of lens. **(C-F“)** Anti-Sox2 and anti-pH3 immunofluorescence in 3 dpf WT **(C-C’’)** and *her*UDM combined mutant (D-F’’) embryos as indicated. **(E-E’’)** Triple *(her4;12), her6, her9* mutant *Df(her2;15)* heterozygous embryos showing minimal rescue of *her*UDM by one copy of the WT chromosomal region including *her2, her15.1, her15.2.* **(F-F’’)** Triple *(her4;12), Df(her2;15), her9* mutant and *her6* heterozygous embryo revealing efficient rescue of *her*UDM by one intact *her6* allele. **(C, D, E, F)** Mid-sagittal optical sections generated from orthogonal reconstruction of confocal stacks, cyan lines indicating the horizontal confocal planes 1 and 2 shown in C’, C’’, D’, D’’, E’, E’’ and F’, F’’. The yellow lines indicate the sagittal midline sections shown in C, D, E and F. Plane 1 in C’, D’, E’ and F’ shows the dorsal part of the diencephalon and plane 2 in C’’, D’’, E’’ and F’’ is located more ventrally. C_1_, D_1_, E_1_ and F_1_ are magnifications of the respective boxed areas. A-P indicates anterior-posterior, D-V dorsal-ventral orientation. Scale bar in C’ 50 μm for C-F’’; scale bar in C_1_ 20 μm for C_1_-F_1_. The confocal stacks in C and D as well as in E and F are provided as combined Supplementary Movies M4 and M5. Numbers of embryos analyzed: A n=1 (WT); A’ n=3; C n=4; D n=4, E n=3 (in Fig. 10E, a *her2, her15* heterozygous larva is shown. Two more imaged larvae were WT for *her2, her15*), F n=4 (in Fig. 10F, a *her6* heterozygous larva was used. Three more imaged larvae were WT for *her6*). Abbreviations: hyp, hypothalamus; MHB, midbrain-hindbrain boundary; pth, prethalamus; tel, telencephalon; th, thalamus proper; ve, ventricle.

We analyzed Sox2 expression in *her*UDM embryos and observed a severe but quite variable overall phenotype, ranging in the TPZ from strong reduction of both the Sox2^high^ NSC and the Sox2^low^ NPC compartments (as shown in Fig. 10D-D”) to extreme anatomical malformations rendering analysis of the TPZ difficult (not shown). The Sox2^high^ expressing nuclei in 3 dpf *her*UDM embryos along the ventricular wall appeared disorganized compared to WT in that the ventricular layer of Sox2^high^ nuclei was interrupted (Fig. 10C-D” and Movie M4), and some Sox2 positive nuclei were displaced from the ventricular wall, suggesting that the epithelial integrity of the ventricle may be impaired (Fig. 10C_1_ and D_1_, arrows). Similar loss of neuroepithelial integrity had been reported for *Hes1, Hes5* double mutant mice (Hatakeyama et al., 2004). In the lateral view, it appeared as if there were less Sox2 positive nuclei lining the ventricle wall, which might however also reflect the altered shape of the ventricle in the mutant (Fig. 10C,D). Similar to *her6, her9* double mutants, the Sox2^low^ nuclei located subapically away from the ventricular wall of the thalamus proper were nearly absent and did not show the elongated shape typical for WT NPCs (Fig. 10C_1_,D_1_, arrowheads). The layer of Sox2^high^ nuclei observed in WT along the walls of the lateral ventricle was completely missing (Fig. 10C”,D”). In the diencephalon of *her*UDM embryos we still detected pH3 positive nuclei at the ventricular walls (Fig.10D’ and Movie M4), suggesting that NSC proliferation is not completely eliminated.

We tested whether a single WT allele of a Notch-dependent or -independent *her* gene may rescue the *her*UDM phenotype. A single WT allele of the *her2;her15* gene cluster in *Df(her4;12)* homozygous*, Df(her2;15)* heterozygous, *her6^-/-^, her9^-/-^* mutant larvae did partially rescue the *her*UDM phenotype (Fig. 10E-E” and Movie M5, right). The integrity of the ventricular wall as defined by Sox2^high^ cells was restored and resembled the phenotype in *her6, her9* double mutants (Fig. 10E1, arrow, E”, and Movie M5). Like in the *her6, her9* double mutant, the subapical Sox2^low^ positive nuclei were still missing (Fig. 10E1, arrowhead and Movie M5). In contrast, a single *her6* WT allele in *Df(her4;12), Df(her2;15), her6^+/-^, her9^-/-^* mutant larvae fully restored the Sox2 expressing nuclei along the lateral ventricular wall (Fig. 10F” arrowhead, and Movie M5, left). The ventricular epithelium appeared intact (Fig. 10F_1_, arrow), and the subapical Sox2^low^ immunoreactive NPC nuclei were present and had the normal elongated nuclear shape (Fig. 10F’ and F_1_, arrowhead). Individual effects of *her9* or *her4,12* were not tested, because they are tightly linked on the *her9 ^m1505^, Df(her4;12)* chromosome 23 we used.

In summary, the *her*UDM phenotype revealed that, first, the eleven *her* genes analyzed provide most if not all of the Her activity in the TPZ at 3-4 dpf, given the severe loss of NSCs and NPCs. Second, Notch-dependent and -independent *her* genes appear to act partially redundant in NSC maintenance. Third, the Notch-independent *her6* makes the most predominant contribution to NSC and NPC maintenance and expansion.

## Discussion

Neural proliferation zones mediate brain growth, and need to balance NSC maintenance with generation of NPCs and neurons during neurogenesis. Delta/Notch signaling and HES/HER transcription factors play important roles, both in NSC maintenance and control of NPC progression in neurogenesis (Kageyama et al., 2007). HES/HER activities differ during primary neurogenesis in the early neuroepithelium of anamniotes, during embryonic brain growth in neural proliferation zones, and in adult neural stem cell niches (Alunni et al., 2020; Kriegstein and Alvarez-Buylla, 2009). Here, we analyzed Her activities in the zebrafish TPZ, which is highly active in neurogenesis at late embryonic and early larval stages.

Among the HES/HER gene families (Chapouton et al., 2011; Zhou et al., 2012) we detected zebrafish homologs of mammalian Hes1, Hes5, and Hes6 expressed in the TPZ. *Hes1* is predominantly regulated by Notch signaling (Hsieh et al., 1997; Jarriault et al., 1995; Jarriault et al., 1998; Nishimura et al., 1998), but also by Notch-independent input (Kageyama et al., 2007; Ohtsuka et al., 1999). In contrast, we find expression of the *Hes1* homologs *her6* and *her9* in the larval NPZs to be largely independent of Notch signaling, in line with previous reports on earlier stages (Bae et al., 2005; Hans et al., 2004; Latimer et al., 2005). *Hes5* expression in mice is Notch-dependent (Ohtsuka et al., 1999). Similarly, for the zebrafish *Hes5* homologs *her4* (Takke et al., 1999; Zhou et al., 2012), *her12* (Gajewski et al., 2006), *her2* and *her15* (Cheng et al., 2015; Shankaran et al., 2007), we find expression to depend on Notch signaling in NPZs. For the zebrafish *Hes6* homologs *her8a*, we find expression in NPZs to depend on Notch signaling, while in the neural plate Notch-independent expression has been reported (Webb et al., 2011). Similarly, *her8.2* appears to be regulated in NPZs by both Notch-dependent and -independent mechanisms.

To identify distinct contributions of Notch-dependent and -independent *her* genes to NPZs, we characterized *her* expression and cell type composition of the TPZ in detail (Fig. 11A). At the boundary of thalamus and prethalamus, the ZLI cells (blue) at the ventricular surface express the stem cell marker Sox2, *shh*, and high levels of *her4* and *her15*, as well as low levels of *her9.* In the prethalamus and rostral thalamus, directly adjacent to the ZLI organizer, are Sox2 and, exclusively with respect to *her* genes, *her6* expressing NSCs (green). Within the thalamus, caudal to the *her6* expressing stripe of cells, are Sox2 positive NSCs expressing high levels of *her4* and *her15*, as well as low levels of *her9* (red), similar to the ZLI cells. In the layer of cells directly below the ventricular NSCs, we observed different populations of Sox2^low^ expressing NPCs, expressing low levels of *her4, her15*, and/or *her9* as well as proneural genes: *ascl1a* expression is higher in the rostral prethalamus and rostral thalamus (orange), while *neurog1* expression is higher in a stripe of prethalamic cells adjacent to the ZLI and in the caudal thalamus (yellow). Cells further distal from the ventricular surface are mature neurons, or late progenitors expressing *neurod* family members (grey). Given that TPZ NSCs appear to be characterized by either Notch-dependent or Notch-independent *her* expression, we name the NSCs expressing high *her6* NSCtype1, and those expressing high *her4* and *her15* NSCtype2. The different NSC types emerge during maturation of the thalamic complex, as early *her6* expression comprises the entire presumptive thalamic complex and becomes gradually restricted (Scholpp et al., 2009). The complementary expression of *Hes1* and *Hes5* homologs in zebrafish is consistent with previous findings on *Hes1* and *Hes5* expression in mice (Hatakeyama et al., 2004). However, there are differences: High *Hes1* cells in mice with absent *Ascl1* expression represent quiescent NSCs (Sueda et al., 2019), while NSCtype1 are proliferative. Quiescent versus active NSC populations may not have been established in the larval TPZ, but diversify later in development (Than-Trong et al., 2020).

**Fig. 11.**
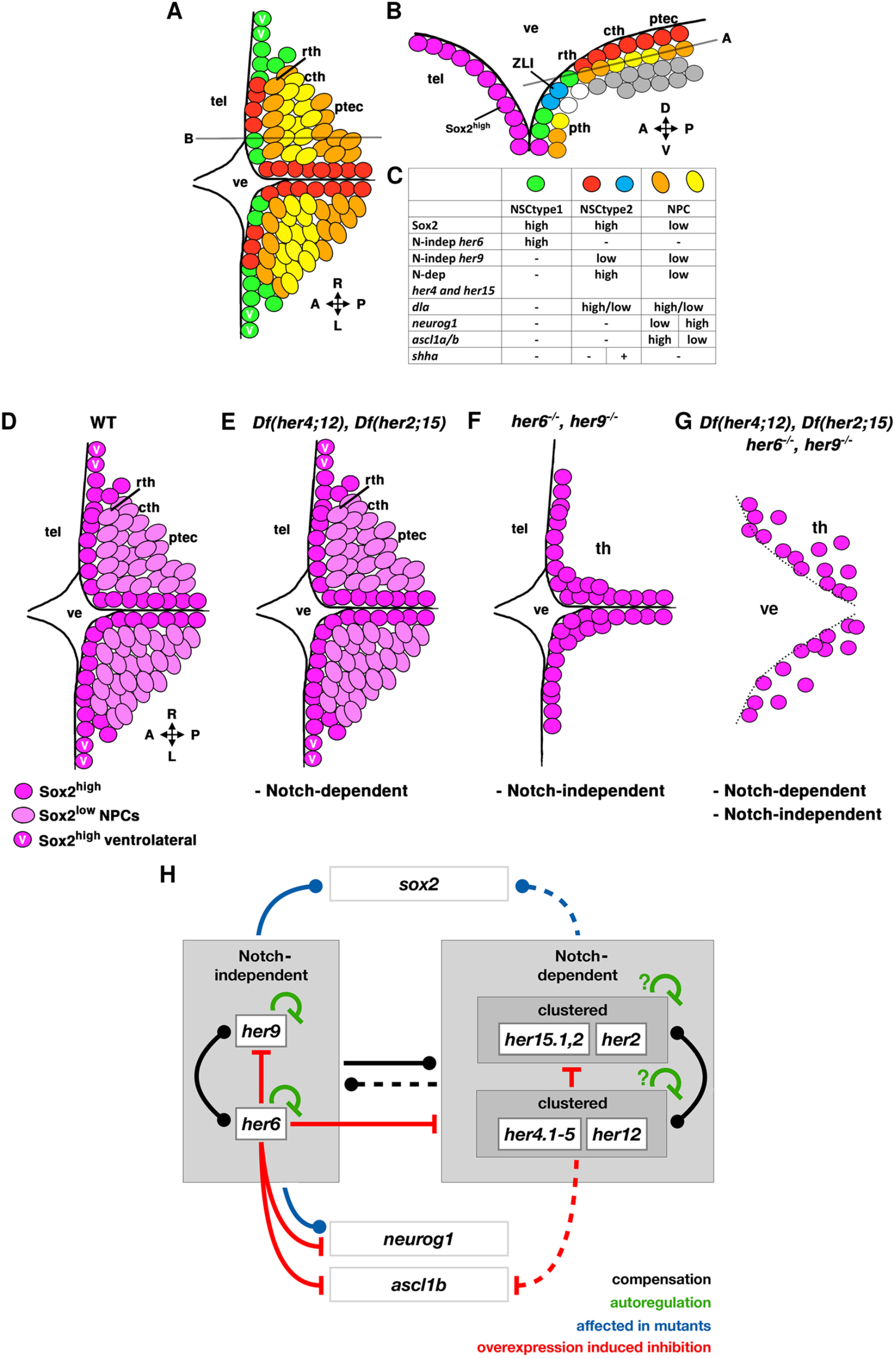
Model of the cellular organization, observed *her* deficient phenotypes and model for *her* genetic interactions in the TPZ. **(A-C)** Model of cellular organization in the TPZ. (A) Horizontal plane dorsal view, plane indicated in (B), which shows parasagittal section as indicated in (A). The table in panel C indicates marker gene expression in different color-coded cell populations identified. Cells expressing *her6* are categorized as NSCtype1; Sox2^high^ cells expressing *her4* and *her15* are categorized as NSCtype2 and expression of *neurogenin1* and *ascl1a/b* mark NPCs. Magenta cells are high Sox2 expressing cells of the telencephalic ventricular wall, *her* expression not determined. White cells were not analyzed for expression of markers. Grey cells summarize differentiating early and mature neurons, some of which express *neurod* family members. **(D-G)** Schematic of TPZ Sox2 expression in WT and different *her* combined mutant larvae. Cells (NSCs) with higher Sox2^high^ levels are shown in dark magenta, and NPCs with lower Sox2^low^ levels in light magenta. Cells with high Sox2 levels (and *her6* levels) that are only present in a more lateral and ventral region close to the lateral wall of the third ventricle are shown as magenta cells labelled “V”. Black lines indicate the ventricular surface. **(D)** Schematic distribution of Sox2 in WT larvae. **(E)** Sox2 expression appears unchanged in the *Df(her4;12), Df(her2;15)* double deficiency mutants. **(F)** *her6, her9* double mutants lack Sox2^low^ expressing early neurogenesis progenitors and lack ventrally located Sox2^high^ cells in the lateral walls of the third ventricle. **(G)** *her*UDM larvae, in addition to the *her6, her9* double mutant phenotype, show Sox2^high^ nuclei scatter away from the ventricular surface. The dotted line suggests that the integrity of the ventricle is impaired. **(H)** Postulated *her* gene network interactions. Boxes show analyzed genes and gene expression. Lines indicate how mutations or overexpression influence expression of other genes, interactions may not be direct. Black lines indicate which *her* genes may compensate for other *her* gene activity (dot indicates directionality). The dashed black line indicates genes that only partially rescue loss of *her6* and *her9*. Green inhibition signs illustrate the autoregulation of *her6* and *her9*, and postulated negative autoregulation in the *her4/her12* and *her15/her2* clusters. Blue influence signs show changed gene expression of Sox2 and *neurog1* in *her6, her9* double mutants. The blue dashed influence sign indicates that loss of Notch-dependent *her* activity in *Df(her4;12), Df(her2;15)* only affects Sox2 expression when combined with the absence of *her6* and *her9* activity as detected in *her*UDMs. Red inhibition signs denote the downregulation of other genes upon overexpression of *her6* or *her4.* The dashed red inhibition signal indicates only mild downregulation of *ascl1b* by *her4* overexpression. Abbreviations: A, anterior; P, posterior; L, left; R, right; D, dorsal; V, ventral; cth, caudal thalamus; ptec, pretectum; pth, prethalamus; rth, rostral thalamus; tel, telencephalon; ve, ventricle; ZLI, zona limitans intrathalamica.

The role of *her4* during neural plate stages was reported to be similar to Notch-dependent *Drosophila E(spl)* genes, as both mediate repression of neurogenesis in the neuroepithelium by lateral inhibition (Nakao and Campos-Ortega, 1996; Takke et al., 1999). We find that in the larval TPZ, *her4* and *her15* continue to depend on Notch signaling and are largely co-expressed. In contrast to the neural plate, where *her4* and *her15* are expressed in domains distinct from *her9* (Stigloher et al., 2008), in the larval TPZ, *her4* and *her15* are co-expressed with *her9*, but expressed in domains distinct from *her6*. A similar dichotomy was observed for the expression of Notch-independent *hairy* versus Notch-dependent *E(spl)* genes in the fruit fly (Fisher and Caudy, 1998; Nakao and Campos-Ortega, 1996).

In the zebrafish neural plate, prepatterning genes mark the inter-proneural domains in a Notch-independent manner (Stigloher et al., 2008). For example, *her3* is expressed in two elongated stripes along the anteroposterior axis, flanked by *neurog1* positive progenitor pools (Bae et al., 2000), and *her5* inhibits neurogenesis in the intervening zone at the prospective midbrain-hindbrain boundary (Geling et al., 2004; Ninkovic et al., 2005). These homologs of the *Drosophila hairy* gene appear to act by positioning the proneural domains in the neuroectoderm independent of Notch signaling and are in that respect similar to Drosophila *hairy* function (Fisher and Caudy, 1998; Geling et al., 2004). It is therefore tempting to speculate that *her6* and *her9* execute prepattern information to maintain NSCs, and thus might have similar functions as *her3* and *her5* in the neural plate.

It was shown that *her6* morphant embryos lose *shha* expression in the mid-diencephalic organizer, which was explained by a premature differentiation of organizer cells accompanied by the loss of the boundary-characteristic *shha* expression (Scholpp et al., 2009). A similar *Shh* phenotype was observed in *Hes1, Hes5* double mutant mice (Baek et al., 2006). In *her6* mutants, we find a severe reduction of *shha* ZLI expression, which is even more severe in *her6, her9* double mutants that lack *shha* expression along the ZLI lateral ventricular walls completely. Despite *her6* and *her9* being expressed in distinct cell populations, *her9* can thus partially compensate *her6* function in the ZLI. Thus, *her6* and *her9* may indeed, in a Notch-independent manner, execute patterning information to maintain ZLI NSCs and signaling activity, similar to *her5* at the zebrafish midbrain-hindbrain boundary (Geling et al., 2004). Analysis of *Hes1* in mice revealed a high expression level mode of *Hes1* function in neural boundaries and organizing centers (Baek et al., 2006), which resulted in a model of two distinct *Hes1* functions. In so-called boundaries, high levels of Hes1 constitutively represses neurogenesis, whereas in compartments, its expression oscillates and regulates progression of neurogenesis (Kageyama et al., 2007).

Instead of just two in mice, zebrafish have evolved two *Hes1* and nine *Hes5* homologous genes, which appear to differ more strictly in their Notch-dependency. We used a genetic approach to identify differences in Notch-dependent versus -independent *her* gene functions, as well as potential redundancies in the *her* gene network (Fig.11H). Redundancy of *her* genes in zebrafish has been shown for *her1, her7* and *her13.2* in somitogenesis (Henry et al., 2002; Oates and Ho, 2002; Sieger et al., 2006), as well as for *Hes1* and *Hes5* in mice (Hirata et al., 2001) and *her5* and *her11/him* in zebrafish (Ninkovic et al., 2005) at the mid-hindbrain boundary. However, a detailed analysis of *her* gene interactions and redundancies, as well as a comprehensive understanding of Notch-dependent versus -independent activities has not been obtained.

*her6* and *her9* show negative feedback autoregulation and crossregulation, the latter however largely masked in the presence of functional feedback autoregulation, such that the expression of both *her* genes is most strongly upregulated in their double mutants. Due to the cluster deletions, we could not investigate autoregulation of Notch-dependent *her* genes in mutants, however *her4* overexpression reveals negative cross-regulation of *her15.* Overexpression of *her4* did not affect *her6* and *her9* expression, confirming that these genes are indeed completely independent of Notch signaling. Overexpression of *her6* repressed *her9, her4* and *her15*, revealing that Notch-independent *her6* activity is in a sense dominant over these *her* genes. Accordingly, *her6* and *her9* expression is largely unaffected in *Df(her4;12)*, *Df(her2;15)* double mutants. We observed alterations in *her4* and *her15* expression in *her6, her9* double mutants, however given that these double mutants affected NSCs composition of the TPZ, the effects at 3 dpf would not necessarily reflect direct regulation. The observed interactions and redundancies are outlined in Fig. 11H.

The loss- and gain-of-function experiments also revealed differential contributions of Notch-dependent versus -independent *her* genes to distinct types of NSCs and NPCs in the TPZ. We find that Notch-independent *her6* and *her9* are together critical for a subset of Sox2^high^ NSCs at the lateral ventricular wall of the TPZ, as well as of Sox2^low^ NPCs. This is consistent with an interpretation that Notch-independent *her6* and *her9* activity would be responsible for long-term maintenance of TPZ NSCs, tightly linked to the activity of the ZLI. *her6, her9* double mutants also display a TPZ phenotype as well as severe malformations in eye development (data not shown), similar to those reported for *Hes1* mutant mice (Nakamura et al., 2008; Nakamura et al., 2000; Tomita et al., 1996).

We were surprised to find that in *Df(her4;12)* or *Df(her2;15)* mutants, as well as in double deficient larvae lacking all 9 zebrafish Notch-dependent *Hes5* homologs, the TPZ appeared morphologically largely normal, and developed an apparently normal NSC and NPC composition of the TPZ (Fig. 11D-F). This suggests that in zebrafish, Notch-dependent *her* genes may have only a minor contribution to NSC maintenance. Combined deletion of the eleven zebrafish *Hes1* and *Hes5* homologs in *her*UDMs however, causes loss of most NSCs and essentially all NPCs in the TPZ, resulting in severe morphological malformations of the CNS and retina (Fig. 11G). The *her*UDM phenotype in severity appears similar to the severe *Hes1, Hes5* double mutant phenotype in mice with premature differentiation and stem cell depletion (Baek et al., 2006; Hatakeyama et al., 2004; Ohtsuka et al., 1999). The *her*UDM mutant phenotype is more severe than the *her6, her9* double mutant phenotype, revealing that indeed Notch-dependent *her* genes also contribute to NSC maintenance. Nevertheless, *her6* functions most prominently in this context, as revealed by *her*UDMs rescue by a single WT allele of *her6.* In summary both, Notch-dependent and - independent *her* genes act together to regulate NSC maintenance in the TPZ, with the most prominent activity provided by *her6.*

We note that even in the *her*UDMs, proliferating NSCs can be observed at 3 dpf. NSCs in *her*UDMs may be maintained into 3 dpf by additional *her* genes which have their strongest expression early in development, but might still provide some *her* activity to the TPZ, such as *her8a* or *her8.2, Hes6* homologs, which we show have both Notch-dependent and -independent aspects of expression in the 2 dpf and at very low levels 3 dpf TPZ.

Notch-dependent *Hes/her* genes have been demonstrated to be important regulators of neurogenesis (Nakao and Campos-Ortega, 1996; Takke et al., 1999). Surprisingly, in *Df(her4;12), Df(her2;15)* double deficiency mutants, expression of *ascl1a/b, neurog1* and *olig2* in the TPZ appears largely normal. Thus, in the TPZ, these nine Notch-dependent *her* genes together appear not essential for the initiation of NPCs and proneural gene expression. Our *her4* overexpression experiments support this interpretation: in *her4* heat-shocked embryos, expression of *neurog1* and *neurod1/6b* are largely normal, and of *ascl1b* in most embryos only slightly reduced. In contrast, loss of Notch-independent *her6* and *her9* results in smaller and disorganized domains of *ascl1a/b, neurog1* and *olig2* in the TPZ. Overexpression of *her6* eliminates *neurog1* and *ascl1b* expression, indicating loss of NPCs, and causes a reduction of *neurod1/d6b* expression, revealing reduction of NPCs entering neuronal differentiation. Our findings resemble results in mice, where *Hes5* is upregulated in *Hes1* mutants, but *Hes5* does not regulate *Mash1* (Ishibashi et al., 1995). Similar to our observations with *her6*, *Hes1* single and *Hes1, Hes5* double mutants have much stronger neurogenesis phenotypes than *Hes5* mutants (Hatakeyama et al., 2004).

In summary, we were able to show that the expanded number of *Hes1/Hes5* homologous genes in zebrafish goes along with a clear separation of Notch-dependency for different *her* genes, and enables us to differentiate Notch-dependent and -independent *her* activities in NSCs and NPCs. Our genetic analysis demonstrates that the *her* gene network is largely redundant within the Notch-dependent and -independent *her* groups, and that Notch-independent *her* genes may better substitute loss of Notch-dependent genes than vice versa. The observed robustness of the network is supported by several regulatory feedback loops and crossregulation (Fig. 11H). In an intact TPZ, Notch-independent *her* genes predominantly regulate NSC maintenance and NSC transition into the NPC pool, while Notch-dependent *her* genes may predominantly regulate progenitor dynamics and timing of differentiation, but not progenitor generation itself. This is in line with *Hes5* reported in mice to be responsible for regulation of the transition timing between phases of neurogenesis and gliogenesis, which is accompanied by changes in epigenetic regulation (Bansod et al., 2017). Within the TPZ, we identify two NSC populations: NSCtype1 depends predominantly on Notch-independent *her6* and may interpret patterning information at the ZLI (similar to “boundary” cells by Baek et al., 2006), while NSCtype2 expresses predominantly Notch-dependent *her4* and *her15.* It is tempting to speculate that NSCtype1 may serve long-term NSC maintenance, while NSCtype2 may generate NPCs. How both types, NSCtype2 *her4* and *shha* expressing ZLI organizer cells themselves, and adjacent *her6* expressing NSCtype1, work together to maintain the organizer should help to understand the complexity of the ZLI.

## Materials and Methods

### Zebrafish maintenance and breeding

ABTL strain zebrafish were kept and bred under standard conditions (Westerfield, 2000). Experiments were performed in accordance with the German Animal Welfare Guidelines. All experiments were approved by the ethics committee for animal experiments at the Regierungspräsidium Freiburg.

Zebrafish embryos were obtained through natural breeding and embryos were raised in E3 medium (5 mM NaCl, 0.17 mM KCl, 0.33 mM CaCl_2_, 0.33 mM MgSO_4_, supplemented with 2 ppm Methylene blue). 0.2 mM N-Phenylthiourea (PTU, Sigma Aldrich) was added to the E3 medium to prevent pigmentation. Embryos were staged as previously described (Kimmel et al., 1995).

### Embryo fixation and storage

Embryos were incubated in E3 (containing 0.2 mM PTU) supplemented with 1x tricaine methanesulfonate (MS-222) and fixed in 4 % PFA for 4 hours at RT or overnight at 4 °C. Embryos were washed several times with PBST and dehydrated stepwise with 25 %, 50 %, and 75 % MeOH in PBST, and stored at −20 °C in 100 % MeOH.

### Genotyping

DNA was isolated from embryos or tail biopsies of fixed larvae. Samples were heated 5 min to 95 °C in 60 μl TE buffer, after addition of 3 μl Proteinase K (20 mg/ml) digested for at least 4 hrs at 55 °C. After heat-inactivation of Proteinase K for 10 min at 95 °C, 3 μl of the DNA sample was used in 30 μl PCR volume. The primer concentration was 0.3 μM unless stated otherwise. PCR programs are supplied in Supplementary Table 3. For genotyping of embryos used in the qPCR, an alkaline lysis was performed. Tails were cut and transferred to 40 μl 50 mM NaOH and incubated for 45 min at 95 °C. The lysis was stopped by addition of 4 μl 1 M Tris-HCl (pH 7.5).

### Cas9 mRNA and sgRNA generation

Cas9 mRNA was transcribed from the hspCas9 plasmid (Ansai and Kinoshita, 2014) and Cas9-nanos from the Cas9-nanos-3’-UTR plasmid (Moreno-Mateos et al., 2015) using the SP6 mMessage mMachine kit (Thermo Fischer), and a polyA tail was added (PolyA tailing kit from Thermo Fischer). sgRNAs were designed with CRISPRscan (Moreno-Mateos et al., 2015) and ordered as oligonucleotides to generate DNA templates (Gagnon et al., 2014). Annealing of the constant oligonucleotide with the variable oligonucleotide was done by cooling from 95 °C down to 25 °C in 1 hr in a PCR machine. Annealed oligonucleotides were filled with T4 DNA Polymerase (New England Biolabs) for 20 min at 11 °C. sgRNAs were transcribed 3-4 hrs at 37 °C using the MEGAscript T7 kit according to protocol (Thermo Fischer) and dissolved in 15 μl H_2_O.

### Targeted induction of knockouts and large deficiencies

#### *her6 m1358* mutant allele

Two sgRNAs (transcribed from p3-Oligo04 and p4-Oligo05) were used to generate the *her6* knockout allele *m1358.* The intragenic deletion generates a frameshift that truncates the protein after amino acid 36 before the bHLH domain. Lower case letters are deleted in the mutant allele *m1358*:

5’TATAGTGCTTCAAGAagtgattagacagattgactcattttgtttttattttgcattttagtcttctaaacccattatggagaaaagaagaagagcgagaatcaacgaaagcttgggtcagctgaaaacgttaatcttggatgctctgaaaaaagatgtaagtaccgaaagtccgactgagtctttcagttaggctatatctggctggaatattaatgcatttctttatatagccgcatatctaaatcagtctctttcttcactctcagagctccagacactctaaacttgagaaagccgacATCCTGGAGATGACA3’

#### *her9 m1368* mutant allele

Two sgRNAs (transcribed from p186-Oligo26 and p187-Oligo27) were used to generate the *her9* knockout allele *m1368.* The intragenic deletion generates a frameshift that truncates the protein after amino acid 18 before the bHLH domain. Lower case letters are deleted in the mutant allele *m1368*:

5’ATTGCTGGTGCCCCTgccagtggatctcatactcctgacaagccaaagaatgccagcgagcatagaaagtcttcaaagccaatcatggaaaagcgccgcagagcgagaatcaacgagagccttgggcagctgaagactctcattcttgatgctcttaaaaaagatagctccagacactctaaattggagaaagctgatattctggagatgacagtcaagcacctgcgcaatttacaacgtgttcagatgagcgcagccttgtcagctgacacaaacgtcctcagcaagtaccgcgcaggattcaacgagtgcatgaacgaggtgactcgatttctctctacctgcgagggagtgaatacagaggtcagatcgcgacttcttaaccacctgtCCGGTTGTATGGGAC3’

#### Df(her2;15) m1490

Two sgRNAs (transcribed from p211-Oligo38 and p219-Oligo43) were used to generate the 30 kb deficiency *Df(Chr11:her2,her15.1,her15.2)*, which we abbreviate to *Df(her2;15)* (Fig. S10). 1 nl of the injection mix (6.5 μl total volume), which consisted of 1 μl of each sgRNA (transcribed from Oligo p211 and Oligo p43) plus 4.5 μl Cas9-nanos mRNA (500-800 ng/μl), was injected into 1 cell stage zebrafish embryos. 24 hpf single embryos were genotyped by PCR with two primers p213 + p222 (0.3 μM each) to test for knockout events (Supplementary Table 1). The PCR was performed using MyTaq polymerase (Bioline; PCR program in Supplementary Table 3). If embryos showed the desired deletions, siblings were raised and crossed to ABTL fish. F1 embryos carrying deletions were identified by PCR and siblings were raised to adulthood. Adult F1 fish were fin clipped and genotyped with primers p213 + p222 (0.3 μM each) + p220 (0.08 μM) (Supplementary Tables 1 and 3). Stable heterozygous F2 lines were established and appear to grow normally.

#### Df(her4;12) m1364 and m1365

Two sgRNAs (transcribed from p39-Oligo12 and p47-Oligo16) were used to generate the 28 kb deficiency *Df(Chr23:her12,her4.1,her4.2,her4.3,her4.4,her4.5)*, which we abbreviate to *Df(her4;12) (m1364* and *m1365* are shown in Fig. S10). 1 nl of the injection mix (8.3 μl total volume), which consisted of 1 μl of each sgRNA (transcribed from Oligo12 and Oligo16) plus 6 μl hspCas9 mRNA (600-1200 ng/μl), was injected into 1 cell stage zebrafish embryos. 24 hpf single embryos were genotyped by PCR with p42 + p51 (0.3 μM each) to test for knockout events (Supplementary Table 1). The PCR was performed using MyTaq polymerase (Bioline; PCR program in Supplementary Table 3). If embryos showed deletions, siblings were raised and crossed to ABTL fish. F1 embryos were identified by PCR and siblings were raised to adulthood. Adult F1 fish were fin clipped and genotyped with primers p42 + p43 + p51 (0.3 μM each) (Supplementary Tables 1 and 3). Stable heterozygous F2 lines were established and appear to grow normally.

#### *her* activity depleted *Df(her2;15), Df(her4;12), her4, her6* (*her*UDM) strain

The *her9* gene (Chromosome 23: 23,399,520-23,401,305) is only 1.9 Mb away from *her4.5* (Chromosome 23: 21,469,203-21,471,022), which is its closest neighbor in the *her4* gene cluster. To obtain the combined *her6*, *her9*, *her4;12* cluster and *her2;15* cluster mutant strain (abbreviated: *her*UDM), *Df(her4;12) m1364* heterozygous embryos were injected with sgRNAs (transcribed from p186-Oligo26 and p187-Oligo27) to target *her9* on the *Df(her4;12)* chromosome. For this mutant, 1 nl of the injection mix (5 μl total volume), which consisted of 1 μl of each sgRNA and 3 μl Cas9-nanos mRNA (500-800 ng/μl), was injected into 1 cell stage zebrafish embryos. The establishment of stable F2 lines was similar as described for the large deficiency mutants. The PCRs were performed with primer p197 and p198 and the program can be found in Supplementary Table 3. The allele was named *her9 m1505.* Lower case letters are deleted in the mutant:

5’ TGCTGGTGCCCCTGCcagtggatctcatactcctgacaagccaaagaatgccagcgagcatagaaaggtaaaaatcacttataatgcgattgcatattgttttagaagaatgacagctgcatcagttattctaaaaaaagagagaatgtcttttaattacgtaaattaattaattttccatttcatctgcagtcttcaaagccaatcatggaaaagcgccgcagagcgagaatcaacgagagccttgggcagctgaagactctcattcttgatgctcttaaaaaagatgtaagttttatctacaactcttgtcatgcttcagtcacccgacgttagtaattctgaaacagtttctaaccgaattctgctcattcacagagctccagacactctaaattggagaaagctgatattctggagatgacagtcaagcacctgcgcaatttacaacgtgttcagatgagcggtaagttgcaagtcagattcctcaagatgataaacttttaacgtgcttttaaaaacgcaatttaatttcctgaatacacaatctaaatacgttttatctttgtttaccagcagccttgtcagctgacacaaacgtcctcagcaagtaccgcgcaggattcaacgagtgcatgaacgaggtgactcgatttctctctacctgcgagggagtgaatacagaggtcagatcgcgacttcttaaccacctGTCCGGTTGTATGGG3’

When breeding the *her9 m1505, Df(her4;12)* chromosome, we also checked for unwanted crossing over events by PCR for the *Df(her4;12)* allele.

### LY-411575 treatment

For pharmacological inhibition of Notch signaling by LY-411575 (Rothenaigner et al., 2011), embryos were incubated in E3 supplemented with 0.2 mM PTU until 64 hpf. Then embryos were incubated in 1x E3 supplemented with 0.2 mM PTU, 2 % DMSO, 10 μM LY-411575 for 8 hrs. Control embryos were treated in 1x E3 supplemented with 0.2 mM PTU and 2 % DMSO for 8 hrs in the same six-well plate as the LY treated embryos. Embryos were checked for vital functions and fixed at 72 hpf in 4 % PFA.

### NICD overexpression

For the overactivation of the Notch signaling pathway, *Tg(hsp:Gal4;UAS:RFP) kca4Tg* were crossed with *Tg(UAS:NICD)kca3Tg* (Scheer and Campos-Ortega, 1999).This allowed heat shock controlled overexpression of NICD. At 64 hpf embryos were heat shocked for 30 min at 40 °C in pre-heated E3 (with 0.2 mM PTU). Heat shocked embryos were then sorted for the heat shock induced RFP marker. Non-fluorescent embryos were used as negative controls. Embryos were tested for the *UAS:NICD* element by PCR (Supplementary Tables 1 and 3).

### Whole mount immunofluorescence

Fixed and stored embryos were rehydrated with 75 %, 50 %, 25 % MeOH in PBST and washed several times with PBST. Embryos were digested with Proteinase K (10 μg/ml) in PBST for 45 min at RT. Embryos were re-fixed with 4 % PFA for 20 min at RT and then washed several times with PBST. Larvae were blocked in blocking solution (1 % BSA, 5 % goat serum, 1 % DMSO in PBST) for 1 hr and incubated with the primary antibodies diluted in blocking solution. The anti-Sox2 antibody ([20G5], abcam, ab171380, LOT GR3253929-5) and the anti-phospho-Histone H3 (Ser10) antibody (Merck, #06-570) were diluted 1:400. Embryos were incubated in primary antibodies overnight at 4 °C and then washed several times for 30 min each with PBST supplemented with 1 % DMSO. Embryos were incubated with the secondary antibodies 1:1000 in PBST supplemented with 1 % DMSO and 1 % blocking reagent (Roche, #1096176) overnight at 4 °C. Secondary antibodies were Alexa 555 goat anti-mouse IgG (Thermo Fisher Scientific, #A-11001) and Alexa 488 goat anti-rabbit IgG (Thermo Fisher Scientific, #A-11070). Embryos were washed several times in PBST, transferred to 80 % glycerol in PBST and stored at 4°C protected from light. The embryos were recorded soon after staining.

### RNA detection by Hybridization Chain Reaction (HCR)

Whole mount in situ hybridization by HCR was performed as described in the HCR v3.0 protocol for zebrafish embryo and larvae (Choi et al. (2018) and by Molecular Instruments. The following changes to the protocol were made: In preparation stage 11, 3 dpf larvae were treated with proteinase K (30 μg/ml) for 30 min. In detection stage 1, 15 larvae were used in 2 ml tubes. After amplification stage 6, embryos were washed twice in PBST for 5 min and stored in 80 % glycerol in PBS. Probes for *neurog1* (ENSDART00000078563.5), *asc1a* (ENSDART00000056005.5), *ascl1b* (ENSDART00000183550.1), *sox2* (ENSDART00000104493.5) and *her6* (ENSDART00000023613.9) were ordered and designed by Molecular Instruments.

We designed probe sets for *her4* (ENSDART00000079274.4), *her9* (ENSDART00000078936.4), *her15* (ENSDART00000055707.6) and *olig2* (ENSDART00000060006.5) (Supplementary Table 2). For example, the *olig2* probe set consists of ten probe pairs which are evenly distributed along the mRNA sequence (Supplementary Table 2). One probe pair consists of two gene specific 25 bp sequences each with a GC content of 37-85 % and a Tm of 55-77 °C. The two sequences were separated by a gap of two nucleotides. The B1 specific spacer and initiator sequences were added to the reverse complement of the two gene specific sequences. The complete probe set was ordered from Sigma Aldrich. Equal volumes of all 20 single probes were mixed and used as probe mixture containing 5 μM of each probe in detection stage 3.

### Generation of in situ hybridization probes

Specific *her* probes were designed to avoid cross-hybridization of conserved domains. For the generation of *her6, her9, her4.1-her4.5., her15.1-her15.2, her2* and *her12* whole-mount *in situ* hybridizations probes, sequences spanning the last exon and/or 3’ UTR of the respective genes were amplified by PCR (all primers for probe generation are given in Supplementary Table 1). The PCRs were performed on WT genomic DNA with PfuUltra II (Agilent). The 50 μl PCR mix contained 6-10 μl dNTP mix (2.5 mM each), 5 μl 10x Pfu buffer, 0.2 mM of each primer, 2 μl DNA and 1 μl PfuUltra II Polymerase. Before cloning, A-overhangs were added by incubating 20 μl of the purified PCR product with 9.7 μl MyTaq buffer, 0.2 μl MyTaq polymerase and 0.1 μl (25 μM) dATPs for 20 min at 72 °C. After amplification, the probes were cloned into a TOPO vector according to the manufacturer’s protocol (TOPO TA cloning kit, Thermo Fisher).

For the *her4* probe, which recognizes *her4.1* (ENSDART00000079274.4), *her4.2* (ENSDART00000137573.2), *her4.3* (ENSDART00000104209.4), *her4.4* ENSDART00000079265.6) and *her4.5* (ENSDART00000104206.4), a template was generated with primer p159F and p160R (annealing 57 °C, 40 sec). For the *her15* probe, which recognizes *her15.1* (ENSDART00000055707.6) and *her15.2* (ENSDART00000055706.6), a template was cloned using primers p157F and p158F (annealing 56 °C, 40 sec). The *her2* (ENSDART00000055709.5) probe template was amplified with p154F and p155R (annealing 56 °C, 40 sec) and the *her12* (ENSDART00000044080.7) probe template was amplified with p152F and p153R (annealing 56 °C, 40 sec). The *her6* (ENSDART00000023613.9) probe template was amplified using p147F and p148R (annealing 57 °C, 55 sec) and the *her9* (ENSDART00000078936.4) probe template was amplified with p149F and p150R (annealing 56 °C 40 sec). All primers are given in Supplementary Table1.

The *her8a* (ENSDART00000123395.4) probe template was amplified as published in Webb et al. (2011) with p322F and p327R (annealing 56 °C, 1 min) and the *her8.2* (ENSDART00000101578.4) probe template was amplified with p260F and p259R (annealing 56 °C, 1 min). For *her8a* and *her8.2* cDNA was used as a template in the PCR.

*sox2* (ENSDART00000104493.5), *neurod1* (ENSDART00000011837.6), *neurog1* (ENSDART00000078563.5), *shha* (ENSDART00000149395.3), *neurod6b* (ENSDART00000185805.1), *irx1b* (ENSDART00000079114.6; (Scholpp et al., 2009), *ascl1a* (ENSDART00000056005.5; Allende and Weinberg (1994), *ascl1b* (ENSDART00000183550.1; Allende and Weinberg (1994) probes have been previously published (www.zfin.org) and plasmids were validated by sequencing.

Plasmids were linearized and transcribed with T7 or SP6 RNA Polymerases (Thermo Fischer). Antisense RNA probes were generated using the DIG or DNP based RNA labelling kits from Roche (Merck, Germany). The quality of the RNA probe was checked by agarose gel electrophoresis.

### Whole-mount in situ hybridizations

Chromogenic whole mount in situ hybridizations were performed as previously described (Holzschuh et al., 2003). After fixation, storage and rehydration, embryos were washed three times in PBST and treated with 10 μg/ml Proteinase K (AppliChem, 15 min incubation per 24 hrs of development). Embryos were washed with PBST and fixed in 4% PFA for 20 min at room temperature. Embryos were washed 5 times in PBST and incubated in hybridization mix (50% formamide, 5x SSC, 5 mg/ml torula yeast RNA type IV, 50 μg/ml heparin, 0.1% Tween-20) at 65 °C for 4 hrs. Next, embryos were incubated with digoxigenin labelled antisense RNA probes in hybridization mix (dilution 1:200 - 1:500) at 65 °C overnight. After hybridization, embryos were washed at 65 °C for 20 min each with the following solutions: twice with 50% formamide in 2x SSCT, twice with 25% formamide in 2x SSCT for 20 min each, twice in 2x SSCT for 20 min each, three times in 0.2x SSCT for 20 min each. Next, embryos were washed once in 0.1x SSCT in 0.5x PBST for 10 min and twice in PBST for 10 min each at room temperature. Embryos were blocked for 2 to 3 hrs with 2 % heat inactivated goat serum (Vector Laboratories) and 4 mg/ml BSA (AppliChem) in PBST. Embryos were incubated overnight at 4 °C in blocking solution supplemented with the alkaline phosphatase coupled anti-digoxigenin antibody (Roche, REF 11093274910) 1:3000. Embryos were washed six times in PBST at room temperature for 20 min each and three times in NTMT (100 mM NaCl, 100 mM Tris-HCl at pH 9.5, 50 mM MgCl2, 0.1 % Tween-20) for 15 min each. For the staining reaction, 0.18 mg/ml BCIP (5-bromo-4-chloro-3-indolyl-phosphate, AppliChem) and 0.45 mg/ml NBT (4-nitroblue tetrazolium chloride, AppliChem) were added to NTMT. Staining was performed in the dark and stopped by 3 washes in PBST supplemented with 0.1 mM EDTA. For longer storage, embryos were fixed in 4 % PFA for 1 hr at room temperature. After three more washes in PBST, embryos were transferred to 80% glycerol in PBST supplemented with 0.1 mM EDTA and stored at 4 °C protected from light.

### Fluorescent whole-mount in situ hybridizations

Fluorescent in situ hybridizations were performed as described in the section whole mount in situ hybridization until addition of probes. The digoxigenin and dinitrophenol labelled antisense probes (see “Generation of in situ hybridization probes”) together were added to the hybridization mix and the embryos were incubated at 65 °C overnight. Washes were performed at 65 °C as described above. Embryos were washed in TNT (100 mM Tris-HCl at pH 7.5, 150 mM NaCl, 0.5 % Tween-20) and blocked in TNT supplemented with 1 % Boehringer Block (Roche) for 2 to 3 hrs at room temperature. The anti-digoxigenin-POD Fab fragment antibody (Roche, REF 11207733910) was used 1:400 in blocking solution and embryos were incubated overnight at 4 °C. Embryos were washed eight times 15 min each in TNT at room temperature, rinsed in 100 mM borate buffer pH 8.0 and incubated in staining mix containing 2 % dextrane sulfate, 1% tyramide stock solution Alexa Fluor 488 (Thermo Fischer), 0.0015 % H2O2 and 112.5 μg/ml 4-iodphenole in 100 mM borate buffer pH 8.0 for 1 hr at room temperature. Embryos were washed three times in TNT, incubated in 0.3 % H2O2 in TNT for 30 min and then washed five times in TNT for five minutes each at room temperature. Embryos were blocked in 1% blocking reagent (Roche) in 100 mM maleic acid, 150 mM NaCl at pH 7.5 for 2 to 3 hrs at room temperature. The anti-DNP HRP antibody (Perkin Elmer, FP1129) was used 1:200 in blocking solution and embryos were incubated overnight at 4 °C. Embryos were washed eight times 15 min each in TNT at room temperature, rinsed in 100 mM borate buffer pH 8.0 and incubated in staining mix containing 2 % dextrane sulfate, 1 % tyramide stock solution Alexa Fluor 555 (Thermo Fischer), 0.0015 % H2O2 and 112.5 μg/ml 4-iodphenole in 100 mM borate buffer pH 8.0 for 1 hr at room temperature. Embryos were washed three times in TNT and three times in PBST for five minutes each at room temperature. Embryos were transferred to 80 % glycerol in PBS and stored at 4 °C protected from light.

### Microscopy

Imaging of chromogenic whole mount in situ hybridizations was performed with the Zeiss Axioskop2, Axiovision SE64, Rel.4.9.1 with AxioCam ICc1 with a Plan-Neofluar 20×/0.5 or 10×/0.3 objective and DIC optics. Embryos were mounted in 80 % glycerol in PBST supplemented with 0.1 mM EDTA or in 80 % glycerol in H2O supplemented with 1 % standard agarose (Bioron). All shown images are single z-planes from an image stack.

Imaging of fluorescent stainings was performed with an inverted microscope with LSM-880 Airyscan (Zeiss). A 40x objective LD LCI Plan Apochromat 40x/1.2 autocorr (Zeiss) was used. Embryos were embedded in 80% glycerol in H2O supplemented with 1 % standard agarose (Bioron). The immersion medium was glycerol. The imaging program ZEN 2.3 SP1 FP3 (black) version 14.0.22.201 was used. If Airyscan was used, image stacks were processed with the ZEN black software using the Airyscan algorithm (v. 2.3 SP1 Zeiss). All images shown are single z-planes from an image stack, except for Fig. 2A,B,C; Fig. 4A,B,C,C”, D, D”; Fig. 7A,B,C,C”,D, D”, Fig. 10C,D, E,F and Fig. S3A,B, which show orthogonal projections of an image stack. Fig. 2D, D’ and D” show single z planes from an image stack of laterally mounted embryos.

Figures were assembled in Photoshop. Image levels were linearly adjusted except for Fig. 5B-C’, F-G’, Fig. S8 and Fig. S9A-F’ to improve visibility in very dark stained areas.

### Numbers of embryos analyzed

For WISH experiments analyzing WT, compound treated or heat shock overexpression treated embryos, shown in Fig. 1G-R’, Fig 8, Fig. 9, Fig. S1, Fig. S2 G-J’ and Fig. S11, for each condition 10-20 larvae were examined for their expression patterns and representative images are shown.

When the (n) numbers are provided in images or legends, the number indicates how many larvae were imaged. If staining patterns or intensity varied within one experimental condition (Fig. 9B’, E’ and J’), one embryo with a staining pattern representative for the majority of embryos analyzed is shown, and the numbers in the bottom right corner indicate the number of embryos with the representative expression pattern and the total number of embryos analyzed for this condition.

For analysis of genetic mutants or fluorescent WISH shown in Fig. 1A-F’, Fig. 2, Fig. 3C-N’, Fig 4, Fig. 5, Fig. 6C-F’, Fig. 7, Fig. 10, Fig.S2A-D’, Fig. S3, Fig. S8, Fig. S9, the numbers (n) of embryos imaged for each condition are provided in the Figure legends or in the image panels, and in Supplementary Table 4.

### qPCR

The tail of each 4 dpf larva was cut and transferred to a new tube for genotyping as described above. The rest of each larva was stored individually in 75 μl RNA later solution (Ambion, AM7024) on ice. Following genotype detection by PCR, total RNA was extracted from 2-3 embryos pooled per genotype using the RNeasy Mini Kit (QIAGEN, 74106) and QIAshredder columns (QIAGEN, 79656). The sample was homogenized in 1 % β-mercaptoethanol in 600 μl RLT buffer (QUIAGEN, 74106). The suspension was transferred to a QIAshredder column and centrifuged for 1 min at 10,000 g. 700 μl 70 % ethanol was added to the flow through and transferred to RNeasy Mini spin columns. After centrifugation (15 s, 10,000 g), 350 μl RW1 buffer (QUIAGEN, 74106) was added to the membrane and centrifuged. A digestion step with DNase I was performed by adding 80 μl of the DNase I incubation mix (consisting of 10 μl DNase I stock solution with a concentration of 2.73 Kunitz units/μl and 70μl RDD buffer; QUIAGEN, 74106) to the RNeasy spin column membrane for 15 min on the benchtop to remove DNA from the membrane. After two more washing steps according to manufacturer’s protocol, the column was dried by centrifugation for additional 2 min. Total RNA was eluted by adding 30 μl H2O to the filter membrane. Tubes were incubated for 1 min at RT and centrifuged for 1 min at 10,000 g. The RNA concentrations were determined by NanoDrop measurements.

100 ng total RNA was used as a template for reverse transcription into cDNA. cDNA was generated with the SuperScript III RT Kit according to manufacturer’s protocol using oligo(dT)12-18 (0.5 μg/μl) primers. The qPCR was performed with the SYBR Green Supermix (Sso Advanced Universal, Biorad). The 10 μl qPCR mix contained 5 μl 2x SYBR Green Supermix, 1 μl primer mix (0.5 μM final concentration of each primer) and 2 μl cDNA. The PCR plate was carefully sealed and centrifuged for 10 min at 3000 rpm. qPCRs were performed with the LightCycler (Roche) and programs are given in Supplementary Table 3. The data analysis was performed as in Taylor et al. (2019), and *actb2* was used as reference gene. We note that for calculations of relative expression, we excluded one of the three biological replicates for the double mutant since the CT value for *actb2* control in this replicate was off. Therefore, the number of biological replicates for *her6* and *her9* qPCR in *her6, her9* double mutants is two. For all others the number of biological replicates is three.

## Acknowledgements

We thank Sylke Lange for technical assistance, the Life Imaging Center (LIC) for support in confocal imaging and image analysis, and Sabine Götter for excellent fish care. We thank Beatrice Weber for initial characterization of *her4* and *her6* mutant embryos, Masha Miranda Tolentino Voigt for initial characterization Notch-dependency of *her* genes, Samuel Wöhrle for initial characterization of *Tg(hsp:her6-FLAG)* embryos, Leonie Bohnert for establishing the HCR-RNA FISH technique in our lab, and Niklas Mayle for performing the WISH procedure for *shha* expression in *her6* and *her9* mutants. We thank Christian Altbürger for scientific discussion. The *neurod1* and *neurog1* plasmids were obtained from Patrick Blader. We thank Chaitanya Dingare for advice on designing the *olig2* HCR probe, and Rebecca Peters for help in designing Fig. 11H.

## Competing interests

The authors declare no competing or financial interests.

## Ethics Oversight

All animal experiments were approved by state authorities (Regierungspraesidium Freiburg permits 35-9185/G-16/123, 35-9185.81/G-19/19 and 35-9185.81/G-19/54).

## Data availability

All primary image data are available on request.

## Funding

This work was funded by grants from the Deutsche Forschungsgemeinschaft (DFG, German Research Foundation) under Germany’s Excellence Strategy – EXC-2189 CIBSS– Project ID: 390939984 (Gefördert durch die Deutsche Forschungsgemeinschaft (DFG) im Rahmen der Exzellenzstrategie des Bundes und der Länder – EXC-2189 – Projektnummer 390939984), the Excellence Initiative of the German Federal and State Governments (BIOSS - EXC 294), and CRC 850.

## Supplementary information

Supplementary Figures S1-S11 (one PDF file)

Supplementary Data Tables S1-S5 (one PDF file)

Supplementary Movies S1-S5 (separate mp4 files)

## Notes

### Competing Interest Statement

The authors have declared no competing interest.

### Summary of Updates

Update includes Supplemental Materials file

