## Supplemental Materials (11 Figures, 5Tables, Movie Legends) for "A network of Notch-dependent and -independent *her* genes controls neural stem and progenitor cells in the zebrafish thalamic proliferation zone"

**Supplementary Data:**

**Supplementary Figures S1-S11 (pages 2 – 15)**
**Legends for Supplementary Movies M1 – M5 (pages 16 – 17)**
**Supplementary Data Tables (following page 17)**

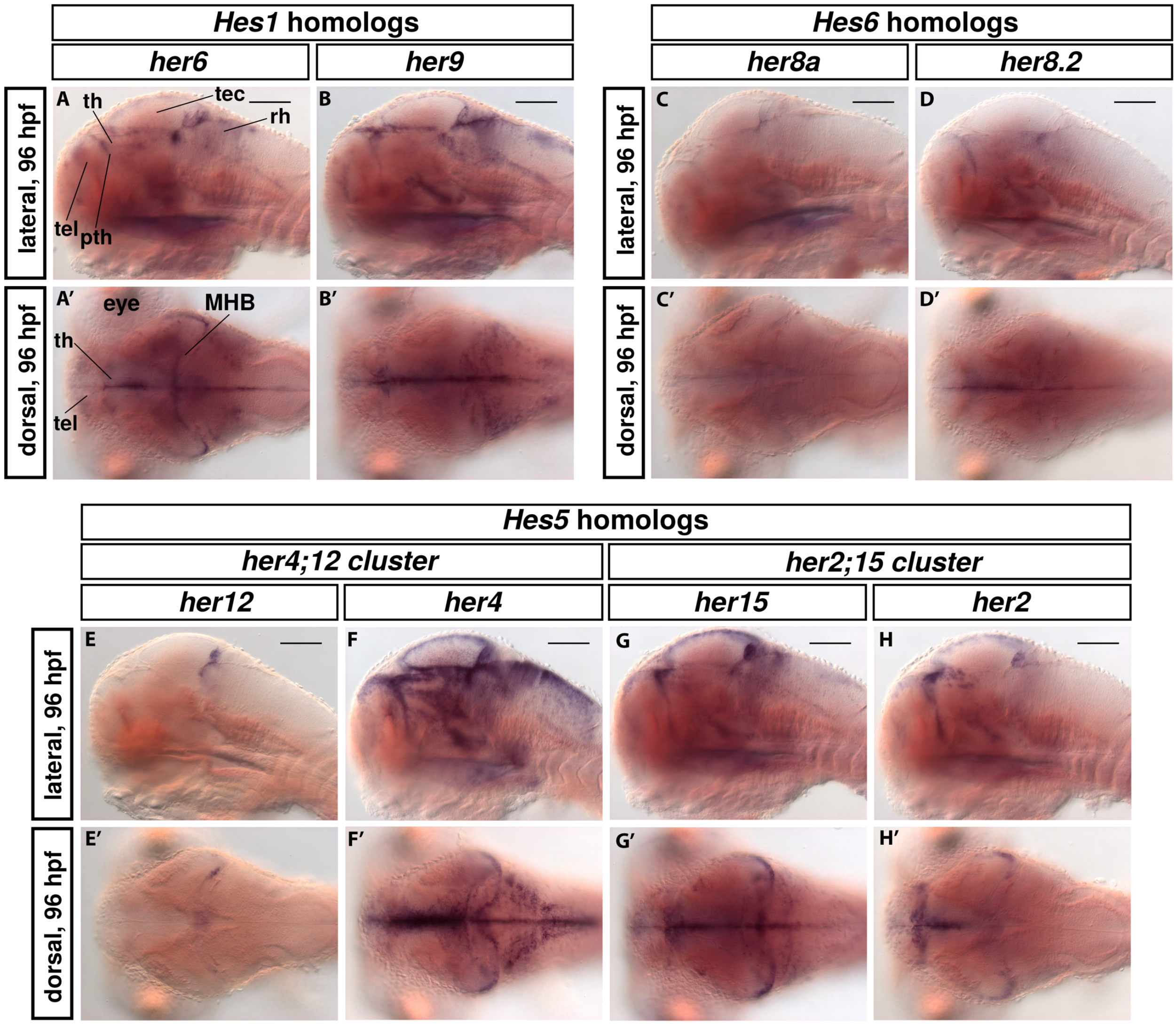

**Fig. S1. *her* gene expression in 4 dpf zebrafish larvae.**
**(A-H')** Expression of *her6*, *her9*, *her8a, her8.2, her12, her4, her15* and *her2* in 4 dpf zebrafish larvae visualized by WISH. Top rows show sagittal midline optical sections (A-H). Bottom rows show dorsal views of single horizontal planes at the level of the thalamus (A'-H'). **(A-B')** *Hes1* homologs. **(C-D')** *Hes6* homologs. **(E-H')** *Hes5* homologs. The *her4* probe detects *her4.1-her4.5* and the *her15* probe detects *her15.1* and *her15.2*. For each probe, three larvae were imaged and consistent WISH staining patterns observed. Abbreviations: MHB, midbrain-hindbrain boundary; pth, prethalamus; rh, rhombencephalon; tec, tectum; tel, telencephalon; th, thalamus proper. Scale bars, 100 μm.

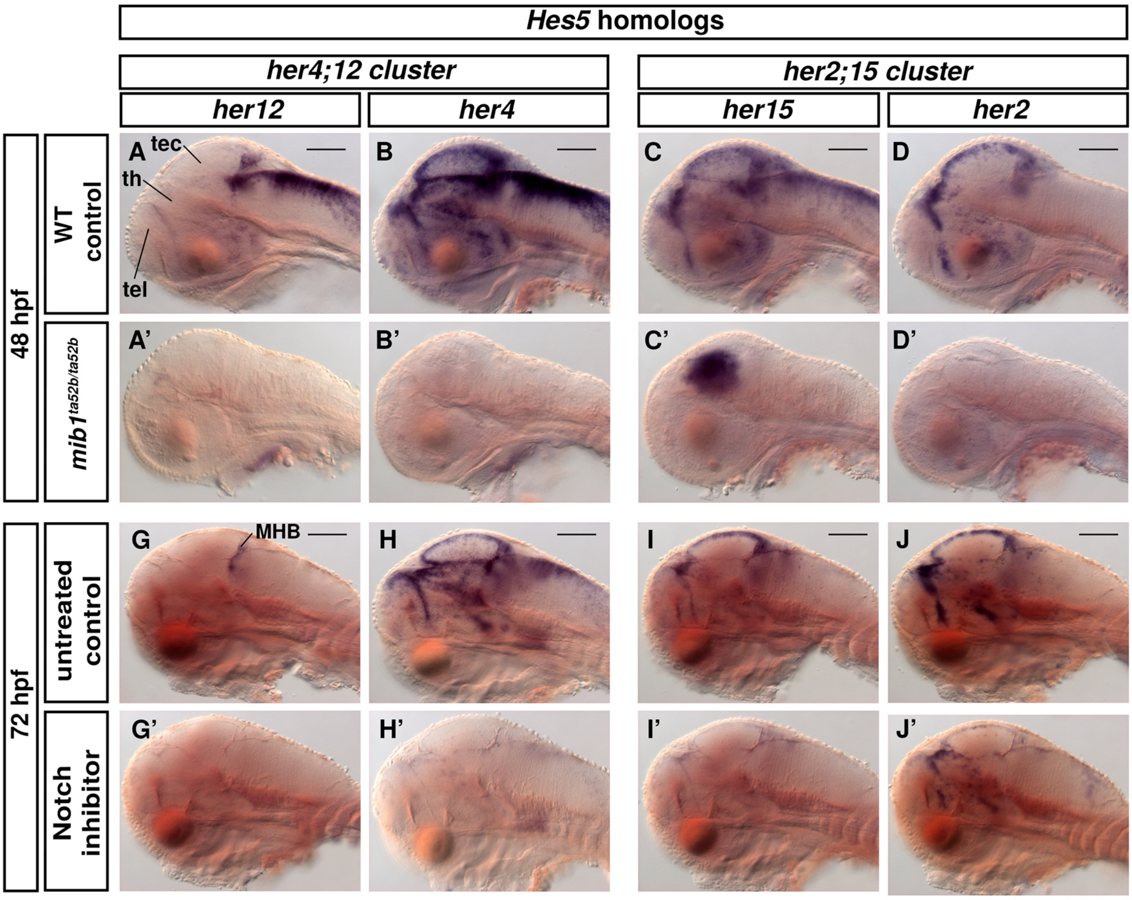

**Fig. S2. Expression of *Hes5* homologs after interference with Notch signaling.**
**(A-J')** Expression of *her12, her4, her15* and *her2* visualized by WISH, lateral views. **(A-D')** 48 hpf *mind bomb* mutants (*mib1*$^{ta52b}$) in comparison with control siblings (*mib1*$^{+/+}$ or *mib1*$^{+/ta52b}$). **(G-J')** *her* gene expression after LY-411575 Notch inhibition in comparison with DMSO treated controls. Larvae were treated for 8 hours with 10 µM LY-411575 or 2% DMSO, respectively, and fixed at 72 hpf. **(A-J')** Single sagittal midline image planes are shown. Three larvae per condition were imaged and consistent WISH staining patterns observed. The *her4* probe detects *her4.1-her4.5* and the *her15* probe detects *her15.1* and *her15.2*. Scale bars, 100 µm. Abbreviations: MHB, midbrain-hindbrain boundary; tec, tectum; tel, telencephalon; th, thalamus proper.

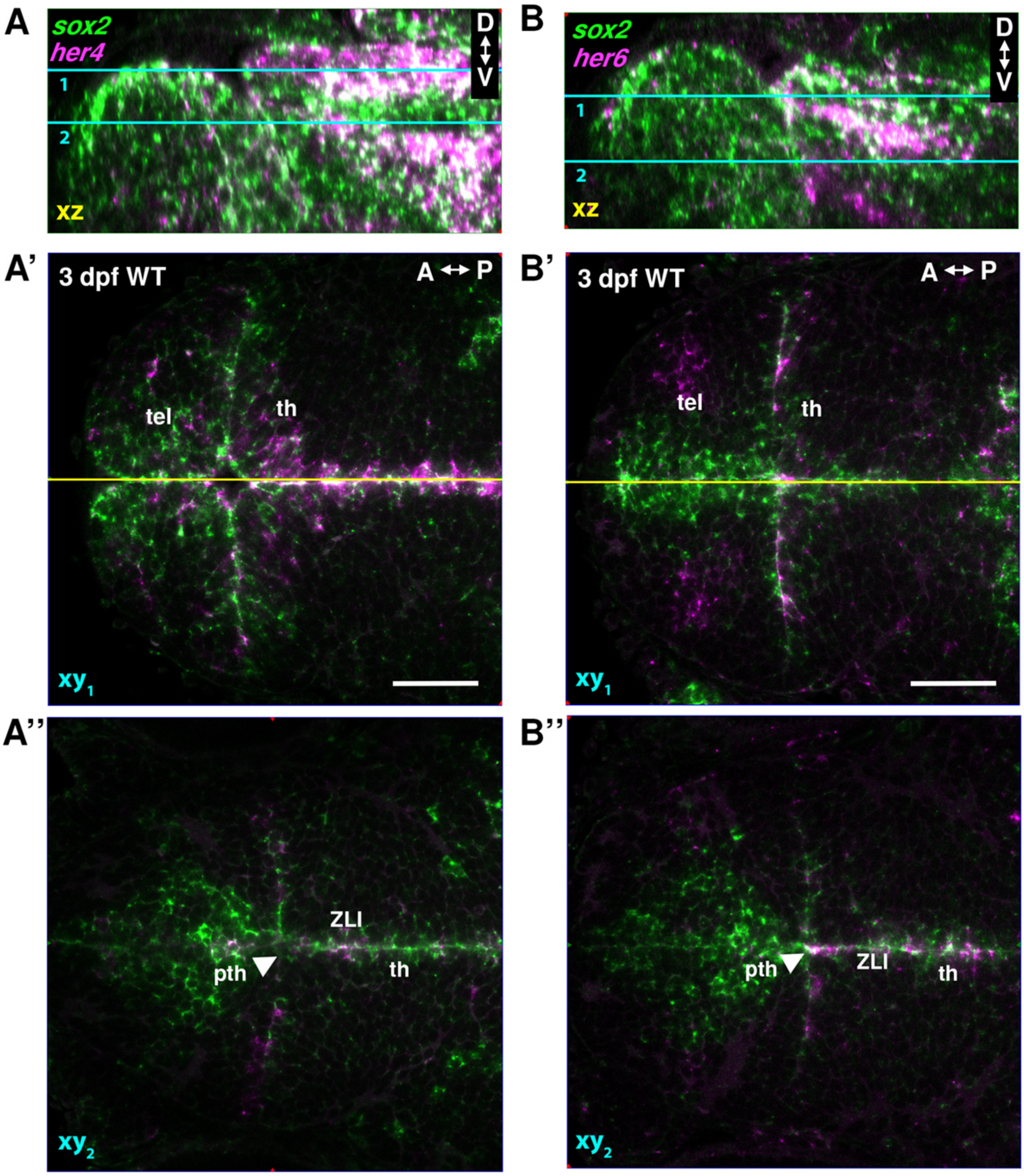

**Fig. S3. Expression analysis of *her4* and *her6* in relation to *sox2*.**
**(A-A’’)** Co-expression of *sox2* (green) and *her4* (magenta), and **(B-B’’)** Co-expression of *sox2* (green) and *her6* (magenta) visualized by double-fluorescent WISH. **(A, B)** Midsagittal planes (orthogonal reconstructions from dorsal view confocal stacks) with cyan lines indicating the horizontal confocal image planes 1 and 2 shown in A’, A’’ and B’, B’’. The yellow lines in A’ and B’ indicate the sagittal planes of A and B. A’ and B’ are dorsal views of the thalamus proper at the level of line 1 shown in cyan in A and B. A’’ and B’’ are dorsal views of the prethalamus at the level of line 2 shown in cyan in A and B. The *her4* probe detects *her4.1*-*her4.5* and the *her15* probe detects *her15.1* and *her15.2*. Scale bars, 50 µm. Numbers of embryos analyzed: A, n=3; B, n=3. Abbreviations: A-P indicates anterior-posterior, D-V dorsal-ventral orientation. pth, prethalamus; tel, telencephalon; th, thalamus proper; ZLI, zona limitans intrathalamica.

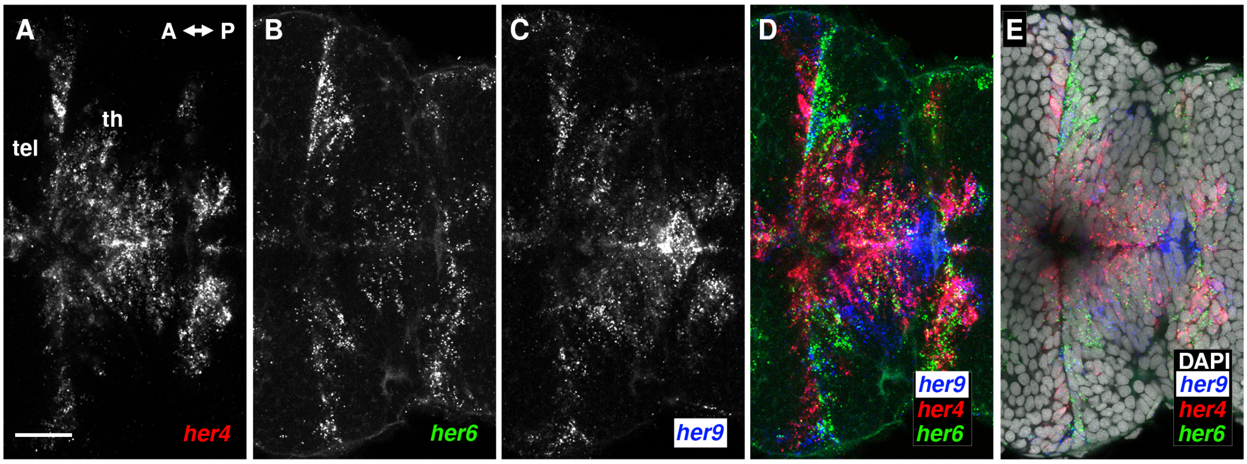

**Fig. S4 TPZ expression of *her4, her6* and *her9* in relation to each other.**
**(A-E)** *her4, her6* and *her9* co-detection by whole mount hybridization chain reaction (HCR). Single confocal plane with focus on the dorsal thalamus. A, *her4*; B, *her6*; C, *her9*; D, merge; E, merge with nuclear stain (DAPI). Two embryos were analyzed. Scale bar in A, 20 µm. Abbreviations: A-P indicates the anterior-posterior orientation; tel, telencephalon; th, thalamus proper.

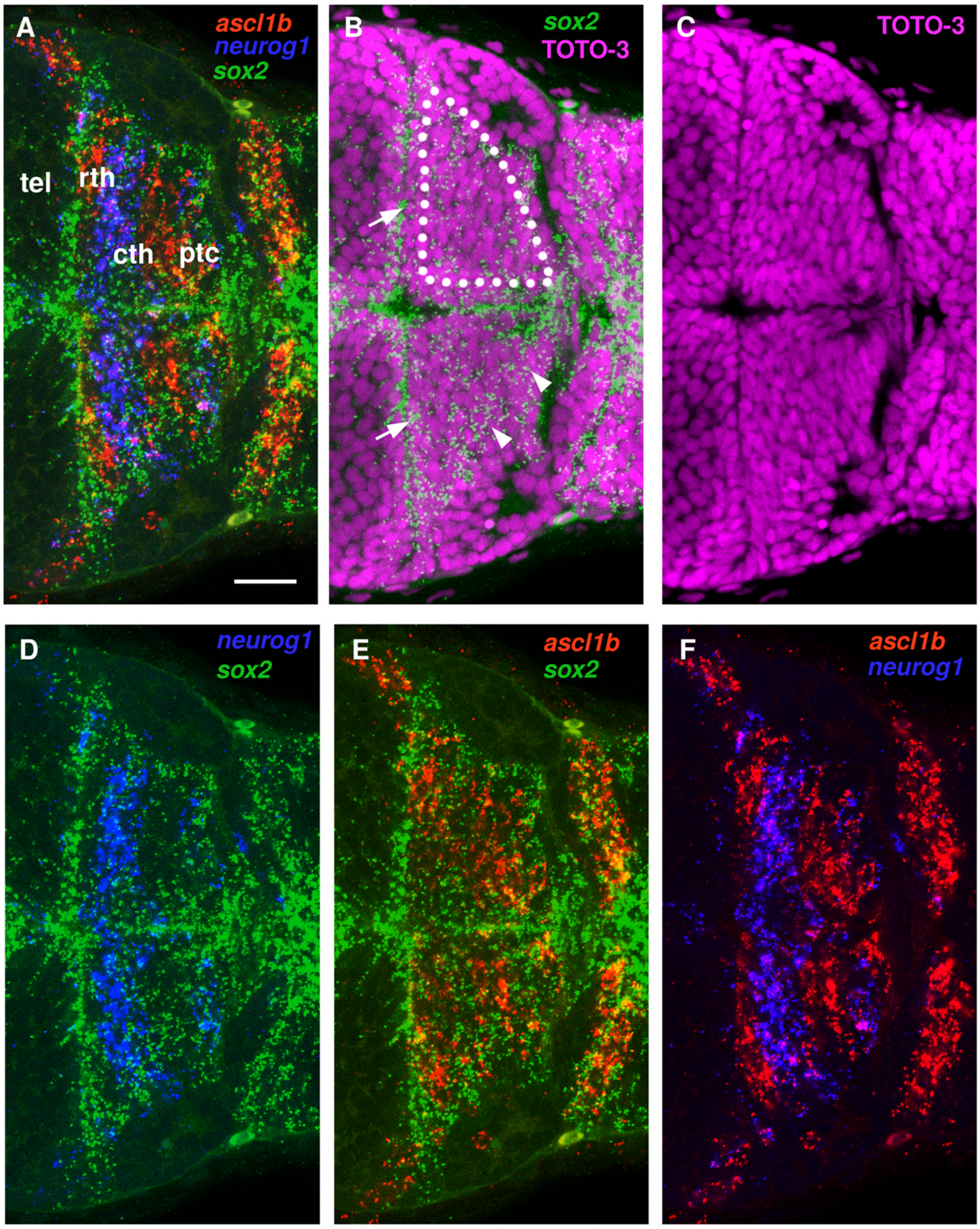

**Fig. S5 Analysis of NPC marker expression reveals *sox2*$^{low}$ expressing cells as progenitors.** **(A-F)** Triple probe HCR-RNA FISH showing *ascl1b* (red), *neurog1* (blue) and *sox2* (green) expression in the thalamus of a 3 dpf WT larva. **(A)** In *sox2$^{low}$* cells, *sox2* appears co-expressed with *neurog1* or *ascl1b*. **(B)** Nuclear stain with TOTO-3 reveals the location of nuclei relative to *sox2* mRNA. The dotted lines mark the area of *sox2*$^{low}$ expressing cells in the thalamus proper. Arrowheads mark *sox2*$^{low}$ expressing cells. Arrows mark ventricular *sox2*$^{high}$ expressing cells, with most *sox2* HCR signal apical to the nuclear layer at the ventricular surface. **(C)** The extent of the ventricular layer of nuclei (and cells) can be estimated from the TOTO-3 staining. **(D)** *neurog1* and *sox2* are co-expressed in *sox2$^{low}$* cells in the caudal thalamus. **(E)** *ascl1b* and *sox2* are co-expressed in *sox2$^{low}$* cells in the rostral thalamus. **(F)** *ascl1b* and *neurog1* are expressed at high levels in largely separated domains, but with partial overlap of low expression in the thalamus proper. Images A-F show the same embryo and horizontal confocal image plane as in Fig. 4C' with color channels linearly adjusted to emphasize distinct and overlapping expression domains (Supplementary Movie M1). Scale bar in A, 20 µm. Three embryos were analyzed. Abbreviations: cth, caudal thalamus; ptc, pretectum; rth, rostral thalamus; tel, telencephalon.

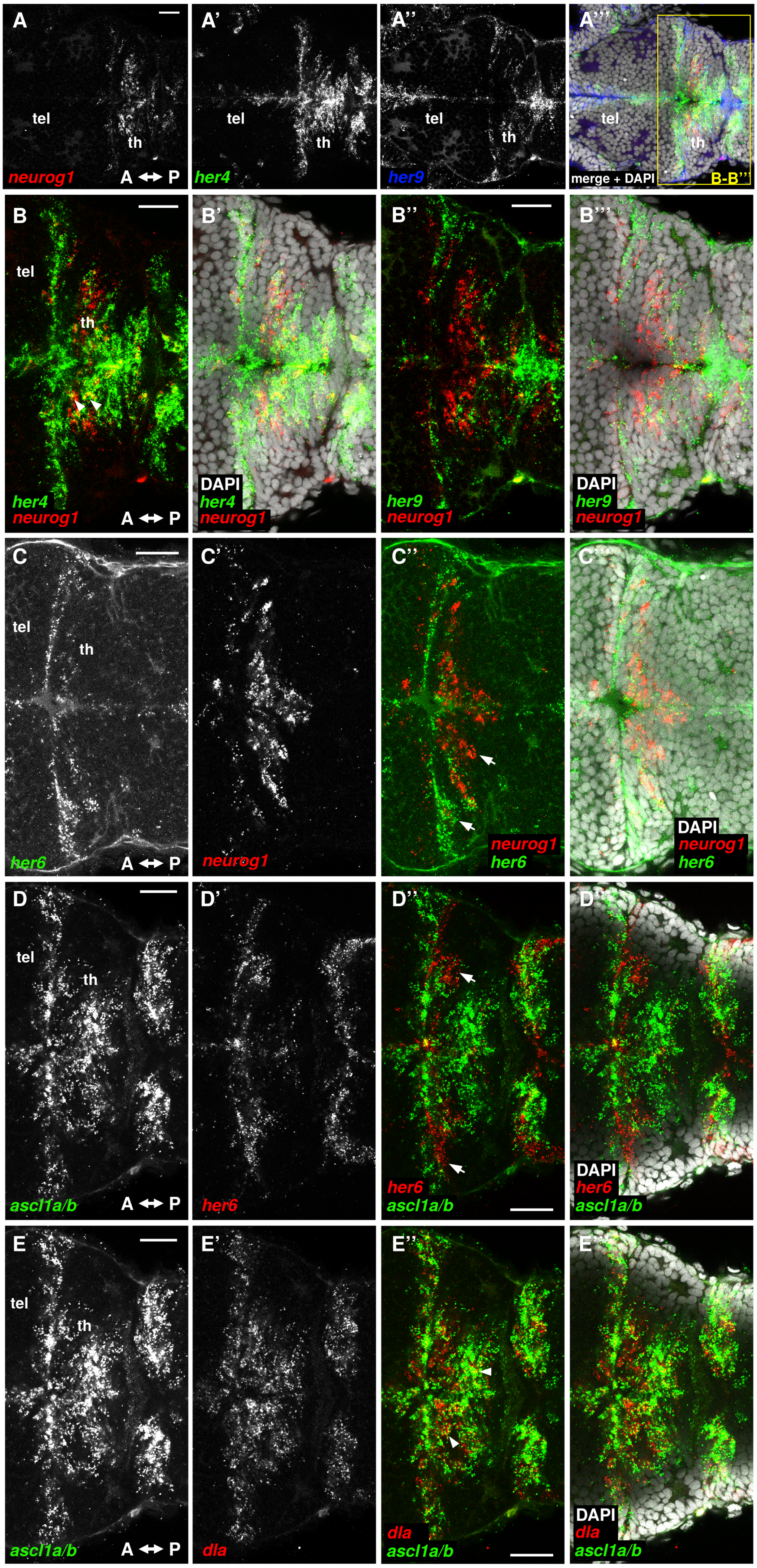

**Fig. S6 Expression analysis of *her* genes in relation to proneural genes at 3 dpf.**

**(A-E''')** Whole mount hybridization chain reaction (HCR), confocal horizontal image planes at level of thalamus proper. **(A-A''')** Overview of triple HCR for *neurog1* (A, red in A'''), *her4* (A', green in A''') and *her9* (A'', blue in A''') and merged with DAPI stain (A'''). **(B'-B''')** Magnifications of the yellow rectangle in A''' with focus on the dorsal thalamus. B and B' show overlapping expression domains of *neurog1* (red) and *her4* (green) in the thalamus (arrowheads). B'' and B''' show largely non-overlapping expression of *neurog1* and *her9*. **(C-C''')** *neurog1* (red) and *her6* (green) show largely non-overlapping expression (arrows). **(D-D''')** *ascl1a/b* (green) and *her6* (red) also show largely non-overlapping expression (arrows). **(E-E''')** *ascl1a/b* (green) and *dla* (red) show overlapping expression domains (arrowheads). DAPI nuclear stain (white) as indicated. Three embryos each were analyzed per HCR combination. Scale bars in A, B, C, D and E, 20 μm. Abbreviations: A-P anterior-posterior orientation; tel, telencephalon; th, thalamus proper.

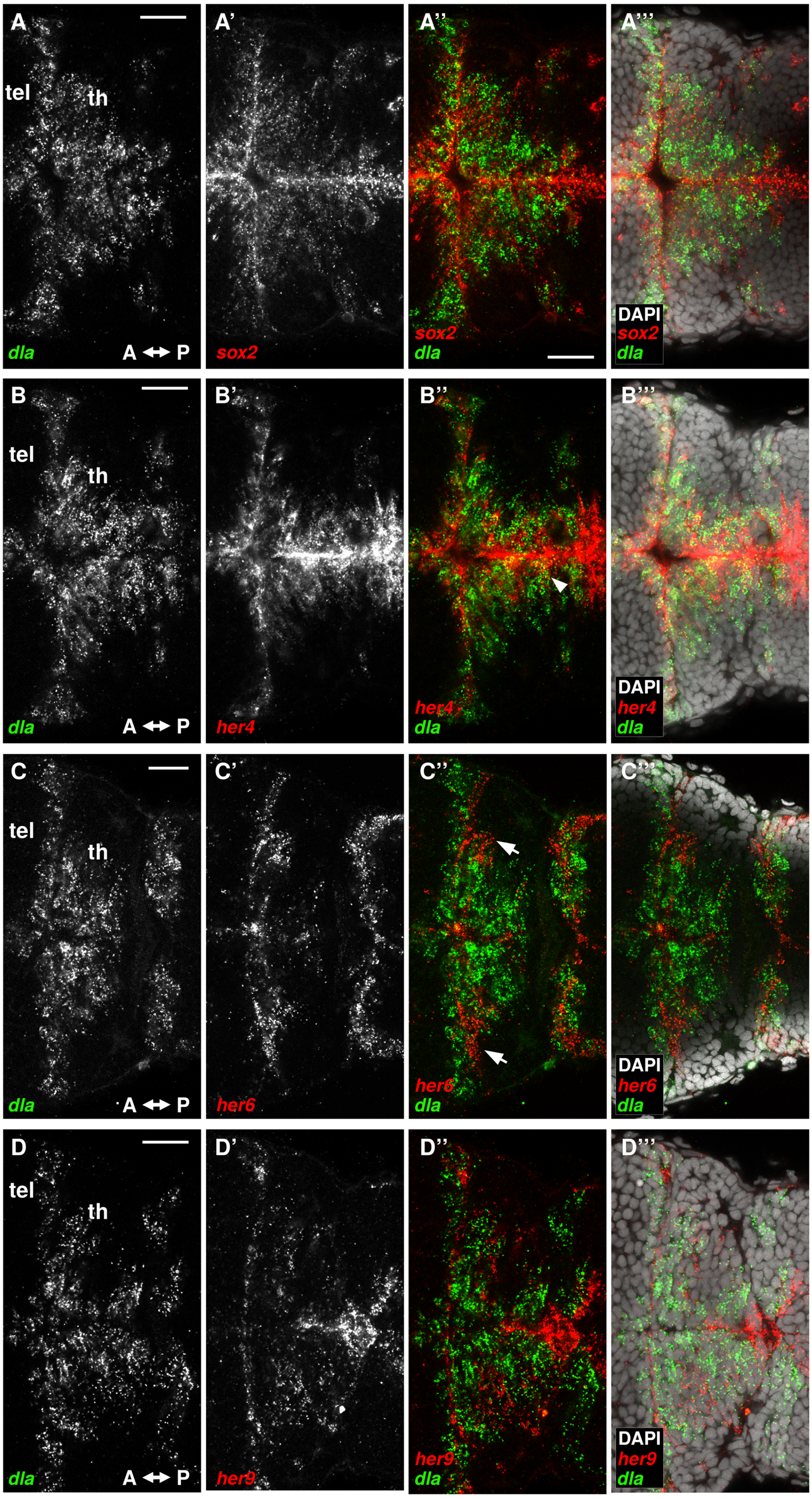

90

91 **Fig. S7 Expression analysis of *her* genes in relation to *dla*.**

**(A-D''')** Whole mount hybridization chain reaction (HCR) confocal horizontal image planes at level of thalamus proper. **(A-A''')** Overlapping expression domains of *sox2* (red) and *dla* (green). **(B-B''')** Mostly overlapping expression domains of *her4* (red) and *dla* (green) (arrowhead). **(C-C''')** *dla* (green) and *her6* (red) have largely non-overlapping expression (arrows). **(D-D''')** *dla* (green) and *her9* (red) show mostly non-overlapping expression. DAPI nuclear stain (white) as indicated. Numbers of embryos analyzed: A to C n=3 each; D n=2. Scale bars, 20 µm. Abbreviations: A-P anterior-posterior orientation; tel, telencephalon; th, thalamus proper.

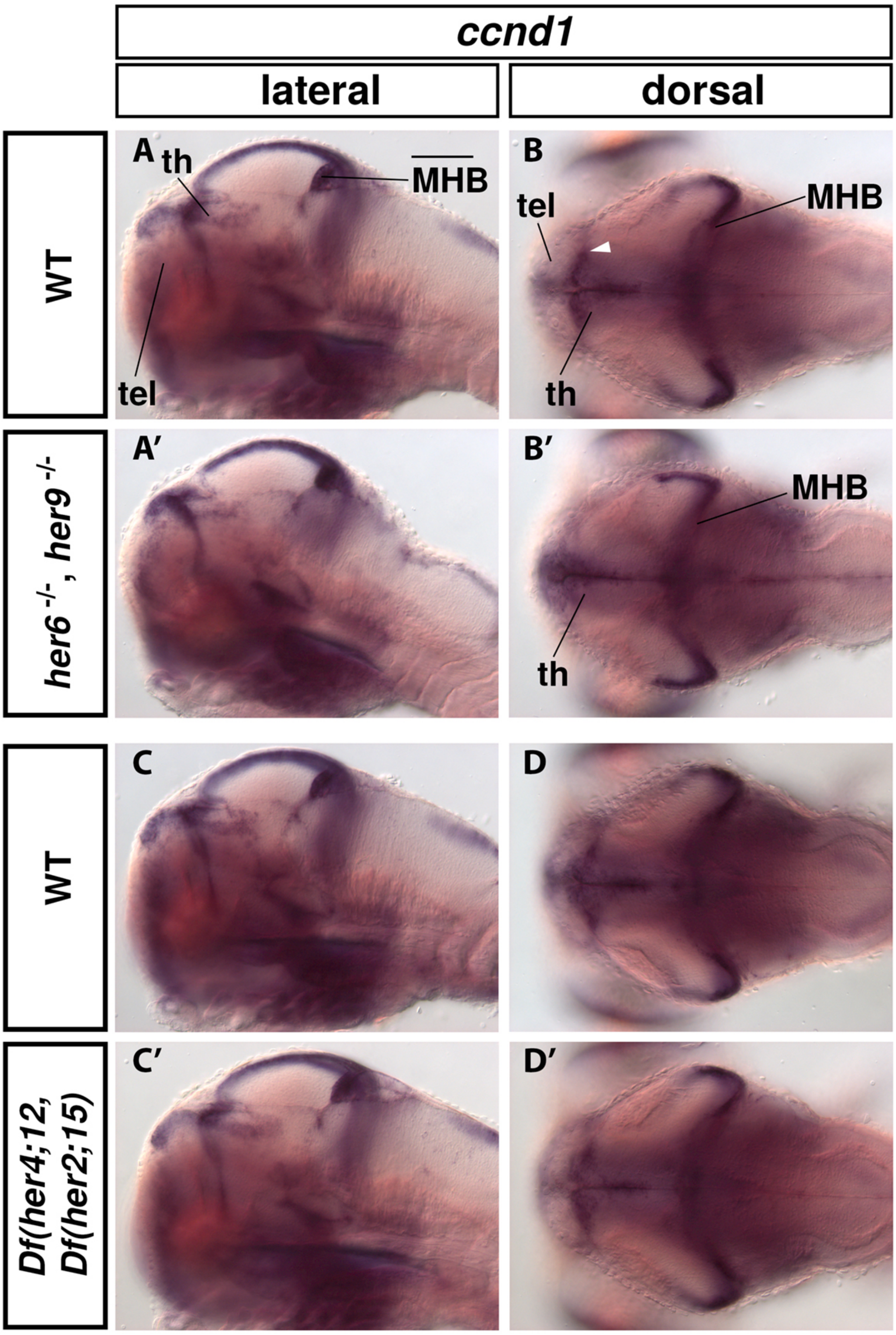

**Fig. S8 *ccnd1* expression in *her6, her9* double mutants and *Df(her4;12)*, *Df(her2;15)* double deficiency mutants.**

**(A-D')** *ccnd1* expression in 4 dpf wild type and mutant embryos visualized by WISH, genotypes as indicated. In the lateral views, single midline sagittal DIC optical sections are shown (A, A', C, C'). In the dorsal views, single horizontal DIC optical sections at the level of the thalamus are shown (B, B', D, D'). At least three larvae were imaged per condition and one representative image was selected. Anterior at left, dorsal at top. Scale bar 100 µm for all images. Abbreviations: MHB, midbrain-hindbrain boundary; tel, telencephalon; th, thalamus proper.

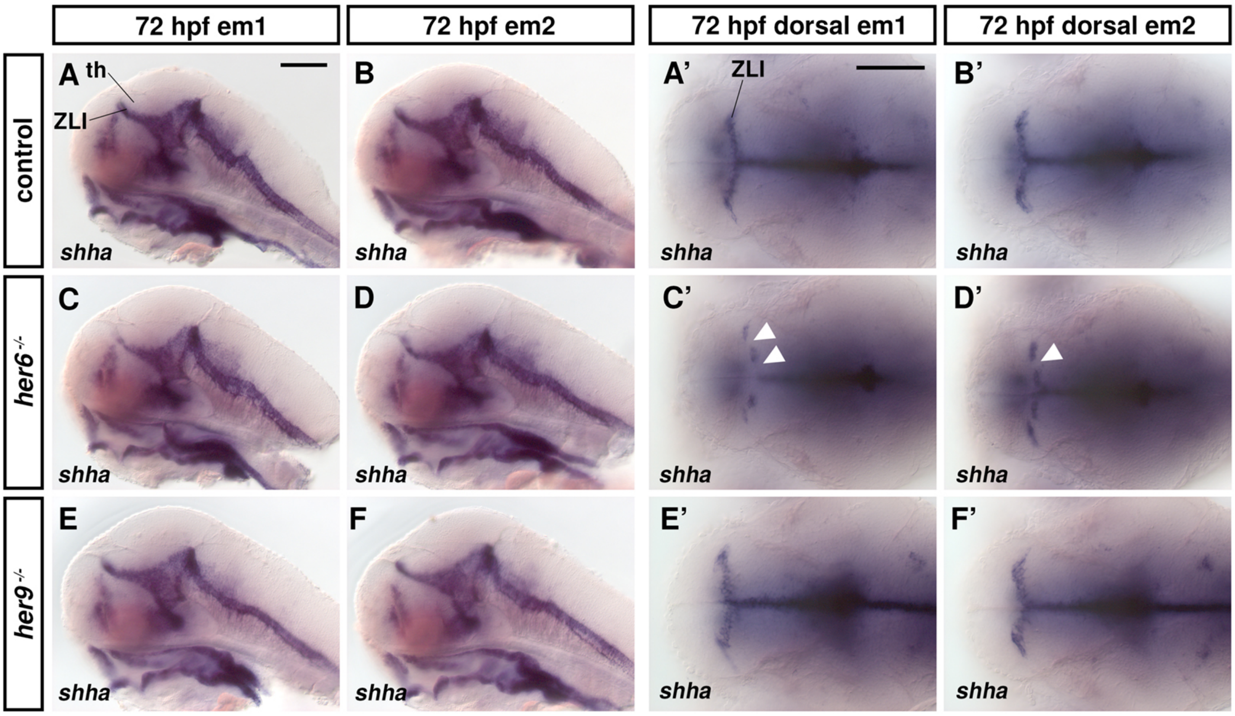

**Fig. S9 *shha* expression in *her6* and *her9* single mutants.**
**(A-F')** WISH of *shha* in control (heterozygous for *her9*) and *her6* or *her9* single mutants as indicated. **(A-F)** Lateral view midsagittal optical sections of *shha* expression in 72 hpf larvae. **(A'-F')** Dorsal view horizontal optical sections of *shha* expression in 72 hpf larvae at the level of the thalamus. Three larvae were imaged per condition and two representative images (em1 and em2) were selected since phenotypes of different severity were observed in *her6*$^{-/-}$ single mutant embryos (C' and D'). A and A' as well as B and B' show different embryos each. Anterior at left, dorsal at top. Scale bar in A is 100 µm for A-F; scale bar in A' is 100 µm for A'-F'. th, thalamus proper; ZLI, zona limitans intrathalamica.

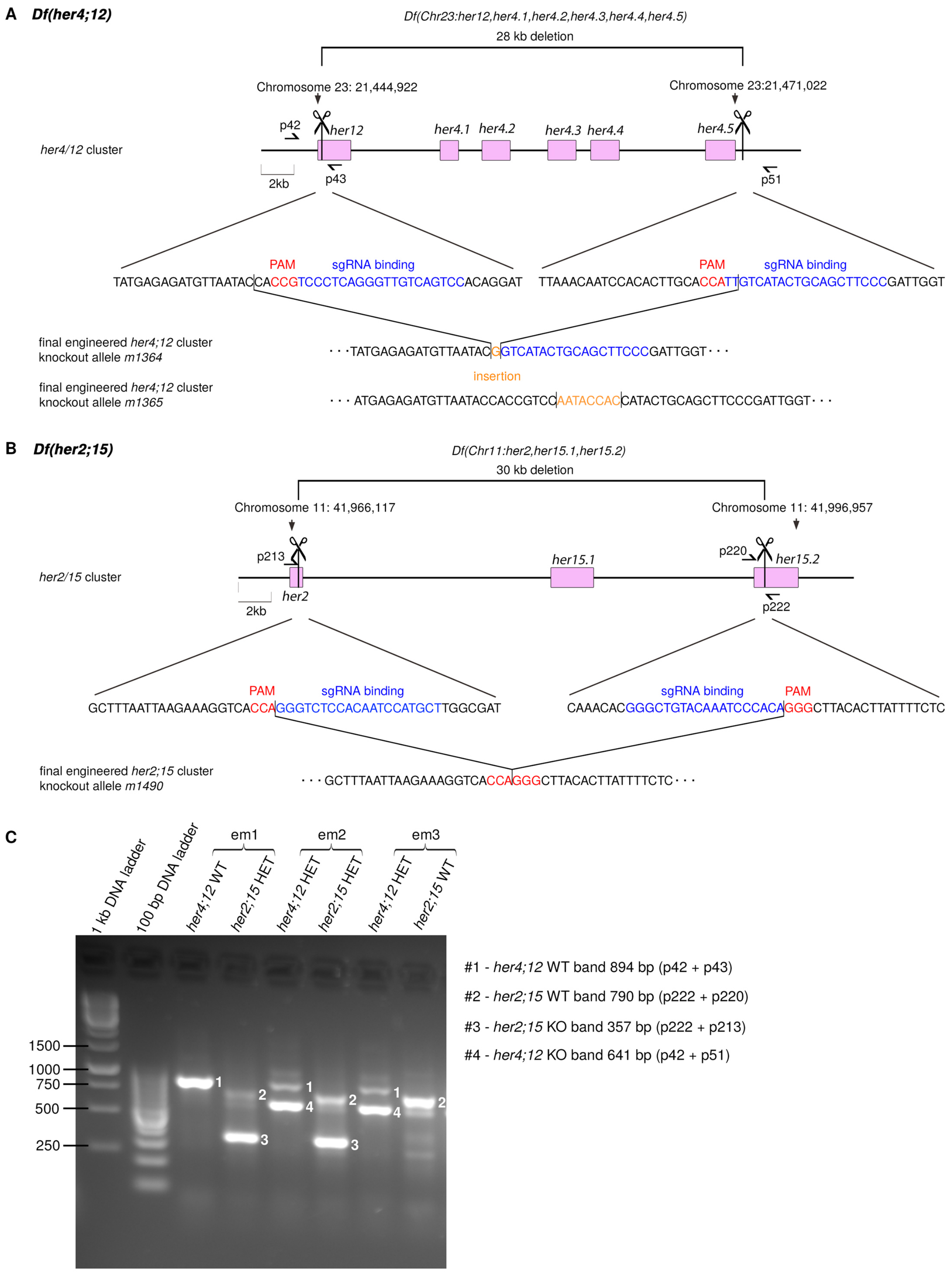

**Fig. S10 Strategy for generating *Df(her4;12)* and *Df(her2;15)*.**
**(A, B)** Knockout strategy to generate *Df(her4;12)* and *Df(her2;15)* using the CRISPR/Cas9 system. Scissors indicate where the sgRNAs bind and square brackets indicate the deletion. Half-arrows indicate binding sites for primers which were used in genotyping PCRs (tagged primer names). In the base sequence, PAM sequences are indicated in red, sgRNA binding sites are indicated in blue, and insertions are indicated in orange. Three alleles (*Df(her4;12)m1364, Df(her4;12)m1365* and

*Df(her2;15)m1490*) were used in this study, and the exact sequence mutation of the final cluster deficiencies are shown. **(C)** Example gel picture of PCR products amplified for WT and *Df(her4;12)* and *Df(her2;15)* deficiency alleles, primers as indicated in (A, B). Primer p42 and p43 amplify a *her4;12* WT genomic fragment, while p42 + p51 amplify a *Df(her4;12)m1365* allele specific genomic fragment linking the deficiency ends. Primer p222 + p220 amplify the indicated genomic sequence of the *her2;her15* WT allele, while p222 and p213 amplify the *Df(her2;15)m1490* specific genomic fragment linking the deficiency ends. p43 and p220 binding sites are deleted in *Df(her4;12)* or *Df(her2;15)* alleles, respectively. The numbers 1 to 4 shown in the gel picture next to gel bands mark the identity of the bands as indicated in the index #1 - #4 at the right. Three embryos (em1 to em3) with the indicated genotypes are shown as examples.

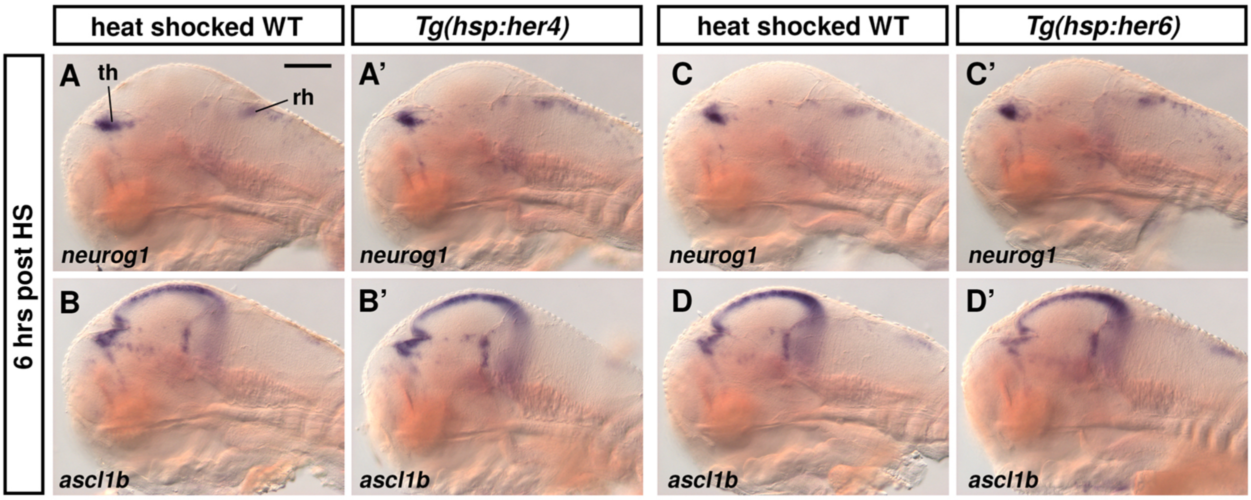

**Fig. S11 Proneural gene expression is back to normal 6 hrs after heat shock overexpression of Her4 or Her6.**
**(A-D')** WISH for expression of *neurog1* and *ascl1b* in larvae fixed 6 hrs after heat shock induced overexpression of Her4 or Her6, respectively. **(A', B')** *Tg(hsp:her4-FLAG)* was used to overexpress Her4. **(C', D')** *Tg(hsp:her6-FLAG)* was used to overexpress Her6. Heat shocked wild type siblings serve as control (A, B, C, D). All larvae shown in this panel were heat shocked at 70 hpf for 30 min and were fixed at 76 hpf (6 hrs after heat shock start). Three larvae per condition were imaged and the WISH staining patterns were consistent. th, thalamus proper; rh, rhombencephalon. Lateral views midsagittal optical sections; scale bar 100 µm in A for all panels.

### Supplementary Movies Legends

**Movie M1. Analysis of NPC marker expression reveals *sox2*[low] expressing cells as progenitors.** Movie from stack of HCR expression analysis shown in Fig. 4C-C'' and Fig. S5 (see legends for details). The movie analyzes *ascl1b* (red), *neurog1* (blue) and *sox2* (green) expression in the thalamus of a 3 dpf WT larva. The movie starts with the dorsal most horizontal plane of the image stack and ends with the ventral most plane. Orientation: anterior to the left. pin, pineal gland; pth, prethalamus; th, thalamus proper.

**Movie M2. Combined movie of WT (left) and *her6, her9* double mutant larvae (right).** Combined movie of larvae shown in Fig. 4A (WT; left half) and Fig. 4B (*her6*$^{-/-}$, *her9*$^{-/-}$ double mutant; right half). The movie shows an anti-Sox2 (magenta) and anti-pH3 (green) double immunostaining horizontal image plane confocal stack of 3 dpf WT larvae recorded from a dorsal view. The movie starts with the dorsal-most plane of the image stack and ends with the ventral-most plane. Black slides were inserted when one stack had less z-planes than the other stack (here the ventral slides of the WT part). Since mutant and WT have different phenotypic appearances and also differ in size, the planes were aligned according to the thalamic Sox2 expression. This means that more ventral regions might be shifted. Orientation: anterior to the left.

**Movie M3. Combined movie of WT (left) and *Df(her4;12), Df(her2;15)* double mutant (right).** Combined movie of larvae shown in Fig. 6A (WT; left half) and Fig. 6B [*Df(her4;12), Df(her2;15)* double mutant; right half]. The movie shows an anti-Sox2 (magenta) and anti-pH3 (green) double immunostaining confocal horizontal image stack of 3 dpf WT larvae recorded from a dorsal view. The movie starts with the dorsal-most plane of the image stack and ends with the ventral-most plane. Black slides were inserted when one stack had less z-planes than the other stack. Orientation: anterior to the left.

**Movie M4. Combined movie of WT (left) and *Df(her4;12), Df(her2;15), her6*$^{-/-}$, *her9*$^{-/-}$ mutant (*her*UDM, right).** Combined movie of larvae shown in Fig. 9C (WT; left half) and Fig. 9D [*Df(her4;12), Df(her2;15), her6*$^{-/-}$, *her9*$^{-/-}$ combined mutant; right half]. The movie shows an anti-Sox2 (magenta) and anti-pH3 (green) immunostaining confocal horizontal image stack of 3 dpf WT larvae recorded from a dorsal view. The movie starts with the dorsal most plane of the image stack and ends with the ventral most plane. Since WT and *her*UDM have different phenotypic appearances and also differ in size, black slides were inserted at the beginning and end to align zslides according to the thalamic Sox2 expression. This means that more ventral and dorsal regions might be shifted. Orientation: anterior to the left.

**Movie M5. Combined movie of *her6* rescue (left) and *her2, her15* rescue (right) of *her*UDM mutants.**

Combined movie of larvae shown in Fig. 9E ("*her6* rescue" of *her*UDM mutant; left half; genotype *Df(her4;12), Df(her2;15), her6*$^{+/-}$, *her9*$^{-/-}$ mutant) and Fig. 9F ("*her2, her15* rescue" of *her*UDM mutant; right half; genotype *Df(her4;12), Df(her2;15) heterozygous, her6*$^{-/-}$, *her9*$^{-/-}$ mutant). The movie shows an anti-Sox2 (magenta) and anti-pH3 (green) immunostaining confocal horizontal image stack of 3 dpf WT larvae recorded from a dorsal view. The movie starts with the dorsal most plane of the image stack and ends with the ventral most plane. Since both embryos have different phenotypic appearances and also differ in size, the planes were aligned according to the thalamic Sox2 expression. Orientation: anterior to the left.

**Supplementary Table 1:** **Primer and sgRNA templates (page 02-04)**

**Supplementary Table 2:** **HCR Oligonucleotides (page 05-08)**

**Supplementary Table 3:** **PCR conditions (page 09-10)**

**Supplementary Table 4:** **Numbers of embryos documented or analyzed in each experiment (page 11-13)**

**Supplementary Table 5:** ***her4* and *her6* Q-PCR data and p-values for comparisons (page 14-16)**

**Supplementary Table 1: Primer and sgRNA templates**

Primers are given with a short description, a short ID and the primer sequence. Oligonucleotide Sequences for the transcription of specific sgRNAs include: The T7 promoter (left) shown in lower case letters, the target specific region of the sgRNA (middle) in capital letters, and the annealing site for the constant oligonucleotide (right, lower case).

| Description of primer | Short ID | sequence |
|---|---|---|
| *Df(her2;15)* | | |
| sgRNA for *Df(her2;15)* at *her2* end | p211 - Oligo38 | 5′- taatacgactcactataGGCATGGATTGTGGAGACCCgttttagagctagaaatagcaag -3′ |
| sgRNA for *Df(her2;15)* at *her15.2* end | p219 - Oligo43 | 5′- taatacgactcactataGGGCTGTACAAATCCCACAgttttagagctagaaatagcaag -3′ |
| *Df(her2;15)* control primer F at *her2* end | p213F | 5′- AACACCTCTGCAGGCTACAC -3′ |
| *Df(her2;15)* control primer R at *her15.2* end | p222R | 5′- CTTGCATGCAGTTCAATCTCACA -3′ |
| *Df(her2;15)* control primer F at *her15.2* end | p220F | 5′- GGATCCTGCTGCTGGAACTC -3′ |

| | | |
|---|---|---|
| *Df(her4;12)* | | |
| sgRNA for *Df(her4;12)* at *her2* end | p39 - Oligo12 | 5′- taatacgactcactataGGACTGACAACCCTGAGGGAgttttagagctagaaatagcaag -3′ |
| sgRNA for *Df(her4;12)* at *her4.5* end | p47 - Oligo16 | 5′- taatacgactcactataGGGAAGCTGCAGTATGACAAgttttagagctagaaatagcaag -3′ |
| *Df(her4;12)* control primer F at *her12* end | p42F | 5′- TATTGTACTTCTTGCACATTTTGCG -3′ |
| *Df(her4;12)* control primer R at *her12* end | p43R | 5′- GAAAAGATGCGTCGGGATCG -3′ |
| *Df(her4;12)* control primer R at *her15.2* end | p51R | 5′- CTCATGGCGAAGGATTGTCG -3′ |

| | | |
|---|---|---|
| *her9* knockout | | |
| sgRNA for *her9* knockout | p186 - Oligo26 | 5′-taatacgactcactataGGACTTCTTAACCACCTGTCgttttagagctagaa-3′ |
| sgRNA for *her9* knockout | p187 - Oligo27 | 5′-taatacgactcactataGGGTATGAGATCCACTGGCAgttttagagctagaa-3′ |
| *her9* m1368 knockout control PCR F | p197F | 5′-CCGGACTCAACTTTGGTGTTTG-3′ |
| *her9* m1368 knockout control PCR R | p198R | 5′-GATGGGTAACGTTGAAGGAAGC-3′ |

| Description of primer | Short ID | sequence |
|---|---|---|
| *her6* knockout | | |
| sgRNA for *her6* knockout | p3-Oligo04 | 5′-taatacgactcactataGCGAGAATCAACGAAAGCTTgttttagagctagaaatagcaag-3′ |
| sgRNA for *her6* knockout | p4-Oligo05 | 5′-taatacgactcactataCGGTACTTCCCAAGAACGGTgttttagagctagaaatagcaag-3′ |
| *her6* m1358 knockout control PCR F | p5F | 5′-TCAGCGTACTTGACAGCGTT-3′ |
| *her6* m1358 knockout control PCR R | p10R | 5′-TCACATGTGGACAGGAACCG-3′ |

| | | |
|---|---|---|
| qPCR primer | | |
| pPCR *her6* F | p372F | 5′- TCCAAACGGCCCTGTTATTCC-3′ |
| pPCR *her6* R | p373R | 5′- AACGGAGTCTGACGTGACGG-3′ |
| pPCR *her9* F | p376F | 5′- CAACCCAGCGTTTGCTTCTG-3′ |
| pPCR *her9* R | p377R | 5′- CTGACACCAACGGGACTGAC-3′ |

| Description of primer | Short ID | sequence |
|---|---|---|
| For cloning of in situ hybridization probes | | |
| ISHprobe recognising *her4.1 ,4.2, 4.3, 4.4, 4.5* F | 159F | 5′- CTGAGAAAGCCCATGGTGGA-3′ |
| ISHprobe recognising *her4.1 ,4.2, 4.3, 4.4, 4.5* R | 160R | 5′- CTACCAGGGTCTCCAGATGTGA-3′ |
| ISHprobe *her6* F | p147F | 5′- GCTGCCCTAAACACAGATCCC-3′ |
| ISHprobe *her6* R | p148R | 5′- TCATTTCAAATGTCATTTATTTGTCTTCCAAAAAG-3′ |
| ISHprobe *her9* F | p149F | 5′- CAGCCTTGTCAGCTGACACAA-3′ |
| ISHprobe *her9* R | p150R | 5′- CGTTATTTTGATTTATTCAGCATCAAACATAAACAC-3′ |
| ISHprobe *her15.1, her15.2* F | p157F | 5′- CAAGTCCATGCTGGAGAAAGAGT-3′ |
| ISHprobe *her15.1, her15.2* R | p158R | 5′- CATCTTTCATATAAAGAAACACCAATAAAGCA-3′ |
| ISHprobe *her2* F | p154F | 5′- CTGAGGAAACCGGTGGTGG-3′ |
| ISHprobe *her2* R | p155R | 5′- TGGTTTCAACTCAACATGAGATACTTAAAC-3′ |
| ISHprobe *her12* F | p152F | 5′- CTGAGGAAGCCGATAGTTGAAAAGA-3′ |
| ISHprobe *her12* R | p153R | 5′- CTAACAACAAACAAACAATAACACATCATTGG-3′ |
| ISHprobe *her8a* F | p322F | 5′- CACTGCTTGGAAGCAAATGA-3′ |
| ISHprobe *her8a* R | p327R | 5′- agaaaaatCAGCTTCAATATGAGGTACTAGTC-3′ |
| ISHprobe *her8.2* F | p260F | 5′- GAGCCCGACAGAGATACAGC-3′ |
| ISHprobe *her8.2* R | p259R | 5′- CAACACAGTATTAGTAGCCAAAGGG-3′ |

| primer for identification of *Tg(UAS:NICD)kca3Tg* zebrafish | | |
|---|---|---|
| NICD F | | 5′-CATCGCGTCTCAGCCTCAC-3′ |
| NICD R | | 5′-CGGAATCGTTTATTGGTGTCG-3′ |

**Supplementary Table 2: HCR Oligonucleotides**

Oligonucleotide sequences for *olig2, her4, her9*, and *her15* HCR. Upper-case letters indicate gene specific sequence. Spacer 1 and Spacer 2 (aa or ta) are given in lower-case letters adjacent to the gene specific sequences. Initiator 1 and Initiator 2 are given in lower-case letters next to the spacer sequences.

| Description | Sequence |
|---|---|
| **Oligonucleotides for *olig2* HCR probe** | |
| HCR_Dan_olig2_B1_S1-1 | gaggagggcagcaaacggaaCACTCGTGCAGTGTTTGTGTTTGGC |
| HCR_Dan_olig2_B1_S1-2 | GAATTGACTTGTAAGGCTATTCCAGtagaagagtcttcctttacg |
| HCR_Dan_olig2_B1_S2-1 | gaggagggcagcaaacggaaTCCATGGCGTTCAGTGCGCTCTCAG |
| HCR_Dan_olig2_B1_S2-2 | CTGCTGGACACTCGGCTCGTGTCAGtagaagagtcttcctttacg |
| HCR_Dan_olig2_B1_S3-1 | gaggagggcagcaaacggaaCCTCTTAACTCCGGTGGAGAATCGC |
| HCR_Dan_olig2_B1_S3-2 | GCGCTGGAGATGCCCCGCAGATCTGtagaagagtcttcctttacg |
| HCR_Dan_olig2_B1_S4-1 | gaggagggcagcaaacggaaCTCATTCTCGGAGAGAAGTTTACGG |
| HCR_Dan_olig2_B1_S4-2 | GTTGATCTTTAGGCGCATGCTCTGCtagaagagtcttcctttacg |
| HCR_Dan_olig2_B1_S5-1 | gaggagggcagcaaacggaaAGTTGCGCGCGAGCAGCAGAGTGGC |
| HCR_Dan_olig2_B1_S5-2 | CCAGCGAGTTGCTCAGCATAAGGATtagaagagtcttcctttacg |
| HCR_Dan_olig2_B1_S6-1 | gaggagggcagcaaacggaaGACGGCTGAAAGCCCCGTCCCGGAA |
| HCR_Dan_olig2_B1_S6-2 | GGCTTTGAGGAGTCCGTGATGCGGCtagaagagtcttcctttacg |
| HCR_Dan_olig2_B1_S7-1 | gaggagggcagcaaacggaaACGGCGCTCATACCGGGAACGTGAG |
| HCR_Dan_olig2_B1_S7-2 | GAGTCACTGGTCAGCCGTGGCATGCtagaagagtcttcctttacg |
| HCR_Dan_olig2_B1_S8-1 | gaggagggcagcaaacggaaCTAAGGAAGGTTTGCCATTTCCAAC |
| HCR_Dan_olig2_B1_S8-2 | TTGTTCTTTCAGTCCTATAGTCGAGtagaagagtcttcctttacg |
| HCR_Dan_olig2_B1_S9-1 | gaggagggcagcaaacggaaAAGAGCGCTTGGCAAATGAATTCTT |
| HCR_Dan_olig2_B1_S9-2 | GCAGGAATGAACAGGTGCTTCTTTAtagaagagtcttcctttacg |
| HCR_Dan_olig2_B1_S10-1 | gaggagggcagcaaacggaaCGGATAAGCTTGTGCAGAAAGAAGA |
| HCR_Dan_olig2_B1_S10-2 | GTTGATCATTCTTCGCATCGGTAATtagaagagtcttcctttacg |

| Description | Sequence |
|---|---|
| **Oligonucleotides for *her4* HCR probe** | |
| HCR_Dan_her4_B1_S1-1 | gaggagggcagcaaacggaaGAAATCAAGCGTCATCTCCAGGATA |
| HCR_Dan_her4_B1_S1-2 | CGCACTGCTTTTCTGAGAGCGTCTCtagaagagtcttcctttacg |
| HCR_Dan_her4_B1_S2-1 | gaggagggcagcaaacggaaTGCACACATCTGGAGCGTCCGTCAC |
| HCR_Dan_her4_B1_S2-2 | CACTGAGACAGAAAGCTGACGGCCTtagaagagtcttcctttacg |
| HCR_Dan_her4_B1_S3-1 | gaggagggcagcaaacggaaTCTTGTGTGGCTCTGCGTCTGCACT |
| HCR_Dan_her4_B1_S3-2 | CTGCATGTGCAGGAAGAGCTTCATCtagaagagtcttcctttacg |
| HCR_Dan_her4_B1_S4-1 | gaggagggcagcaaacggaaTCCACACGTGTGTGCTGGTCTGCAG |
| HCR_Dan_her4_B1_S4-2 | GCGTGTGTTTCAGTGGTCTGAGGATtagaagagtcttcctttacg |
| HCR_Dan_her4_B1_S5-1 | gaggagggcagcaaacggaaGGGCTGGAGTGTGTTGTTTGGCGCT |
| HCR_Dan_her4_B1_S5-2 | TCTACCAGGGTCTCCAGATGTGACTtagaagagtcttcctttacg |
| HCR_Dan_her4_B1_S6-1 | gaggagggcagcaaacggaaCCTCAATGGTACGGCGGGTGCTCTG |
| HCR_Dan_her4_B1_S6-2 | CTCAGACTGGCATCGTGACTCGTGTtagaagagtcttcctttacg |
| HCR_Dan_her4_B1_S7-1 | gaggagggcagcaaacggaaTACTTTCTGAACTTCAGTCCATGCC |
| HCR_Dan_her4_B1_S7-2 | GAAGCTGCAGAGGCGGAAATCAATTtagaagagtcttcctttacg |
| HCR_Dan_her4_B1_S8-1 | gaggagggcagcaaacggaaCTCTGAAAGCAGATTAAATGTCCTG |
| HCR_Dan_her4_B1_S8-2 | CTCACACAGTTTCACTAAACTCTCTtagaagagtcttcctttacg |
| HCR_Dan_her4_B1_S9-1 | gaggagggcagcaaacggaaATGATTGTGTGCATTGAGATCAAAC |
| HCR_Dan_her4_B1_S9-2 | CACTGTAATAATCATTCTACACAGCtagaagagtcttcctttacg |
| HCR_Dan_her4_B1_S10-1 | gaggagggcagcaaacggaaCAAACCCAATATGGGTGAAATGTTC |
| HCR_Dan_her4_B1_S10-2 | GTGGTCATCGTATAGATGAAGAGAAtagaagagtcttcctttacg |

| Description | Sequence |
|---|---|
| **Oligonucleotides for *her9* HCR probe** | |
| HCR_Dan_her9_B4_S1-1 | cctcaacctacctccaacaaATCTCCCACTGCCACACGCCAGCGG |
| HCR_Dan_her9_B4_S1-2 | GGCGCAGCGCTTCAGAGGGCAAGTCattctcaccatattcgcttc |
| HCR_Dan_her9_B4_S2-1 | cctcaacctacctccaacaaATCAGCTTTATGACTTATAGTCGGG |
| HCR_Dan_her9_B4_S2-2 | TCACGAGAGAGAGTGTGAGTCCCTCattctcaccatattcgcttc |
| HCR_Dan_her9_B4_S3-1 | cctcaacctacctccaacaaCCGGTAGAAGCCAAAAAGTGAATAT |
| HCR_Dan_her9_B4_S3-2 | TGGCATGATCAAACACCAAAGTTGAattctcaccatattcgcttc |
| HCR_Dan_her9_B4_S4-1 | cctcaacctacctccaacaaCCTGTTGAGCGGGGGCAGGCTGAGG |
| HCR_Dan_her9_B4_S4-2 | GCACGTGAAGAGGCTGAGCCAAATGattctcaccatattcgcttc |
| HCR_Dan_her9_B4_S5-1 | cctcaacctacctccaacaaACTGGCGTTGACAGTTACTGGAACG |
| HCR_Dan_her9_B4_S5-2 | CGTGGGAGCCGAGCTGGCCTGGACGattctcaccatattcgcttc |
| HCR_Dan_her9_B4_S6-1 | cctcaacctacctccaacaaGAGGTCATTCCCTGGACAGGAGATG |
| HCR_Dan_her9_B4_S6-2 | CTGACGGCTTGTGGAACGCCCGAGAattctcaccatattcgcttc |
| HCR_Dan_her9_B4_S7-1 | cctcaacctacctccaacaaATTGCTTTCTGCTCCGGCACTGACA |
| HCR_Dan_her9_B4_S7-2 | AGTCTACCAGGGTCTCCACACCGGCattctcaccatattcgcttc |
| HCR_Dan_her9_B4_S8-1 | cctcaacctacctccaacaaCGGAACTGAACGTATATTCTTCACC |
| HCR_Dan_her9_B4_S8-2 | ACAAAAGGTCAACAAAGTCCCTTAGattctcaccatattcgcttc |
| HCR_Dan_her9_B4_S9-1 | cctcaacctacctccaacaaACAACATCCAAATATTGGAATCAGC |
| HCR_Dan_her9_B4_S9-2 | GTTGTATTCACAACAGCACATTCATattctcaccatattcgcttc |
| HCR_Dan_her9_B4_S10-1 | cctcaacctacctccaacaaGCTTTGGCATGATACAGTCTTTTAC |
| HCR_Dan_her9_B4_S10-2 | AACACTCATCATGAAAAGAACAAGGattctcaccatattcgcttc |

| Description | Sequence |
|---|---|
| **Oligonucleotides for *her15* HCR probe** | |
| HCR_Dan_her15_B2_S1-1 | cctcgtaaatcctcatcaaaCTCTGAGGCCGCTGGATGAGGCGTC |
| HCR_Dan_her15_B2_S1-2 | GTCCCAGTCAGGATTAGCTTTCCCAaaatcatccagtaaaccgcc |
| HCR_Dan_her15_B2_S2-1 | cctcgtaaatcctcatcaaaAGGAGCTCATTAGCATTACAAAGCC |
| HCR_Dan_her15_B2_S2-2 | GAATGAGGGTCTCGCTGCTCTATTTaaatcatccagtaaaccgcc |
| HCR_Dan_her15_B2_S3-1 | cctcgtaaatcctcatcaaaCTCTGAGCAGAGCGAGGGGTTTGTG |
| HCR_Dan_her15_B2_S3-2 | ATTGCTTCTTCAGGAGAGATGCTGTaaatcatccagtaaaccgcc |
| HCR_Dan_her15_B2_S4-1 | cctcgtaaatcctcatcaaaAAGCTTGGAGTATTCAGTCATATAC |
| HCR_Dan_her15_B2_S4-2 | TCGCAATTTGTGCTTCTCCTTGTTGaaatcatccagtaaaccgcc |
| HCR_Dan_her15_B2_S5-1 | cctcgtaaatcctcatcaaaTGTGGAGTCTTGGGCCGCAGCTGCT |
| HCR_Dan_her15_B2_S5-2 | TGCGAGTAGCCCTCGATCTGAGCGTaaatcatccagtaaaccgcc |
| HCR_Dan_her15_B2_S6-1 | cctcgtaaatcctcatcaaaCAGCCTCGGAGCCCACAGACAGAAA |
| HCR_Dan_her15_B2_S6-2 | GGGCTTCTTGCTGGAGACGTTGAGCaaatcatccagtaaaccgcc |
| HCR_Dan_her15_B2_S7-1 | cctcgtaaatcctcatcaaaGTGTGCTGCTGATGGGGAGCTTCAG |
| HCR_Dan_her15_B2_S7-2 | TGCGCCCGCGGCTCCTGCTTGATGTaaatcatccagtaaaccgcc |
| HCR_Dan_her15_B2_S8-1 | cctcgtaaatcctcatcaaaCATCTAGTGTTCGAGCGTCTCTACC |
| HCR_Dan_her15_B2_S8-2 | TCTCCTACATAAGAGACGGTAAACCaaatcatccagtaaaccgcc |
| HCR_Dan_her15_B2_S9-1 | cctcgtaaatcctcatcaaaATCAGTGATGTACTATAGTAAATGG |
| HCR_Dan_her15_B2_S9-2 | AATCTCTCATTTCTTCACACGATGTaaatcatccagtaaaccgcc |
| HCR_Dan_her15_B2_S10-1 | cctcgtaaatcctcatcaaaCTAAACGTCAGCAGAAGACGAGGCG |
| HCR_Dan_her15_B2_S10-2 | ACACTAATAAACCTCTGATCATAACaaatcatccagtaaaccgcc |

**Supplementary Table 3: PCR conditions**

PCR programs for genotyping *Df(her4;12)*, *Df(her2;12), her6* and *her9* mutant zebrafish and zebrafish larvae. *her6* and *her9* qPCR programs and NICD genotyping PCR program. The primer sequences are given in Supplementary Table 1.

| *Df(her4;12)* | | | |
|---|---|---|---|
| step | Temp in °C | duration | primer<br>p42 |
| 1 | 95 | 3min | p43 |
| 2 | 95 | 30s | p51 |
| 3 | 59 | 40s | |
| 4 | 72 | 50s | |
| 5 | 35x to 2 | | |
| 6 | 72 | 5min | |
| 7 | 4 | end | |

| *her6* qPCR | | | |
|---|---|---|---|
| step | Temp in °C | duration | primer<br>p372 |
| 1 | 95 | 30s | p373 |
| 2 | 95 | 15s | |
| 3 | 60 | 1 min | |
| 4 | 40x to 2 | | |
| 5 | 95 | 1min | |
| 6 | 40 | 1min | |
| 7 | from 65 °C to 95 °C with 0.07 °C per second | | |

| *Df(her2;15)* | | | |
|---|---|---|---|
| step | Temp in °C | duration | primer<br>p213 1/4 of normal amount |
| 1 | 95 | 3min | p220 |
| 2 | 95 | 30s | p222 |
| 3 | 60 | 40s | |
| 4 | 72 | 60s | |
| 5 | 35x to 2 | | |
| 6 | 72 | 5min | |
| 7 | 4 | end | |

| *her9* qPCR | | | |
|---|---|---|---|
| step | Temp in °C | duration | primer<br><br>p376 |
| 1 | 95 | 30s | p377 |
| 2 | 95 | 15s | |
| 3 | 60 | 1 min | |
| 4 | 40x to 2 | | |
| 5 | 95 | 1min | |
| 6 | 40 | 1min | |
| 7 | from 65 °C to 95 °C with 0.07 °C per second | | |

*her6* knockout

| step | Temp in °C | | duration | primer |
|---|---|---|---|---|
| | | | | p5 |
| 1 | | 95 | 3min | p10 |
| 2 | | 95 | 30s | |
| 3 | | 56 | 40s | |
| 4 | | 72 | 50s | |
| 5 | 35x to 2 | | | |
| 6 | | 72 | 5min | |
| 7 | | 4 | end | |

*her9* knockout

| step | Temp in °C | | duration | primer |
|---|---|---|---|---|
| | | | | p197 |
| 1 | | 95 | 3min | p198 |
| 2 | | 95 | 30s | |
| 3 | | 59 | 40s | |
| 4 | | 72 | 50s | |
| 5 | 35x to 2 | | | |
| 6 | | 72 | 5min | |
| 7 | | 4 | end | |

NICD genotyping

| Step | Temp in °C | | duration | primer |
|---|---|---|---|---|
| 1 | | 95 | 3min | NICD F |
| 2 | | 95 | 1min | NICD R |
| 3 | | 80 | 1min | |
| 4 | | 95 | 3min | |
| 5 | | 95 | 1min | |
| 6 | | 62 | 1min | |
| 7 | | 72 | 75s | |
| 8 | 3x to 2 | | | |
| 9 | | 95 | 1min | |
| 10 | | 62 | 45s | |
| 11 | | 72 | 45s | |
| 12 | 35x to 9 | | | |
| 13 | | 72 | 5min | |
| 14 | | 8 | end | |

**Supplementary Table 4: Numbers of embryos documented or analyzed in each experiment**

For each figure panel, the numbers of embryos documented or analyzed in each experiment are provided.

| Fig.1 |
| --- |
| three larvae were imaged per condition and one representative image was selected |

| Fig. 2 | |
| --- | --- |
| | n numbers |
| A, A' | 3 |
| B, B' | 4 |
| C, C' | 2 |
| D, D' | 2 |

| Fig. 3 | |
| --- | --- |
| | n numbers |
| C, C' | 3 |
| D, D' | 3 |
| E, E' | 2 |
| F, F' | 4 |
| G, G' | 2 |
| H, H' | 3 |
| I, I' | 3 |
| J, J' | 2 |
| K, K' | 2 |
| L, L' | 4 out of 5 |
| M, M' | 2 |
| N, N' | 4 out of 6 |

| Fig. 4 | | |
|---|---|---|
| | n numbers | |
| A-A" | 5 | (also in Supplementary Movie M1) |
| B-B" | 4 | (also in Supplementary Movie M1) |
| C-C" | 3 | |
| D-D" | 3 | |

| Fig. 5 | |
|---|---|
| | n numbers |
| A, A' | 3 |
| E, E' | 3 |
| B, B' | 3 |
| F, F' | 3 |
| C, C' | 3 |
| G, G' | 3 |
| D, D' | 3 |
| H, H' | 3 |

| Fig. 6 | |
|---|---|
| | n numbers |
| C | 5 |
| C' | 4 |
| D | 5 |
| D' | 2 |
| E, F | 2 |
| E', F' | 3 |

| Fig. 7 | | |
|---|---|---|
| | n numbers | |
| A-A'' | 4 | (also in Supplementary Movie M2) |
| B-B'' | 5 | (also in Supplementary Movie M2) |
| C-C'' | 3 | |
| D-D'' | 3 | |

| Fig. 8 |
|---|
| Three larvae were imaged per condition and one representative image was selected. |

| Fig. 9 | |
|---|---|
| Three larvae were imaged per condition and one representative image was selected, except for: | |
| B' | 9 out of 14 larvae showed the displayed pehnotype |
| E' | 9 out of 15 larvae showed the displayed pehnotype |
| J' | 11 out of 15 larvae showed the displayed pehnotype |

| Fig. 10 | | |
|---|---|---|
| | n numbers | |
| A, B | 1 | |
| A', B' | 3 | |
| C-C'' | 4 | (also in Supplementary Movie M2) |
| D-D'' | 4 | (also in Supplementary Movie M2) |
| E-E'' | 3* | *in Fig. 10E and Supplementary Movie M4, a *her2, her15* heterozygous larva was used (1 out of 3 larvae). Two more larvae were WT for *her2, her15*. |
| F-F'' | 4* | *in Fig. 10F and Supplementary Movie M4, a *her6* heterozygous larva was used (1 out of 4 larvae). 3 more larvae were *her6* WT |

### **Supplementary Table 5: Data Analysis Table of *her6* and *her9* qPCR** (Legend see page 16)

**Table 5A: qPCR Primary Data**

| A | B | C | D | E | F | G | H | I | J | K | L | M | N | O |
|---|---|---|---|---|---|---|---|---|---|---|---|---|---|---|
| Gene | Genotype _sample | Mean Cq of technical triplicates | Control group Avg Cq per gene | ΔCq per sample per gene | RQ (rel. quantity) (1+E)^ΔCq | GEOMEAN (norm. factor) [=RQ(actb2)] | Normalized expression per sample | Log2 norm. Expression | Genotype expression | Log2 (J) | Standard deviation SD (I) | SEM | Confidence interval lower limit | Confidence interval upper limit |
| her6 | WT_1 | 24.500 | | -0.177 | 0.884 | 0.857 | 1.032 | 0.045 | | | | | | |
| her6 | WT_2 | 24.640 | 24.323 | -0.317 | 0.801 | 0.903 | 0.888 | -0.172 | 1.004 | 0.005 | 0.154 | 0.089 | 0.947 | 1.057 |
| her6 | WT_3 | 23.830 | | 0.493 | 1.412 | 1.293 | 1.092 | 0.127 | | | | | | |
| her6 | dKO_1 | 21.980 | | 2.343 | 5.145 | 0.958 | 5.367 | 2.424 | | | | | | |
| her6 | dKO_2 | 22.370 | | 1.953 | 3.917 | | | | 4.640 | 2.214 | 0.323 | 0.186 | 1.484 | 2.324 |
| her6 | dKO_3 | 22.090 | | 2.233 | 4.764 | 1.218 | 3.912 | 1.968 | | | | | | |
| her6 | her6KO_1 | 23.160 | | 1.163 | 2.255 | 1.025 | 2.199 | 1.137 | | | | | | |
| her6 | her6KO_2 | 23.890 | | 0.433 | 1.354 | 0.958 | 1.412 | 0.498 | 1.734 | 0.794 | 0.331 | 0.191 | 0.992 | 1.587 |
| her6 | her6KO_3 | 23.870 | | 0.453 | 1.373 | 0.863 | 1.590 | 0.669 | | | | | | |
| her6 | her9KO_1 | 23.410 | | 0.913 | 1.893 | 1.173 | 1.614 | 0.691 | | | | | | |
| her6 | her9KO_2 | 23.620 | | 0.703 | 1.635 | 1.130 | 1.447 | 0.533 | 1.327 | 0.408 | 0.430 | 0.248 | 0.790 | 1.696 |
| her6 | her9KO_3 | 24.150 | | 0.173 | 1.129 | 1.227 | 0.920 | -0.120 | | | | | | |
| her9 | WT_1 | 24.300 | | -0.503 | 0.701 | 0.857 | 0.818 | -0.290 | | | | | | |
| her9 | WT_2 | 24.350 | 23.797 | -0.553 | 0.676 | 0.903 | 0.749 | -0.417 | 1.066 | 0.093 | 0.615 | 0.355 | 0.502 | 2.177 |
| her9 | WT_3 | 22.740 | | 1.057 | 2.110 | 1.293 | 1.632 | 0.707 | | | | | | |
| her9 | dKO_1 | 21.210 | | 2.587 | 6.223 | 0.958 | 6.493 | 2.699 | | | | | | |
| her9 | dKO_2 | 21.790 | | 2.007 | 4.130 | | | | 5.730 | 2.519 | 0.273 | 0.158 | 1.559 | 2.164 |
| her9 | dKO_3 | 21.250 | | 2.547 | 6.050 | 1.218 | 4.968 | 2.313 | | | | | | |
| her9 | her6KO_1 | 23.290 | | 0.507 | 1.431 | 1.025 | 1.395 | 0.481 | | | | | | |
| her9 | her6KO_2 | 23.920 | | -0.123 | 0.917 | 0.958 | 0.956 | -0.065 | 1.123 | 0.168 | 0.292 | 0.169 | 0.867 | 1.258 |
| her9 | her6KO_3 | 23.980 | | -0.183 | 0.878 | 0.863 | 1.018 | 0.025 | | | | | | |
| her9 | her9KO_1 | 21.790 | | 2.007 | 4.130 | 1.173 | 3.521 | 1.816 | | | | | | |
| her9 | her9KO_2 | 22.070 | | 1.727 | 3.389 | 1.130 | 2.999 | 1.584 | 2.834 | 1.503 | 0.428 | 0.247 | 1.169 | 2.495 |
| her9 | her9KO_3 | 22.540 | | 1.257 | 2.431 | 1.227 | 1.981 | 0.986 | | | | | | |
| actb2 | WT_1 | 16.740 | | -0.207 | 0.857 | | | | | | | | | |
| actb2 | WT_2 | 16.670 | 16.533 | -0.137 | 0.903 | | | | | | | | | |
| actb2 | WT_3 | 16.190 | | 0.343 | 1.293 | | | | | | | | | |
| actb2 | dKO_1 | 16.590 | | -0.057 | 0.958 | | | | | | | | | |
| actb2 | dKO_2 | excluded (19.000) | | | | | | | | | | | | |
| actb2 | dKO_3 | 16.270 | | 0.263 | 1.218 | | | | | | | | | |
| actb2 | her6KO_1 | 16.500 | | 0.033 | 1.025 | | | | | | | | | |
| actb2 | her6KO_2 | 16.590 | | -0.057 | 0.958 | | | | | | | | | |
| actb2 | her6KO_3 | 16.730 | | -0.197 | 0.863 | | | | | | | | | |
| actb2 | her9KO_1 | 16.320 | | 0.213 | 1.173 | | | | | | | | | |
| actb2 | her9KO_2 | 16.370 | | 0.163 | 1.130 | | | | | | | | | |
| actb2 | her9KO_3 | 16.260 | | 0.273 | 1.227 | | | | | | | | | |

| PCR efficiency: (E) | |
|---|---|
| her6 | 1.011702888 |
| her9 | 1.027519485 |
| actb2 | 1.113346713 |

**Table 5B: Normalized Expression and Error Bars (SEM)**

| EXPRESSION | | WT | dKO | her6 sKO | her9 sKO |
|---|---|---|---|---|---|
| her6 | | 1.004 | 4.640 | 1.734 | 1.327 |
| her9 | | 1.066 | 5.730 | 1.123 | 2.834 |
| ERROR BARS + (absolute value) | | WT | dKO | her6 sKO | her9 sKO |
| her6 | | 1.068 | 5.279 | 1.980 | 1.576 |
| her9 | | 1.364 | 6.392 | 1.262 | 3.363 |
| ERROR BARS - (absolute value) | | WT | dKO | her6 sKO | her9 sKO |
| her6 | | 0.944 | 4.078 | 1.519 | 1.117 |
| her9 | | 0.834 | 5.137 | 0.999 | 2.388 |
| ERROR BARS + (error bar size) | | WT | dKO | her6 sKO | her9 sKO |
| her6 | | 0.064 | 0.640 | 0.245 | 0.249 |
| her9 | | 0.298 | 0.662 | 0.139 | 0.529 |
| ERROR BARS - (error bar size) | | WT | dKO | her6 sKO | her9 sKO |
| her6 | | 0.060 | 0.562 | 0.215 | 0.210 |
| her9 | | 0.233 | 0.593 | 0.124 | 0.446 |

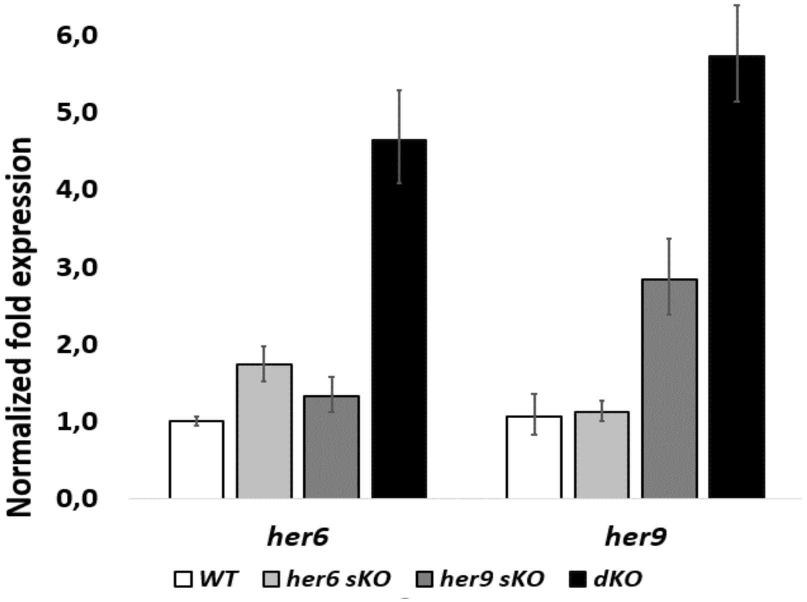

**Table 5C: Statistical Significance of Differences between Genotypes**

| her6 relative expression values | | | | |
|---|---|---|---|---|
| | p-values for comparisons | | | |
| Genotypes | WT | her6 sKO | her9 sKO | dKO |
| WT | | | | |
| her6 sKO | 0.041 | | | |
| her9 sKO | 0.212 | 0.268 | | |
| dKO | 0.007 | 0.019 | 0.012 | |

| her9 relative expression values | | | | |
|---|---|---|---|---|
| | p-values for comparisons | | | |
| Genotypes | WT | her6 sKO | her9 sKO | dKO |
| WT | | | | |
| her6 sKO | 0.866 | | | |
| her9 sKO | 0.030 | 0.022 | | |
| dKO | 0.006 | 0.004 | 0.038 | |

$p < 0.05$

**Supplementary Table 5: Data Analysis Table of *her6* and *her9* qPCR (*her6* and *her9* expression in *her6* and *her9* mutants)**
**Supplementary Table 5A** shows the qPCR Primary Data. *her6* and *her9* expression were measured in wild type and mutant embryos, respectively, each in three biological replicates, which were each analyzed in three technical replicates (numbers are means of technical replicates, data for technical replicates are not shown). *actb2* was used as reference gene to normalize expression. The mean quantitative cycle (mean Cq) of the technical replicates for each sample/target combination was calculated in column C. The average Cq value for the wild type embryos (control group) is shown in column D. The relative difference (ΔCq) between the average Cq for the control group (column D) and the mean Cq (column C) per individual sample within each target is given in column E. The relative quantities were calculated from the ΔCq (i.e., 2ΔCq) and the reaction efficiency from the standard curve (column F). For each wild type/mutant combination, a normalization factor (column G) is calculated from the geometric mean of the associated reference gene relative quantities. The relative quantity (column F) is divided by the normalization factor (column G) to obtain the relative normalized expression per sample (column H). log transformation of column H results in column I. Column J and K are the result of calculating the geometric mean for the samples within each genotype. Standard deviation (SD), standard error of the mean (SEM) and confidence intervals are calculated from the log transformed normalized expression (columns L–O). The small table below the main table shows the PCR efficiencies that were determined and used for calculating the relative quantities. **Supplementary Table 5B** shows the normalized expression and error bars in numbers. The bars represent data from the genotype expression (column J) and the error bars are calculated by 2^(column K +/- column M). **Supplementary Table 5C** shows the tests for statistical significance (3 samples for each genotype / 2 degrees of freedom). Significant differences with p< 0.05 are marked in orange. The analysis was performed based on Taylor, S. C., Nadeau, K., Abbasi, M., Lachance, C., Nguyen, M., & Fenrich, J. (2019). The ultimate qPCR experiment: producing publication quality, reproducible data the first time. *Trends in Biotechnology*, *37*(7), 761-774.
